# *Pseudomonas aeruginosa lasR* is a keystone gene in polymicrobial cultures

**DOI:** 10.1101/2025.10.06.680631

**Authors:** Eva Bernadette Benyei, Rahan Rudland Nazeer, Jemima E.V. Swain, Pok-Man Ho, Anastasios Galanis, Martin Welch

**Affiliations:** University of Cambridge, Department of Biochemistry, Tennis Court Road, Cambridge CB2 1QW, United Kingdom

**Keywords:** *Pseudomonas aeruginosa*, polymicrobial, cystic fibrosis, quorum sensing, antibiotic action, Type VI Secretion

## Abstract

The airways of people with cystic fibrosis (CF) are often co-infected by *Pseudmonas aeruginosa* and a variety of other co-habiting microbes; the infections are polymicrobial. *P. aeruginosa* isolates from the CF airways are also known to commonly acquire mutations in the quorum sensing regulator, *lasR*. The appearance of these *lasR* mutants is associated with a worsening clinical prognosis. In this work, we show that loss of *lasR* function has a significant impact on the stability of inter-species interactions in a polymicrobial ecosystem, and in particular on the dynamics of a common CF-associated fungus, *Candida albicans*. Titres of *C. albicans* were stable in the presence of wild type *P. aeruginosa* and *Staphylococcus aureus*. However, when wild type *P. aeruginosa* was replaced by a Δ*lasR* mutant, *C. albicans* titres progressively declined over time. This instability could be reversed by ectopic expression of a Type VI Secretion System effector cluster (*tsi*) in the Δ*lasR* mutant. We also noted that challenge of the polymicrobial cultures with a clinically-relevant combination of antibiotics (colistin and ciprofloxacin) led to a hyphal bloom of the fungus. This bloom was abolished in the Δ*lasR* mutant, but again, was restored by ectopic expression of the *tsi* cluster. Finally, we show that whereas wild type *P. aeruginosa* is relatively agnostic to the presence of other microbes, the Δ*lasR* mutant is not and undergoes substantial transcriptional reprogramming. Our data indicate that *lasR* has a large and previously unrecognised impact on inter-species interactions. We therefore propose that *lasR* is an ecological keystone gene.

## Introduction

Cystic fibrosis (CF) is the most common lethal inherited disease in caucasian populations. The disease is caused by mutations in the cystic fibrosis transmembrane conductance regulator (CFTR)^1^. The CFTR is a chloride/bicarbonate transporter whose normal function is to facilitate electrolyte and fluid transport across a range of epithelial tissues, including the airways, gastrointestinal, and reproductive tracts. In the case of airway epithelia, this ensures that the surrounding mucus layer remains hydrated and fluid. However, when the CFTR does not function correctly or fails to localize to the epithelial cell membrane (as in the most common CF-associated mutation, ΔF508^2^), the mucus layer become dehydrated, thereby increasing its viscosity and impairing mucociliary clearance. This accumulation of viscous mucus makes the airways of people with CF susceptible to microbial colonization, leading to unremitting inflammation and consequently, tissue damage and progressive loss of lung function. Perhaps the best known CF-associated pathogen is *Pseudomonas aeruginosa*. Indeed, by the time they reach early adulthood, the airways of many people with CF are chronically infected with *P. aeruginosa*^3^. It is not clear why *P. aeruginosa* displays such a penchant for the CF airways, although its penchant for consuming fatty acids (which are abundant in CF lung secretions) likely plays a role^4^.

*P. aeruginosa* often shares the CF airway niche with a variety of other microbial colonists^5,6,7^. Commonly-encountered co-habiting bacterial species are from the genera *Streptococcus, Staphylococcus, Burkholderia, Stenotrophomonas, Achromobacter, Rothia*, and *Haemophilus*. Unexpectedly for a nominally aerobic organ, other frequently encountered colonists include anaerobes such as *Prevotella, Veillonella*, and *Fusobacterium* spp.^8,9,10,11,12^ and also fungal genera such as *Candida* and *Aspergillus* spp.^13^. The prevalence of fungal species in the CF airways is generally under-appreciated. However, between ca. 34% and 78% of people with CF are reported to be colonised by *Candida* spp., with *Candida albicans* being the most commonly detected^14^. This notwithstanding, a lack of robust experimental models has meant that it is still not clear how these co-habiting species impact on the physiology of *P. aeruginosa*, and *vice versa*. To complicate matters further it is also becoming increasingly clear that even within a given species, the population is often heterogenous, with multiple clonally-derived mutant variants co-habiting the same space. However, some genes are more commonly mutated in CF isolates than others^15,16^. In the case of *P. aeruginosa*, these include genes associated with antibiotic resistance such as *nfxB, mexZ*, and *fusA1*^17^, amino acid auxotrophs^17^,^18^, and also global virulence regulators such as *lasR*^16,19,20,21,22, 23,24^. The appearance of *lasR* mutants is often linked with infection progression and worsening outcomes^25,26^.

LasR is the quorum sensing (QS) “master regulator” ^27,28^. In *P. aeruginosa*, QS controls virulence factor production. Because they do not participate in producing secreted virulence factors, *lasR* mutants are often assumed to be “social cheats” which benefit from the public goods made by cooperating individuals without expending any resources themselves^19^. Because cheats invest fewer resources than their cooperating counterparts, they have a fitness advantage and in principle, should sweep through the population. However, as they do so a tipping point will eventually be met, whereby the servile cooperators can no longer generate enough public goods to sustain overall population growth. This serves to limit the carrying capacity of cheats in the population, so *lasR* mutants rarely account for more than ca. 50% (and typically much lower than this) of the species titre^29^. Furthermore, and given their detrimental impact on intra-species population stability, the appearance of cheats is actively policed, usually through mechanisms involving the production of *lasR*-dependent private goods^30,31^ or toxins^32^. The appearance of cheats and of the associated policing mechanisms is predicted by evolutionary theory, and there is strong experimental support for the notion of cheating in mono-species laboratory co-cultures. However, and as noted above, the CF airways are polymicrobial and there is likely a good deal of interaction (both positive and negative) between different species, as well as within each species. To our knowledge, the impact of *lasR* mutants on the stability of polymicrobial cultures has not yet been investigated.

In this work, we examined the impact of loss of *lasR* function on inter-species interactions in a recently-developed continuous flow polymicrobial model of the CF airways^33^. The model employs an artificial sputum medium that faithfully recapitulates the chemical environment of CF lung secretions, and incorporates three species known to co-habit the CF airways: *P. aeruginosa* (a Gram-negative organism), *Staphylococcus aureus* (a Gram-positive organism) and *C. albicans* (a fungus). The steady-state stability achieved with this setup appears to capture that seen in diseased human airways, allowing us to investigate how perturbations such as the introduction of new strains or antibiotic challenge impact on microbial dynamics. Using this setup, we show that loss of *lasR* function has a significant impact on inter-species dynamics, especially following antibiotic challenge. In ecology, when a species has an impact on the ecosystem that is disproportionate to its own abundance in the population, it is known as a keystone species^34^. By analogy, individual genes that have a disproportionately large impact on the ecosystem have recently been dubbed keystone genes^35^. Our data suggest that *lasR* is a keystone gene in the CF-associated ecosystem, and moreover, that loss of *lasR* function also affects the response of the entire ecosystem to antibiotic challenge.

## Results

### Choice of strains to use in the setup

To examine whether introduction of a *lasR* mutant into a polymicrobial culture impacts inter-species dynamics, we used our previously-described continuous flow polymicrobial setup and artificial sputum medium (ASM)^33^. For those cultures containing “wild type” *P. aeruginosa*, we used the laboratory reference strain (hereafter, PAO1_MW_). This strain was chosen because it harbours a functional *mexT* gene, whereas in many other laboratory PAO1 lineages this global virulence/antibiotic resistance regulator is mutated^36,37^. For the *lasR* mutant, we used PAO-R1 (Δ*lasR*::Tc^R 38^, originally from Barbara Iglewski, University of Rochester). In this strain, the *lasR* ORF is deleted and replaced by a tetracycline resistance cassette, Tc^R 38^. Whole genome sequencing of PAO1_MW_ and PAO-R1 revealed that apart from the Δ*lasR*::Tc^R^ mutation, the strains were identical except for the five SNPs/short indels shown in **Table S1**. In addition to *P. aeruginosa*, the triple species cultures also contained *C. albicans* SC5314 (a clinical isolate obtained from a patient in New York in the 1980s) and *S. aureus* ATCC25923 (a clinical isolate obtained from a patient in Seattle in the 1940s).

### Quorum sensing molecule abundance is low in the continuous flow setup

When PAO1_MW_ and the Δ*lasR*::Tc^R^ mutant were grown (separately) in the continuous flow setup as mono-cultures, after 24 h both strains achieved comparable steady state titres of ca. 10^8^ CFU ml^−1^ (**Figure 1A,B**). Quantitation of QS signaling molecules in the PAO1_MW_ cultures confirmed that their concentrations were very low (presumably due to the continual dilution of the cultures with fresh medium), not exceeding 400 nM for *N*-(3-oxo-dodecanoyl)-L-homoserine lactone (OdDHL), and 100 nM each for *N*-butyryl-L-homoserine lactone (BHL) and for the Pseudomonas quinolone signal (PQS) (**Figure S1**). These concentrations are at the lower end of those normally considered to elicit social behavior in *P. aeruginosa*^33^. Negligible quantities of OdDHL or BHL were detectable in the Δ*lasR*::Tc^R^ mutant cultures. Interestingly, significant PQS production was observed in the Δ*lasR*::Tc^R^ mutant poly-cultures, whereas very little of this signal was detected in mono-cultures of the mutant.

**Figure 1.**
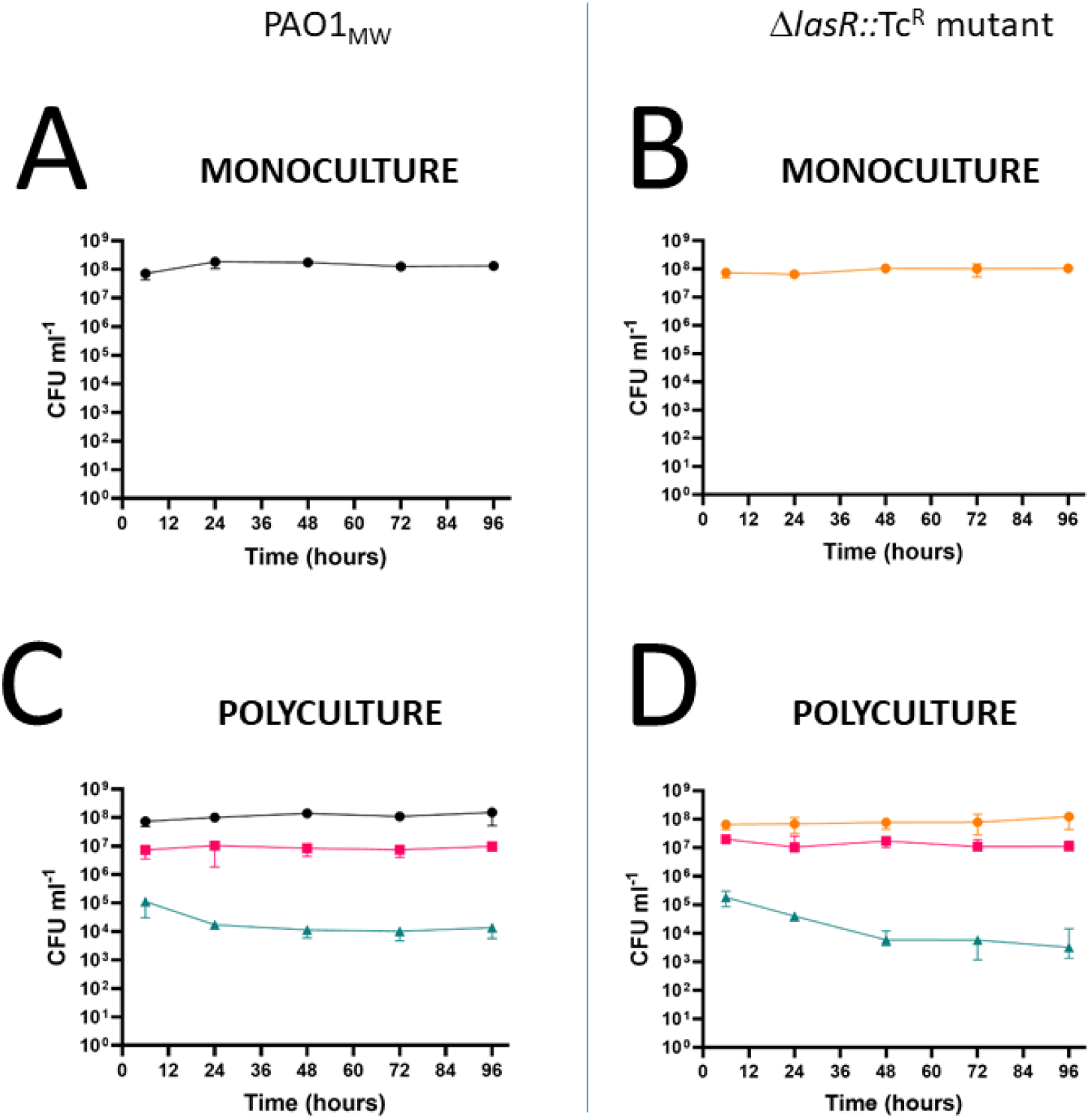
Population dynamics in mono-cultures and poly-cultures containing *Pseudomonas aeruginosa* PAO1_MW_ or the Δ*lasR*::Tc^R^ mutant. Panels A-B show cell titres in monocultures of (**A**) PAO1_MW_ (●), and (**B**) the Δ*lasR*::Tc^R^ mutant 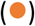. Panels C-D show cell titres of (**C**) PAO1_MW_ 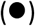, and (**D**) the Δ*lasR*::Tc^R^ mutant 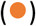grown in co-culture with *S. aureus* 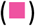 and *C. albicans* 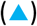. All cultures were inoculated with an initial OD_600_ of the indicated species of 0.05. The growth medium was ASM and the fluid replacement rate (Q) in the bioreactor was 145 μL min^−1^. The data show the median titre with a 95% confidence interval across six independent experiments for **A** and **C**, and four independent experiments for **B** and **D**.

### The *lasR* regulon in the mono-species continuous flow setup comprises just three ORFs

Next, and to establish a transcriptional baseline for each *P. aeruginosa* strain in ASM, we extracted RNA from 48 h mono-cultures of PAO1_MW_ and the Δ*lasR*::Tc^R^ mutant for RNAseq analysis (**Table S2**). With the exception of *lasR* itself, just three transcripts displayed altered (p < 0.05) abundance in the Δ*lasR*::Tc^R^ mutant compared with PAO1_MW_. These were *rsaL* (PA1431), *lasI* (PA1432), and a transcript encoding an uncharacterized c-di-GMP binding protein (PA1433) with probable function similar to that of LapD in *P. fluorescens* Pf0-1^39^. All three transcripts were down-regulated in the absence of *lasR* (**Table S3**). LasR, RsaL, and LasI form part of a positive feedback loop that is thought to be involved in ramping up OdDHL levels during the early pre-quorate period^40^. The absence of LasR-regulated virulence factor transcripts (which would normally be associated with the post-quorate period) and the low overall concentration of OdDHL in the continuous flow cultures is commensurate with this.

### The *lasR* mutant manifests a much larger transcriptional response than the wild type to the presence of other microbes

We next examined the carrying capacities of wild type *P. aeruginosa* (PAO1_MW_) or the Δ*lasR*::Tc^R^ mutant in triple species poly-cultures (**Figure 1C,D**). Both poly-cultures displayed similar overall temporal dynamics, although the carrying capacity of the Δ*lasR*::Tc^R^ mutant appeared to be slightly lower than that of PAO1_MW_, and the *C. albicans* population failed to fully stabilize in cultures containing the Δ*lasR*::Tc^R^ mutant, even after 48 h growth. However, a much larger number of differences were seen when we examined the transcriptome of *P. aeruginosa* in each of the poly-cultures (*cf*. the transcriptome of the same strain measured in mono-culture) (**Figure 2** and **SI Datasets A-C**). To our surprise, PAO1_MW_ appeared to be relatively agnostic to the presence of other species (displaying rather few transcriptional changes in the poly-cultures, **Figure 2A**), whereas the Δ*lasR*::Tc^R^ mutant manifested a much larger number of changes, indicating that it is more sensitive/responsive to the presence of other microbes than the wild type (**Figure 2B**).

**Figure 2.**
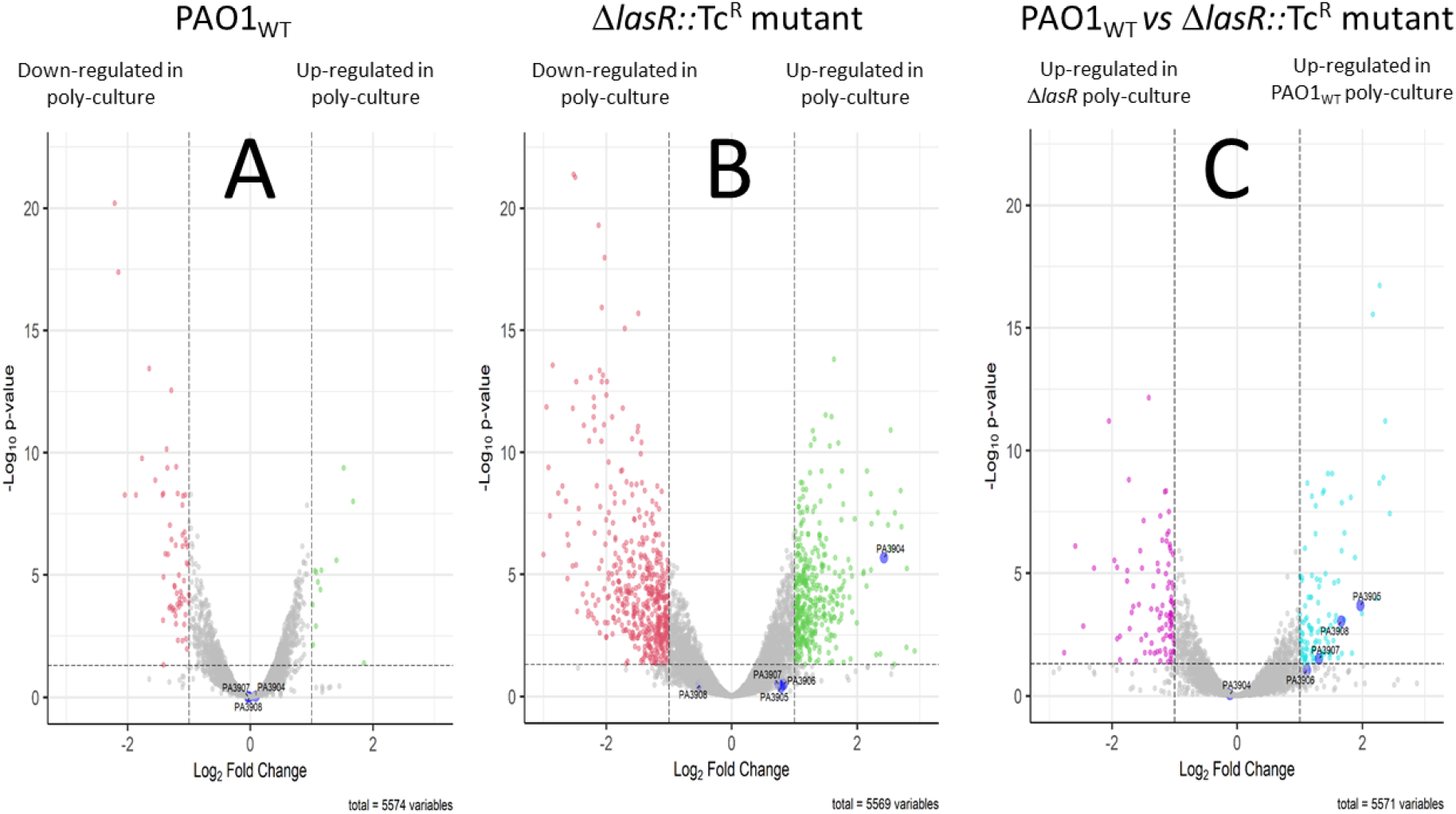
The Δ*lasR*::Tc^R^ mutant displays an altered transcriptional response in the presence of other microbes compared with PAO1_MW_. The figures show enhanced volcano plots illustrating the differentially expressed genes in poly-species *versus* mono-species cultures of (**A**) PAO1_MW_ and (**B**) the Δ*lasR*::Tc^R^ mutant. Panel (**C**) shows the transcripts differentially modulated in poly-cultures of the Δ*lasR*::Tc^R^ mutant compared with poly-cultures of PAO1_MW_. Genes up-regulated in polymicrobial cultures are highlighted in green, whereas those down-regulated are shown in red. Thresholds for significant differential expression are set at log_2_FC > 1.0 and an adjusted p-value < 0.05. ORFs associated with the *tsi* cluster in each volcano plot are highlighted.

To understand better the physiological changes in PAO1_MW_ and in the Δ*lasR*::Tc^R^ mutant in the poly-cultures, we analyzed possible associations between the differentially expressed transcripts using a k means-based STRING analysis. In essence, the AI-based STRING algorithm was instructed to designate the minimal number of putative functional clusters from the data. Next, the algorithm was asked to increment the number of k-means clusters by 1 and the descriptions of clusters were manually compared before and after the increment. This incremental process was then iterated further until all major clusters had explicit functional descriptions, and further increments on the number of k-means clusters neither improved the clarity of the functional descriptions, or generated new clusters. The networks, represented as Voronoi maps, along with a list of the transcripts in each cluster are shown in **Figures S2-S5**. Just 12 transcripts were significantly (p < 0.05) up-regulated in PAO1_MW_ in the poly-cultures, and these were primarily associated with polyamine metabolism, electron transport, and C1-metabolism (**Figure S2**), whereas the largest categories of down-regulated transcripts were related to branched-chain and aromatic amino acid catabolism (16/52 transcripts, cluster 1) and allantoin metabolism (7/52 transcripts, cluster 2) (**Figure S3**). By contrast, the largest categories of significantly up-regulated transcripts in poly-cultures containing the Δ*lasR*::Tc^R^ mutant were associated with translation/protein export (32/117 transcripts, cluster 1) and siderophore production (25 transcripts, cluster 2) (**Figure S4**), whereas the largest clusters of down-regulated transcripts were involved in organic acid and amino acid catabolism (54/243 transcripts, cluster 1) and transcriptional regulation/mixed metabolism (33 transcripts, cluster 2) (**Figure S5**). That transcripts associated with amino acid catabolism (especially branched chain amino acid catabolism) were down-regulated in both PAO1_MW_ and the Δ*lasR*::Tc^R^ mutant was reassuring to see: clearly, both strains are responding to similarly-perceived metabolic cues. However, the much greater modulation (↓) of transcription factors seen in the Δ*lasR*::Tc^R^ mutant poly-cultures (*cf*. the cultures containing PAO1_MW_) also indicates a general physiological “re-wiring” in this mutant in response to the presence of other species.

### A role for the Type VI Secretion machinery

Interestingly, and consistent with our earlier findings that cultures in the continuous flow setup are maintained in a “pre-quorate” state, we observed very few modulated transcripts encoding known LasR-regulated virulence factors. However, an exception to this was a set of ORFs (PA3904-PA3908; hereafter, the *tsi* operon) encoding a Type VI Secretion System (T6SS) effector cluster^41^. Expression of the *tsi* cluster is known to be LasR-regulated, and the *tsi* operon is preceded by a predicted LasR binding motif (*las* box)^42^. Interestingly, the *tsi* cluster is unusual in that, unlike many other QS-regulated genes, it is induced by LasR remarkably early on during the exponential phase of growth. Furthermore, we note that the *tsi* cluster forms part of the “core” QS regulon that is conserved across many *P. aeruginosa* isolates^43^. The possible importance of the *tsi* cluster in inter-species interactions became obvious when we compared the transcriptome of PAO1_MW_ and the Δ*lasR*::Tc^R^ mutant in poly-culture (**Figure 2C**). Here, several of the *tsi*-associated ORFs, including the effector (*tseT*) and immunity (*tsiT*) proteins were up-regulated in PAO1_MW_. Given the function of T6S systems in inter-species (and possibly, also inter-kingdom) conflict, this made us wonder whether restoration of *tsi* operon expression in the Δ*lasR*::Tc^R^ mutant might affect inter-species dynamics in the co-cultures.

To induce expression of the *tsi* cluster independent of LasR in the Δ*lasR*::Tc^R^ background, we cloned the *tsi* operon into mini-Tn7^44^, so that expression of the encoded ORFs come under the control of an IPTG-inducible *lac* promoter (with *lacI* is encoded elsewhere on the mini-Tn7 plasmid). The resulting construct (or as a control, the empty mini-Tn7 vector) was then integrated into the neutral *att* site on the Δ*lasR*::Tc^R^ mutant chromosome. The population dynamics of triple species co-cultures containing one or the other of these two *P. aeruginosa* Δ*lasR*::Tc^R^ variants were monitored over 96 h, with IPTG added at the 48 h timepoint (**Figure 3**). As noted above, in the absence of *lasR*, the *C. albicans* titres appeared somewhat unstable and followed a gradual decline over the 48 h period before addition of IPTG. However, after the addition of IPTG, *C. albicans* titres in the poly-cultures containing the merodiploid strain expressing the *tsi* cluster from the P_lac_ promoter recovered stability, whereas *C. albicans* titres in the poly-cultures containing the empty mini-Tn7 vector continued to decline. Somewhat counter-intuitively, this observation suggests that LasR-dependent expression of the *tsi* effectors leads to increased stability of *C. albicans* in the population.

**Figure 3.**
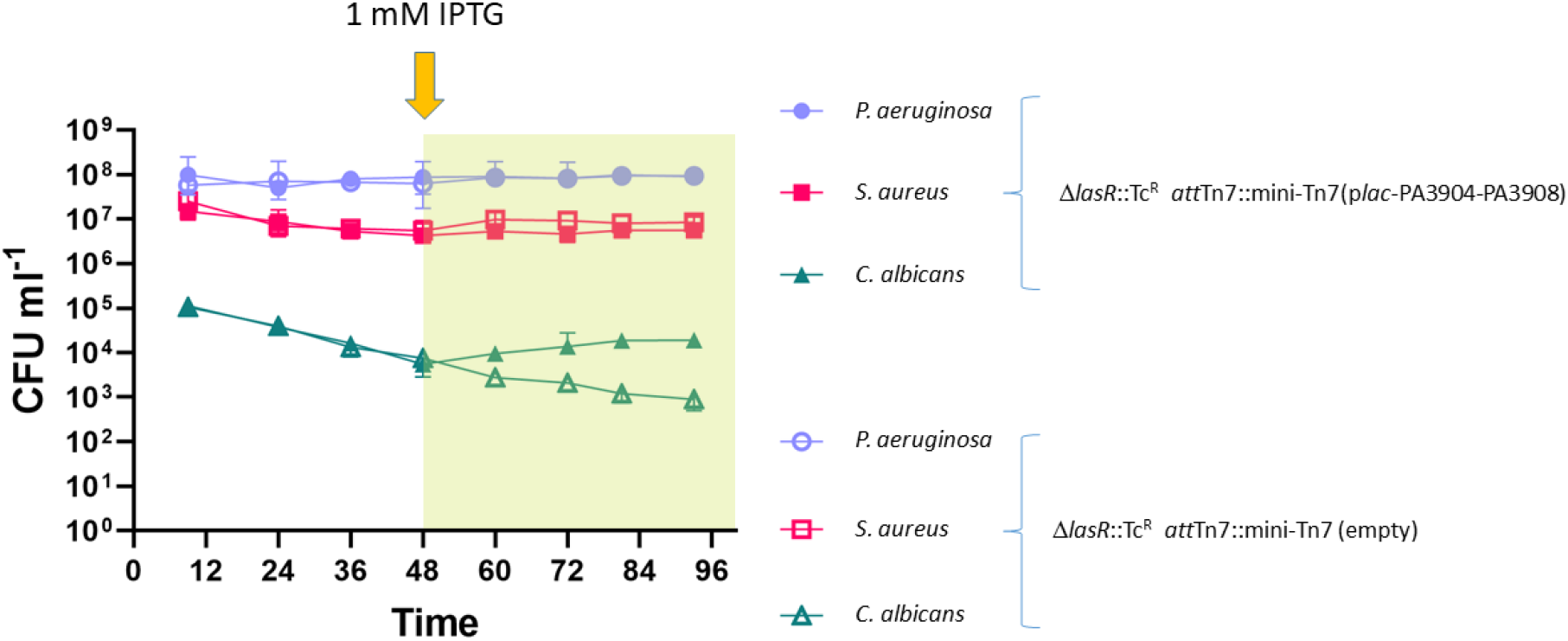
Expression of the *tsi* cluster under IPTG control arrests the decline in *C. albicans* titres in poly-cultures containing the Δ*lasR*::Tc^R^ mutant. Viable cell counts (CFU mL^−1^) of the indicated species in poly-cultures containing either the Δ*lasR*::Tc^R^ *att*Tn7::mini-Tn7(p*lac*-PA3904-PA3908) mutant (filled symbols) or the Δ*lasR*::Tc^R^ *att*Tn7::mini-Tn7 (empty) mutant (open symbols). *P. aeruginosa* (●), *S. aureus* (▪), and *C. albicans* (▴) are shown. Data represent the median with 95% confidence intervals based on three independent experiments. The growth medium was supplemented with IPTG (final concentration 1 mM) from 48 h onwards. The growth medium was ASM and the fluid replacement rate (Q) in the bioreactor was 145 μL min^−1^.

### *LasR* mutant titres are constrained in the presence of a *lasR*^+^ progenitor

Although *lasR* mutants frequently arise in the airways of people with CF, they almost always coexist alongside their *lasR*^+^ counterparts^45^. Therefore, we next examined the carrying capacity and dynamics of mono- and poly-species cultures containing <u>both</u> the Δ*lasR*::Tc^R^ mutant and PAO1_MW_. Intriguingly, we noticed that in the poly-species cultures containing both *P. aeruginosa* strains, *C. albicans* titres were once again stable (**Figure 4B**). This presumably reflects sufficient expression of the *tsi* cluster by the PAO1_MW_ present. However, in both the mono- and poly-species cultures, titres of the Δ*lasR*::Tc^R^ mutant remained at around 1/10^th^ of those of PAO1_MW_ (**Figure 4A,B**). These dynamics are unlikely to reflect the constraints associated with “QS cheats”, since evolutionary theory predicts that the latter should primarily exhibit a fitness advantage in the presence of QS-dependent “public goods” produced by cooperators^19^. By contrast, our data indicate that the cultures are “pre-quorate” with very little LasR-dependent gene expression (**Table S3**) and very low levels of OdDHL accumulation (**Figure S1**). Given our earlier results (**Figure 3**), we also tested whether the absence of *tsi* cluster expression might contribute towards constraining the Δ*lasR*::Tc^R^ mutant titres. It did not: IPTG-dependent induction of *tsi* operon expression in the Δ*lasR*::Tc^R^ mutant background did not change relative competitiveness in mono-cultures containing PAO1_MW_ (**Figure S6**).

**Figure 4.**
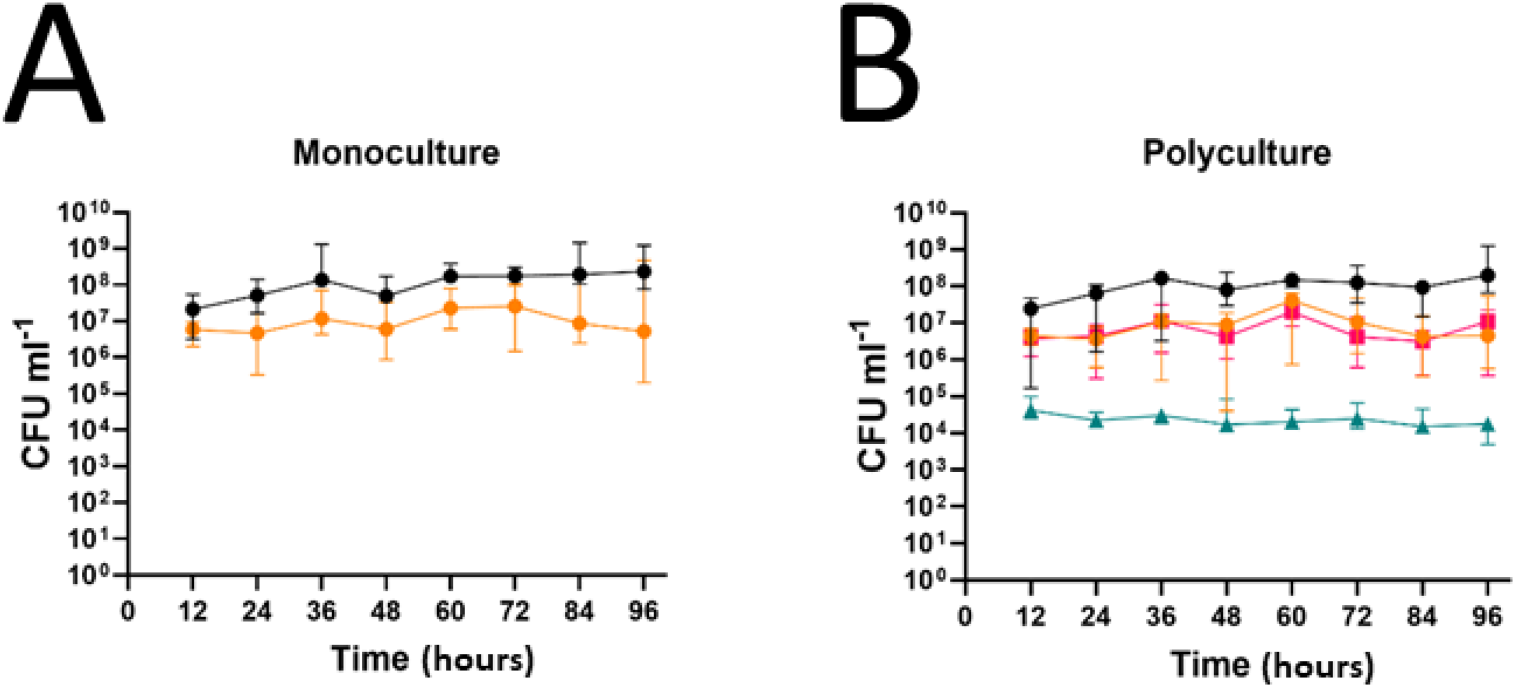
Population dynamics in co-cultures containing both PAO1_MW_ and the Δ*lasR*::Tc^R^ mutant. (**A**) Mono-species co-culture of PAO1_MW_ (●) and the Δ*lasR*::Tc^R^ mutant 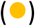, (**B**) Poly-species co-culture of PAO1_MW_ (●), the Δ*lasR*::Tc^R^ mutant 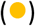, *S. aureus* 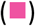 and *C. albicans* 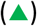. ASM was continuously supplied at a flow rate of 145 μL min^−1^, and all species/strains were initially introduced (at t = 0 h) at an OD_600_ of 0.05. Data are presented as the median with 95% confidence intervals, based on six independent experiments.

### Poly-cultures containing the Δ*lasR*::Tc^R^ mutant display an aberrant response to antibiotic challenge

We have previously shown that growth of *P. aeruginosa* in poly-cultures confers a high degree of protection against colistin action^46^. In the current work, we confirmed that colistin has a much larger impact on PAO1_MW_ in mono-culture (**Figure S7A**) compared with PAO1_MW_ in poly-culture (**Figure 5A**), and further, observe a similar outcome when PAO1_MW_-containing mono- and poly-cultures are challenged with 5 × MIC_P. aeruginosa_ ciprofloxacin (**Figure S7B** and **Figure 5B**). [Note here that, unlike colistin, which has no measurable impact on growth of either *S. aureus* or *C. albicans* (**Table S4**), ciprofloxacin is anti-Staphylococcal. However, given the 16-fold MIC differential of this drug for *P. aeruginosa vs. S. aureus*, at the concentration used in these assays (5 μg mL^−1^) ciprofloxacin should have a disproportionate impact on *P. aeruginosa*.] When colistin and ciprofloxacin were supplied together as a combination (as they often are in the clinic^47^), titres of PAO1_MW_ declined more sharply, as expected (**Figure 5C**). However, and to our surprise – given that these antibiotics should have no effect on the fungus - this was accompanied by a substantial increase (nearly 100-fold) in titres of *C. albicans*. When the same antibiotic combination was added to a continuous flow mono-culture of *C. albicans* there was no change in cell titres (**Figure 5D**).

**Figure 5.**
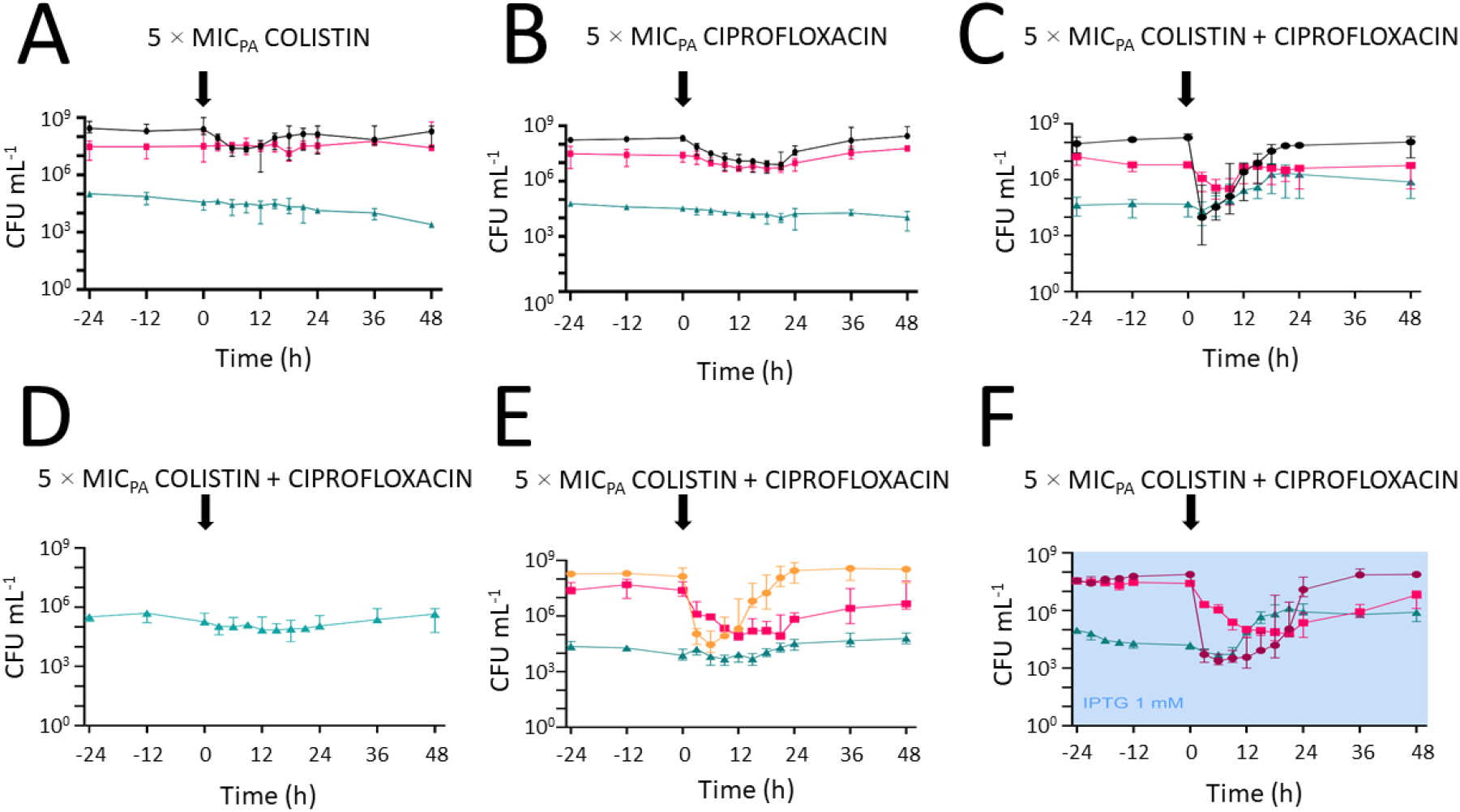
Population dynamics in poly-cultures containing PAO1_MW_ or the Δ*lasR*::Tc^R^ mutant following antibiotic challenge. Steady-state poly-species cultures containing *S. aureus* 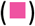, *C. albicans* 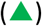, and either PAO1_MW_ (●, (**A**)-(**C**)), Δ*lasR*::Tc^R^ mutant (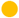, (**E**)), or the Δ*lasR*::Tc^R^ *att*Tn7::mini-Tn7(p*lac*-PA3904-PA3908) mutant (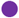, (**F**)) were grown for 48 h and then challenged with a pulse of 5 × MIC_P. aeruginosa_ of the indicated antibiotics. Note that for clarity, we are only showing here the population dynamics in the 24 h period prior to antibiotic addition. The vertical arrow indicates the point at which the antibiotic pulse was introduced into each culture. The culture in (**F**) was supplemented with 1 mM IPTG (indicated by the mauve shading 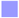) from the start. ASM was continuously supplied at a flow rate of 145 μL min^−1^, and all species/strains were initially introduced (at t = 0 h) at an OD_600_ of 0.05. Data are presented as the median with 95% confidence intervals, based on four (**A,B**), three (**D**), and six (**C,E,F**) independent replicates. Note that in the absence of *tsi* cluster expression by the Δ*lasR*::Tc^R^ mutant (**E**)), *C. albicans* failed to manifest the antibiotic-induced “bloom” seen in the cultures where *tsi* cluster expression was “on” (panels (**C**) and (**F**)).

Microscopic examination of the poly-species cultures before- and after-antibiotic challenge revealed a possible contributor to their elevated antibiotic resistance phenotype. Inspection of steady state cultures prior to antibiotic addition revealed the presence of free-floating cell aggregates (**Figure S8**). Such aggregates have been previously ascribed to a “biofilm-like” lifestyle, with lower antibiotic sensitivity reflecting the impaired access of these agents to the aggregate interior^48^. However, we also noted that whereas all of the antibiotic treatments further enhanced aggregate formation (as might be expected) the combination of colistin and ciprofloxacin in particular was accompanied by the additional appearance of fungal hyphae, many of which appeared to nucleate the aggregative phenotype of the bacterial cells (**Figure S9**). Taken together, these data indicate that antibiotics can have unanticipated impacts on the ecology of microbes that are, in themselves, unaffected by the treatment.

Given our previous observations that loss of *lasR* function can impact on *C. albicans* stability in poly-cultures (**Figure 1D** and **Figure 3**), we next investigated the impact of antibiotics on poly-cultures containing the Δ*lasR*::Tc^R^ mutant. The Δ*lasR*::Tc^R^ mutant had the same measured MIC for colistin and ciprofloxacin as PAO1_MW_ (**Table S4**). Addition of 5 × MIC_P. aeruginosa_ [colistin + ciprofloxacin] to cultures containing the Δ*lasR*::Tc^R^ mutant elicited a similar decline in *P. aeruginosa* cell titres (**Figure 5E**) as we saw in the poly-cultures containing PAO1_MW_ (**Figure 5C**). However, this was also accompanied by a larger than expected decline in *S. aureus* titres, and perhaps more importantly, no hyphal “blooming” of *C. albicans*. In the light of our earlier findings showing that in the Δ*lasR*::Tc^R^ background, the absence of *tsi* cluster expression appears to negatively impact on *C. albicans* population dynamics, we next examined whether heterologous expression of the *tsi* ORFs would restore the antibiotic combination-driven *C. albicans* bloom. It did (**Figure 5F**). Taken together, these data indicate that *C. albicans* titres are indeed destabilized in the absence of *lasR* function, and that this instability is linked with the loss of *tsi* cluster expression. We also note that irrespective of the *tsi* cluster expression status, in the absence of *lasR* function *S. aureus* titres remained perturbed (*cf*. the cultures containing PAO1_MW_) by the combination antibiotic treatment (**Figure 5E,F**). This indicates *lasR* affects inter-species interactions *via tsi*-dependent and *tsi*-independent mechanisms. The greater impact of antibiotics on *S. aureus* grown in co-culture with the Δ*lasR*::Tc^R^ mutant may be related to the QS-dependent formation of small colony variants (SCVs). These slower-growing variants of *S. aureus* are inherently more resistant to antibiotic action than their normo-colony counterparts. We noted SCV formation following co-culture with PAO1_MW_, but normo-colony formation following co-culture with the Δ*lasR*::Tc^R^ mutant.

### Loss of *mexT* function does not affect polymicrobial community dynamics

At this juncture, we wondered whether other virulence-associated global regulators might also impact on polymicrobial dynamics. One such regulator, which is also often found mutated in the airways of people with CF, is *mexT*. MexT activates expression of the *mexEF*-*oprN* multi-drug efflux pump and has recently been shown to be pleiotropic regulator of virulence in *P. aeruginosa*^49^. In this regard, serendipity offered us the opportunity to investigate this further, since the PAO1 variant (hereafter, PAO1_I129F_) that we initially used in this work encoded a SNP in *mexT* leading to an Ile129→Phe substitution. [Note: this is not the PAO1_MW_ strain used in the experiments above, which contains a wild-type *mexT* ORF.] Given that a similar mutation in the adjacent residue (Val130→Phe) was recently shown to give rise to loss-of-function in MexT^36^, it seemed likely that the Ile129→Phe substitution would also inactivate the gene. Nevertheless, and unlike the situation seen in the Δ*lasR*::Tc^R^ mutant, mono- and poly-cultures containing PAO1_I129F_ displayed essentially the same temporal dynamics as PAO1_MW_ (**Figure 6A,B**), suggesting that loss of *mexT* function contributes little to inter-species dynamics.

**Figure 6.**
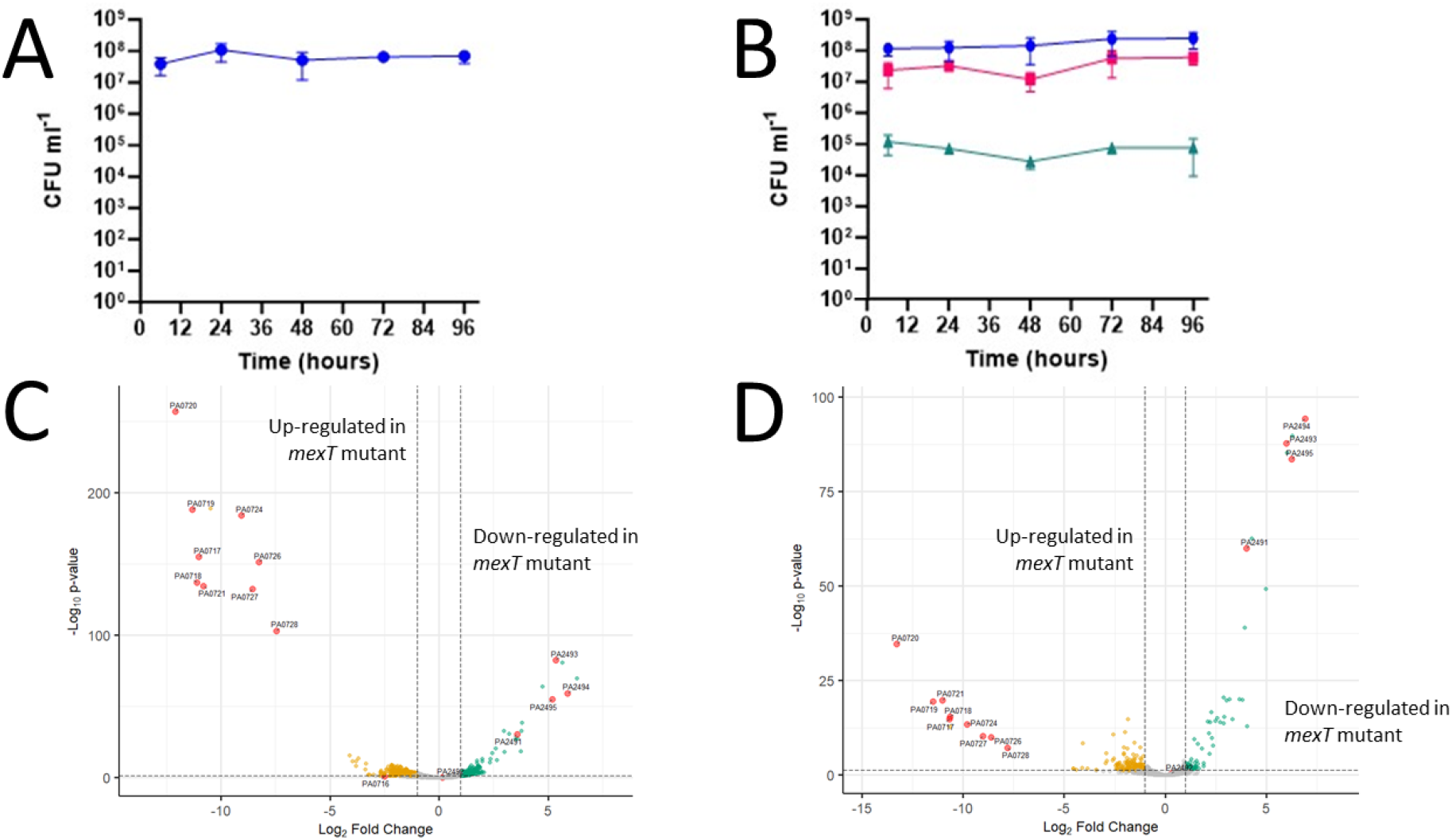
Growth kinetics and transcriptional profile of PAO1_MW_ and the *mexT*^I129F^ mutant in artificial sputum medium. (**A**) Cell titres in a mono-culture of the PAO1-derived *mexT*^I129F^ mutant 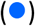, and (**B**) in a poly-culture containing the *mexT*^I129F^ mutant 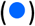, *S. aureus* 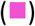 and *C. albicans* 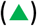. The growth medium was ASM and the fluid replacement rate (Q) in the bioreactor was 145 μL min^−1^. The data show the median titre with a 95% confidence interval. The lower panels show enhanced volcano plots illustrating the differentially abundant transcripts in (**C**) mono-species cultures of the *mexT*^I129F^ mutant *cf*. PAO1_MW_ and (**D**) poly-species cultures containing the *mexT*^I129F^ mutant or PAO1_MW_. The most significantly-modulated ORFs are annotated. PA0716-PA0728 encode a filamentous bacteriophage, Pf1, whereas PA2491-PA2495 encode the *mexS*-*mexT*-*mexEF*-*oprN* cluster of ORFs.

We also examined the transcriptome of PAO1_I129F_ in mono- and poly-culture (**Figure 6C,D, Table S2, SI datasets D** and **E**). These data revealed substantial down-regulation of *mexE* (PA2493), *mexF* (PA2494) and *oprN* (PA2495) in PAO1_I129F_ (consistent with the *mexT*^I129F^ mutation leading to loss-of-function of the gene) as well as many other significant (P <0.05) changes in gene expression (consistent with the pleiotropic impact of this gene, as previously reported^49^). Perhaps most notably, expression of a filamentous bacteriophage, Pf1 (PA0716-PA0728), was strongly up-regulated in the *mexT*^I129F^ mutant. This was also seen in another recent transcriptomic analysis of a *mexT* loss-of-function mutant carried out in buffered LB medium, suggesting that the effect is likely gene-specific rather than driven by growth in a particular environment^50^. The blooming of Pf1 prophage may indicate that the *mexT* mutant cells are experiencing stress.

## Discussion

In this study, we show that loss of *lasR* function can affect inter-species population dynamics in a polymicrobial culture. This is, in part, due to the effect of *lasR* on expression of a Type VI Secretion System (T6SS) effector (*tsi*) cluster. In the absence of *tsi* cluster expression, titres of the fungal pathogen, *C. albicans*, are unstable and progressively decline over time. However, restoration of *tsi* cluster expression in the *lasR* mutant restores *C. albicans* population stability. We further show that the *lasR* mutant manifests a much larger transcriptional response to the presence of other microbes *cf*. the *lasR*^+^ progenitor. In particular, we note that a large number of transcriptional regulators are modulated (↓) in the Δ*lasR*::Tc^R^ mutant following steady-state growth alongside other microbial species, suggesting a radical physiological “rewiring”. Most of these modulated transcriptional regulators have not been characterized, although given that most such regulators are repressors, their down regulation likely leads to de-repression of many genes. We also show that in the absence of the wild type, poly-species cultures containing the *lasR* mutant display an aberrant response to antibiotic challenge. This antibiotic-driven response – associated in the PAO1_MW_ progenitor with the blooming of *C. albicans* hyphae – is restored in cultures of the *lasR* mutant that are forced to express the *tsi* cluster ectopically. By contrast, loss-of-function of another commonly-encountered pleiotropic regulator, *mexT*, had little impact on inter-species dynamics. Taken together, our data indicate that loss of *lasR* function has a disproportionate impact on the ecology of the system; that *lasR* is a “keystone gene” in the CF ecosystem^35^.

LasR-dependent expression of the *tsi* cluster appears to play a particularly important role in controlling *C. albicans* population dynamics in the system. We do not know whether this effect on the fungus is direct or indirect; however, previous studies have indicated that *tsi* cluster expression has little impact on *S. aureus*^42^, and our own studies show that restoration of *tsi* cluster expression in the *lasR* mutant has little impact on its competitiveness with the *lasR*^+^ progenitor in mono-species co-cultures (**Figure S6**). The *tsi* cluster effector protein, TseT, is predicted to be a nuclease secreted *via* the H2-T6SS. Expression of the *tsi* cluster has been previously shown to be both *lasR* and Fe^3+^-regulated, with both factors eliciting strong induction^42^. However, our analyses indicate that siderophore biosynthetic gene expression is up-regulated in poly-cultures containing the Δ*lasR*::Tc^R^ mutant (cluster 2 in **Figure S4**) indicating that iron availability is limited, so *tsi* cluster expression is likely very low in these conditions due to absence of both *lasR* and Fe^3+^. Intriguingly, LasR also regulates expression of the H2 T6SS machinery, raising the question of how any ectopically-expressed TseT might exit the producer cell and enter the target cell. Presumably, even in the absence of LasR-induced H2 expression, enough of the secretory machinery is expressed as a consequence of the action of other regulators^51^ to compensate for this. Interestingly, we note that expression of the H2 T6SS genes has been shown to be stimulated in a *mexT* mutant^50^. This is potentially relevant because mutations in *mexT* are a known and common “bypass” mechanism that partially restores virulence in *lasR* mutants^52^, and *mexT* mutants are also common among CF isolates^50^.

Perhaps more intriguing – given its “conventional” role in killing competitor species – is the question of why expression of a T6SS toxin-encoding cluster should *benefit* (as appears to be the case) a co-habiting microbe (*C. albicans*)? We are not the first to propose this possibility^53^, although to our knowledge, this is the first experimental demonstration of benefit (intended or otherwise) to a co-habiting species. Of course, it is also formally possible that TseT elicits an indirect benefit by acting to limit the loading of other effectors into the H2-T6SS. This “molecular congestion” effect has been reported previously, and prevents the loading and firing of other H2 substrates such as PldA, PldB, Tle3, Tle4, and Tle1^41,54^. Notably, these are all lipases capable of targeting both eukaryotic and prokaryotic cells, which could contribute to modulating inter-species dynamics in a polymicrobial environment.

Taken together, our data indicate that the CF airway should be viewed as a tightly inter-meshed ecosystem, where disruption of one node or spar in the network can have consequences on all the others. We also note that the previous emphasis on “which species are present” in 16S- (and more rarely, ITS-based) analyses ignores a very pertinent additional variable: the impact of intra-species diversity on ecological interactions. Our data indicate that these interactions may be far more complex than previously imagined. For example, it seems that the Candidal bloom that accompanies combination antibiotic therapy is likely not simply due to fungal invasion of a recently-vacated bacterial niche; its dependence on LasR-mediated *tsi* cluster expression begs a more nuanced interpretation. In this regard, we note that fungal blooms following antibiotic intervention have been reported in the clinic too^55,56^, so these observations are likely not simply experimental artifacts. Indeed, the current work raises many new questions. Why does expression of the *tsi* cluster promote a bloom in *C. albicans* hyphae? On a mechanistic level, how does antibiotic intervention impact on this? Why are *lasR* mutant titres constrained in the presence of the wild type, even in pre-quorate growth conditions replete with nutrient; conditions where “QS cheats” should have no obvious fitness advantage? And, do any other commonly-mutated CF-associated *P. aeruginosa* genes also have keystone functions? In this regard, we also note that this study has necessarily focused on a well-characterized, but somewhat minimal three-species experimental model, and on the impact of *P. aeruginosa* mutants. Current efforts are aimed at identifying putative keystone genes in other species, and in expanding the experimental model to include additional CF-associated (and CF airway-derived) species/strains.

## Materials and Methods

### Bacterial strains and culture conditions

PAO1_MW_ was a generous gift from Dao Nguyen (McGill University, Ca). PAO1_I129F_ was a spontaneous *mexT* mutant of PAO1 isolated in the MW laboratory. *C. albicans* SC5314 was a generous gift from Andres Floto (Papworth Hospital CF unit). *S. aureus* ATCC25923 was obtained from the ATCC. The quorum sensing molecule biosensors (*P. aeruginosa* Δ*pqsA* P_*pqsA*_::*luxCDABE* (for PQS), *E. coli* JM109 pSB536 (for BHL), and *E. coli* JM109 pSB1075 (for OdDHL)) were maintained and used as previously described^57,58,59^. The Artificial Sputum Medium (ASM) was identical to SCFM2^11,60,61^ except that porcine stomach mucin was used in place of bovine maxillary mucin, as previously described^33^. The continuous flow setup was as previously described, with Q = 145 μL min^−1^ and the initial OD_600_ inoculation for each strain of 0.05, unless otherwise stated.

For CFU enumeration, aliquots were withdrawn from the bioreactors at the indicated times, and ten-fold serial dilutions were prepared in sterile PBS. Aliquots (20 μL volume) of each dilution were spotted onto selective agar plates. Pseudomonas Isolation Agar (PIA, Millipore) was used to select for *P. aeruginosa*, Mannitol Salt Agar (MSA, Oxoid) for *S. aureus*, and BiGGY Agar (Bismuth Glycine Glucose Yeast agar, Millipore) for *C. albicans*. Plates were incubated at 37°C. Colonies of *P. aeruginosa* and *S. aureus* were typically counted after 16-18 h of growth (or 72 h when SCVs were manifestly present). *C. albicans* colonies were counted after 24 h growth. CFU counts were determined as the average of at least two technical replicates for each biological replicate. Unless otherwise stated, all data are reported as the median with 95% confidence interval based on at least three independent biological replicates (N ≥ 3). Statistical significance was defined as p < 0.05 for all analyses. Growth data from continuous-flow cultures were analysed using the Wilcoxon signed-rank test, conducted in either GraphPad Prism (v10.3.1) or R (v4.4.1).

For experiments involving antibiotics, cultures were first allowed to grow for 48 h to establish a steady-state. Fresh antibiotic stock solutions were prepared on the day of the experiment and diluted with ASM to 100 × the final desired concentration. A 1 mL aliquot of the relevant solution was added to each culture to achieve a final concentration of 5 × MIC_PA_ for the corresponding *P. aeruginosa* genetic variant. Following the addition of antibiotics, the flow of fresh media was paused for one hour before being resumed. When IPTG was required, and unless otherwise indicated, it was added to 1 mM final concentration in the bioreactor, and also to the input flow medium (also to 1 mM final concentration).

### Minimum Inhibitory Concentration determination

Minimum inhibitory concentration (MIC) values for colistin and ciprofloxacin were determined using the broth microdilution method described by the Clinical and Laboratory Standards Institute (CLSI^62^). Briefly, serial two-fold dilutions of the antibiotics were prepared in ASM, such that their concentrations (after mixing with the bacterial cultures) ranged from 0 µg mL^−1^ to 256 µg mL^−1^. Aliquots of each dilution were dispensed into microtitre plates. Overnight cultures of the indicated strain/species were washed three times in sterile PBS, and the cultures were inoculated into wells of the microtitre plates to an initial OD_600_ of 0.05. Each well contained a final volume of 150 μL. The plates were sealed with a gas-permeable membrane (4TITUDE) and incubated at 37°C for 16 h with moderate shaking (50 rpm). The MIC was determined as the lowest antibiotic concentration at which no visible bacterial growth was detected.

### Construction of the Δ*lasR*::Tc^R^ *att*Tn7::mini-Tn7(p*lac*-PA3904-PA3908) mutant

For chromosomal integration of the *tsi* cluster, the mini-Tn7 system was employed^44^. The entire operon was inserted into KpnI-digested pJM101 using HiFi assembly. Cloning was confirmed with whole plasmid sequencing (Plasmidsaurus). The integron was then introduced into PAO1_MW_ by electroporation. Successful integrants were selected with gentamicin.

### Whole genome sequencing (WGS) and analyses

WGS was conducted on *P. aeruginosa* PAO1_MW_ and the Δ*lasR*::Tc^R^ mutant to identify the full spectrum of differences between these strains. MicrobesNG (Birmingham, UK) carried out the sequencing, following its established protocols (MicrobesNG *Methods*, 2024). Briefly, an aliquot (100 µL) of overnight culture of each strain was transferred into 50 mL of fresh LB broth and incubated at 37°C with shaking at 100 rpm until mid-log phase. Cells were then harvested and resuspended in DNA/RNA Shield (Zymo Research, USA). Genomic DNA libraries were prepared using the Nextera XT Library Prep Kit (Illumina, San Diego, USA), and sequencing was performed on an Illumina NovaSeq 6000 platform with a 250 bp paired-end protocol. Sequencing reads were adapter-trimmed using Trimmomatic (v0.30) with a sliding window quality cutoff of Q15^63^. *De novo* assembly was completed using SPAdes (v3.7), and the resulting contigs were annotated with Prokka (v1.11)^64,65^. Reads were aligned to the reference genome (GCF_000006765.1) using BWA mem (v0.7.17), and SAMtools (v1.9) was used for processing^66^. Variants were called with VarScan (v2.4.0), using thresholds of 90% for sensitive and 10% for specific allele frequencies. Variant effects were predicted and annotated using SnpEff (v4.3). The Δ*lasR*::Tc^R^ mutant was compared with PAO1_MW_ to identify genetic differences.

### RNAseq analyses

The RNA extraction protocol was tailored for isolation of *P. aeruginosa* transcripts. Briefly, samples (2 mL volume) for bulk RNA sequencing were collected at the 48 h time-point directly from the bioreactor setup. The cells were immediately pelleted by centrifugation at 15,000 × *g* for 5 min at 4°C, and the supernatant was discarded. The cells were then treated with 300 µL of 3 mg mL^−1^ lysozyme (Sigma) and 600 µL RNA Lysis Buffer. The sample was then clarified (15,000 × *g* for 2 min at room temperature) to pellet cellular debris, after which the supernatant was transferred to an RNase-free microfuge tube. DNA depletion and RNA purification was carried out using a Monarch Total RNA Miniprep Kit (NEB) following the manufacturer’s instructions, with an additional DNase treatment to ensure complete removal of any residual gDNA. RNA Priming Buffer and Wash Buffer were used to eliminate contaminants, and RNA was eluted with nuclease-free water. Following RNA elution, the Nucleic Acid Sequencing Facility (Department of Biochemistry) carried out the subsequent sample processing. First, RNA sample quality was assessed using an Agilent 2100 Bioanalyzer to confirm RNA integrity and provide RNA Integrity Number (RIN) values. Ribosomal RNA was depleted using the Ribo-Zero rRNA Removal Kit (Illumina). The library was prepared using the TruSeq Stranded mRNA Library Prep Kit (Illumina). Sequencing was conducted on an Illumina NextSeq 500 platform with a 150-cycle high-output run. Paired-end (75 bp) sequencing yielded approximately 400 million total reads, with an average of 22.5 million reads per sample. The output FASTQ files were transferred back *via* the Illumina BaseSpace Sequence Hub and downloaded for data analysis. Quality control (QC) of the raw reads was performed using FASTQC (v12.0) to assess base quality scores, GC content, and other metrics^67^.

Sequence reads were processed using the Cambridge HPC cluster and a custom pipeline developed for this project. Miniconda3 (Conda v24.5.0) was used to manage the software environment and dependencies. Adapter sequences and low-quality bases were trimmed using Cutadapt (v4.9), and QC was repeated to confirm the integrity of the processed reads^68^. Although the purification protocol was tailored to enrich for *P. aeruginosa* mRNA, the samples inevitably also contained reads from the other microbial species present (*S. aureus* and *C. albicans*) so a ‘virtual’ genome was created by combining the genomes of all three microorganisms. The sequence reads were aligned to this composite genome using Bowtie2 (v2.5.4), generating SAM files^69^. Samtools (v1.20) was then used to filter out reads mapped to the *C. albicans* and *S. aureus* genomes, resulting in BAM files containing only reads mapped to the *P. aeruginosa* genome^70,71^. FeatureCounts from the Subread package (v2.0.6) was used to count reads assigned to coding sequences (CDS) of the *P. aeruginosa* genome^72^. The results were exported as CSV files. For detailed data analysis, R (v4.4.1) was used. DESeq2 package (v1.44.0) was utilised for normalisation and differential expression analysis^73^. The analysis and visualisation pipeline were based on the “Analyzing RNA-seq data with DESeq2” protocol available on the Bioconductor website. Principal Component(s) Analysis (PCA) was used to evaluate sample clustering using the DESeq2 (v1.44.0) and ggplot2 (v3.5.1) packages^73^. EnhancedVolcano was employed to define and visualise differentially expressed genes^74^. The significance thresholds were set at log_2_FC > 1.0 and an adjusted p-value < 0.05. Functional and biological interactions between differentially expressed genes and related proteins were investigated using STRING (v12.0)^75^ and k-means clustering as described in the main text. Briefly, connections and interaction networks were visualised using the Markov Cluster Algorithm (MCL) with the inflation parameter set to 1.2, which allowed for less stringent clustering and the identification of broader interaction groups.

## Contributions

EBB carried out most of the experimental work and prepared the draft manuscript. JEVS and RRN assisted in making strains and mutants, in designing the experiments, and in analyzing the data. PMH carried out computational modelling of the data. MW secured funding for the study and wrote the final paper. All authors contributed towards editing and reviewing the content.

