## Supplementary Figures and Tables for "*Pseudomonas aeruginosa lasR* is a keystone gene in polymicrobial cultures"

***Table S1.*** **Mutations (apart from the Δ*lasR*::Tc$^{R}$ insertion) found in the *Pseudomonas aeruginosa* Δ*lasR*::Tc$^{R}$ mutant compared with PAO1$_{MW}$.**

| Position | Locus | Product | Type | Mutation | Annotation |
|---|---|---|---|---|---|
| 1,215,657 | PA1122 | Putative peptide deformylase | Frameshift mutation | A → AG | --- |
| 4,212,201 | PA3760 | N-acetyl-D-glucosamine phosphotransferase system transporter | Missense mutation | A → G | H636R |
| 4,771,865 | PA4268 | 30S ribosomal protein S12 | Missense mutation | T → C | K88R |
| 5,655,220 | PA5024 | Conserved hypothetical protein | In-frame triplet deletion | C**CGG** → C | GW141W |
| 5,676,046 | PA5040 | Type 4 fimbrial biogenesis outer membrane protein PilQ precursor | Missense mutation | G → T | P605T |

***Table S2.*** **Quality control metrics and RNAseq read counts.** The table shows the QC metrics for each sample submitted for RNAseq analysis, including RIN, GC content, and Base Quality Score (Phred Score), the total number of reads, the percentage of paired reads, the overall alignment rate to a combined 'virtual' genome of *P. aeruginosa* (PAO1$_{MW}$), *C. albicans* (SC5314) and *S. aureus* (ATCC25923) – done to fuilter out reads matching the *C. albicans* or *S. aureus* genomes (using Bowtie2 and Samtool), and percentage of reads mapped to the PAO1 genome. Feature counts were generated from the reads that matched the PAO1$_{MW}$ genome, with an additional quality filter applied (using FeatureCounts).

| Sample | RIN | GC content | Base Quality Scores | Total reads | Paired | Overall alignment rate | Mapped to PAO1 |
|---|---|---|---|---|---|---|---|
| *lasR*_mono_A | 9.9 | 57 | 30+ | 22,508,069 | 100% | 97.63% | 100% |
| *lasR*_mono_B | 10 | 57 | 30+ | 20,945,546 | 100% | 97.90% | 99.99% |
| *lasR*_mono_C | 10 | 57 | 30+ | 18,200,990 | 100% | 97.67% | 100% |
| WT_mono_A | 9.7 | 56 | 30+ | 25,078,076 | 100% | 98.70% | 100% |
| WT_mono_B | 9.8 | 57 | 30+ | 20,452,156 | 100% | 98.51% | 99.99% |
| WT_mono_C | 8.8 | 58 | 30+ | 15,248,241 | 100% | 97.61% | 100% |
| *lasR*_poly_A | 10 | 59 | 30+ | 15,062,703 | 100% | 95.97% | 98.98% |
| *lasR*_poly_B | 9.7 | 59 | 30+ | 17,906,557 | 100% | 87.01% | 99.42% |
| *lasR*_poly_C | 10 | 58 | 30+ | 15,566,407 | 100% | 84.58% | 98.42% |
| WT_poly_A | 9.5 | 57 | 30+ | 21,406,291 | 100% | 98.17% | 99.95% |
| WT_poly_B | 9.6 | 57 | 30+ | 24,394,084 | 100% | 97.97% | 99.89% |
| WT_poly_C | 9.3 | 60 | 30+ | 19,977,580 | 100% | 95.09% | 99.84% |
| *mexT*_mono_A | 9.2 | 58 | 30+ | 21906864 | 100% | 97.01% | 99.93% |
| *mexT*_mono_B | 10 | 58 | 30+ | 23818538 | 100% | 95.77% | 99.98% |
| *mexT*_mono_C | 10 | 57 | 30+ | 21627022 | 100% | 96.34% | 99.93% |
| *mexT*_mono_D | 9.7 | 51 | 28+ | 11693252 | 100% | 64.81% | 66.43% |
| *mexT*_poly_A | 10 | 58 | 30+ | 20315530 | 100% | 94.46% | 99.93% |
| *mexT*_poly_B | 9.8 | 53 | 28+ | 16186784 | 100% | 77.10% | 79.28% |
| *mexT*_poly_C | 9.9 | 54 | 28+ | 32396948 | 100% | 98.25% | 99.98% |
| *mexT*_poly_D | 9.5 | 53 | 30+ | 23011300 | 100% | 96.72% | 99.13% |

***Table S3*. List of transcripts significantly modulated in ∆*lasR*::Tc$^R$ *cf*. PAO1$_{MW}$ during growth of each strain in mono-culture.** All of the indicated transcripts were less abundant in the ∆*lasR*::Tc$^R$ mutant. No significant up-regulated transcripts were observed.

| Gene | Name | Product | log2 FC | Adjusted p-value |
|---|---|---|---|---|
| PA1430 | *lasR* | transcriptional regulator LasR | 5.551577 | 3.26E-45 |
| PA1431 | *rsaL* | NA | 1.767839 | 0.022466 |
| PA1432 | *lasI* | autoinducer synthesis protein LasI | 3.591374 | 6.13E-11 |
| PA1433 | conserved hypothetical protein | NA | 1.696242 | 0.006615 |

***Table S4*. Minimum inhibitory concentration of colistin and ciprofloxacin for the strains/species used in this study.** MICs were measured by broth microdilution in liquid cultures of the indicated strains grown in ASM.

| Species | Strain | Minimum Inhibitory Concentration (µg mL$^{-1}$) | |
|---|---|---|---|
| | | Colistin | Ciprofloxacin |
| *P. aeruginosa* | PAO1$_{MW}$ | 4 | 1 |
| | ∆*lasR*::Tc$^R$ | 4 | 1 |
| | ∆*lasR*::Tc$^R$ *att*Tn7::mini-Tn7(p*lac*-PA3904-PA3908) | 4 | 1 |
| *S. aureus* | ATCC 25923 | >256 | 16 |
| *C. albicans* | SC5314 | >256 | >256 |

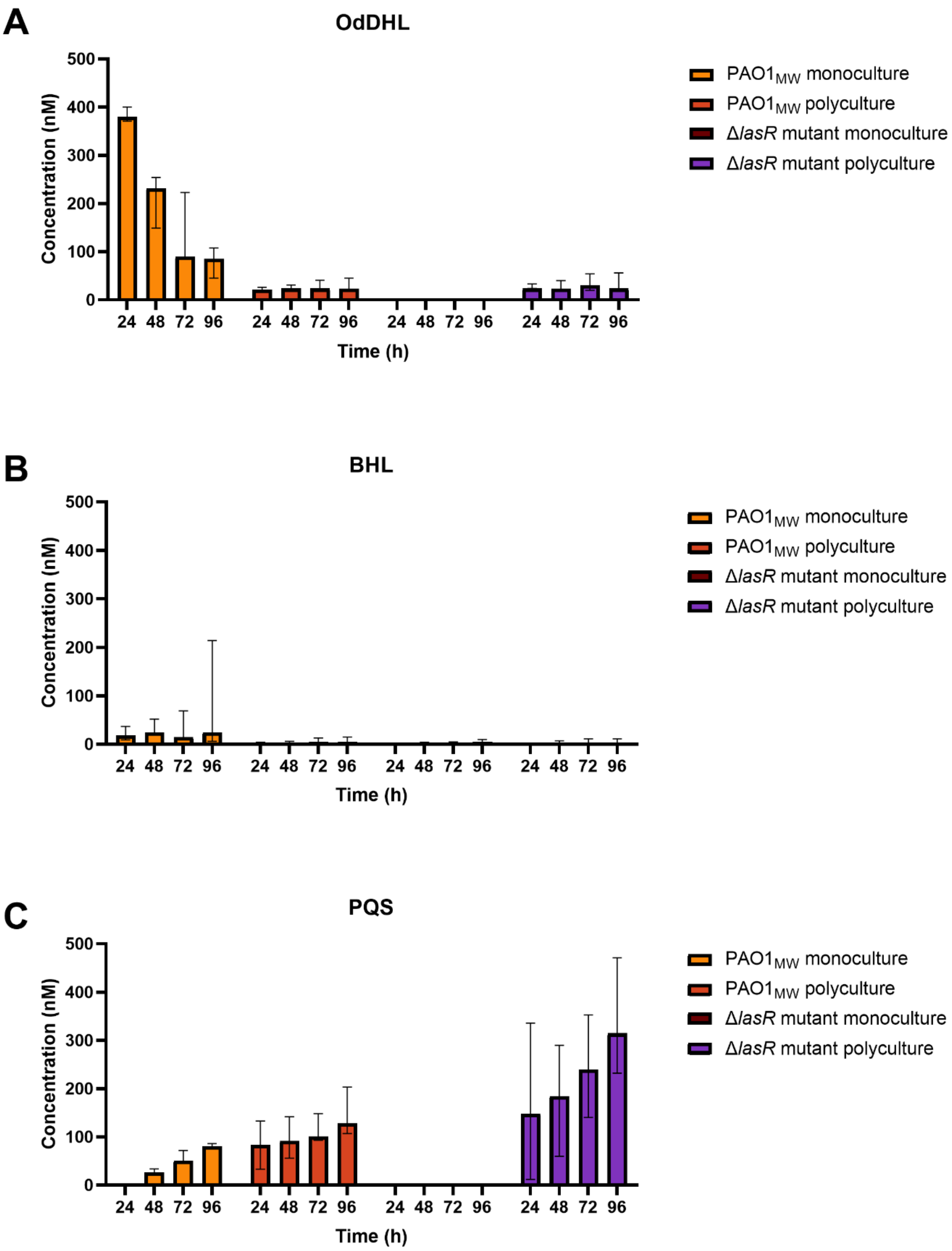


***Figure S1.*** **Quantification of steady-state quorum sensing molecule concentrations in the continuous-flow system.** The concentration (in nM) of the indicated QS molecules was measured in supernatants from the indicated cultures (PAO1$_{MW}$ or the Δ*lasR*::Tc$^{R}$ mutant) after 24, 48, 72, and 96 h incubation. (**A**) *N*-(3-oxododecanoyl)-L-homoserine lactone (OdDHL); (**B**) *N*-butanoyl-L-homoserine lactone (BHL); (**C**) Pseudomonas quinolone signal (PQS). The data show the median ± 95% confidence interval from three independent experiments.

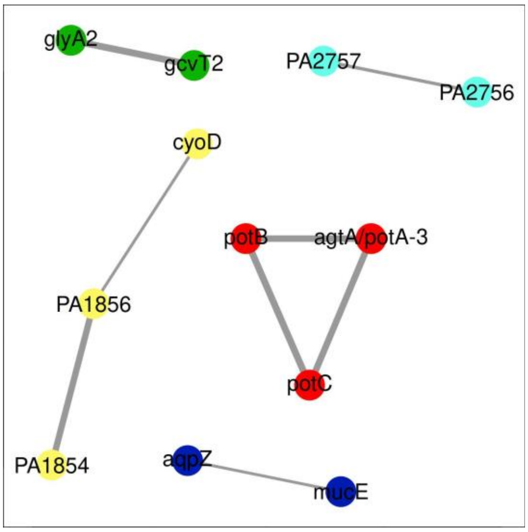


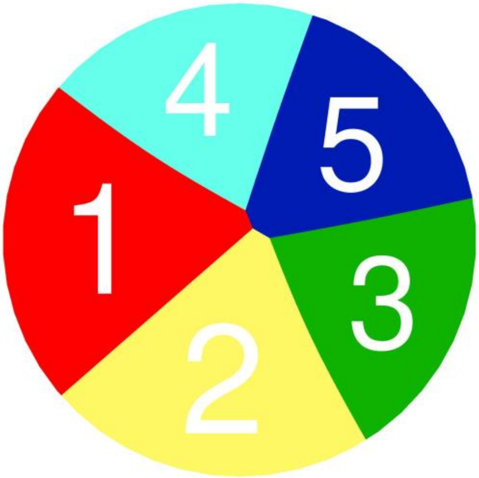


| Cluster | Gene Count | Description | Genes |
|---|---|---|---|
| 1 | 3 | Polyamine transport, and MetI-like superfamily | *potA-3*, potB, *potC* |
| 2 | 3 | Cytochrome complex | PA1854, PA1856, *cyoD* |
| 3 | 2 | One carbon pool by folate; Glycine catabolism | *gcvT2*, *glyA2* |
| 4 | 2 | Mixed, incl. Annealing activity, and Chlorhexidine efflux transporter | PA2756, PA2757 |

| | | |
|---|---|---|
| 5 | 2 | *aqpZ, mucE* |

***Figure S2.*** **Markov clustering of transcripts displaying increased abundance in PAO1$_{MW}$ during growth in poly-culture. The data clusters are represented as STRING maps and as Voronoi tesselations. The colour coding in the STRING maps and Voronoi tesselations is the same, and the identities of the modulated transcripts in each cluster are shown in the colour-coded table below the Figure.**

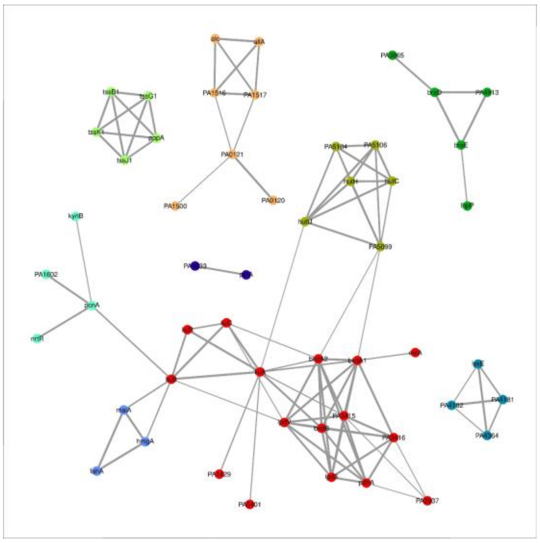

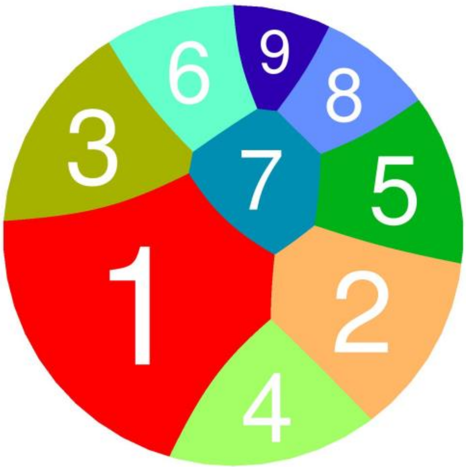


| Cluster | Gene Count | Description | Genes |
|---|---|---|---|
| 1 | 16 | Branched-chain amino acid catabolism; Valine, leucine and isoleucine degradation; Oxidoreductase activity, acting on the aldehyde or oxo group of donors, disulfide as acceptor, and Fatty acid binding | *liuB*, *liuA*, *liuR*, *liuC*, *bkdA2*, *lpdV*, PA3416, *pdhA*, *lpd3*, PA3415, PA2937, *bkdA1*, *bkdB*, *estA*, PA5401, PA1429 |
| 2 | 7 | Allantoin metabolism; Mixed, incl. Purine metabolism, and NodB homology domain | PA0121, PA1517, PA1516, *alc*, *allA*, PA1500, PA0120 |
| 3 | 6 | Histidine catabolism | PA5099, *hutU*, PA5106, *hutC*, PA5104, *hutH* |
| 4 | 5 | Biofilm formation - Pseudomonas aeruginosa, and Tetratricopeptide repeat | PA0079, PA0080, PA0089, PA0083, *pppA* |
| 5 | 5 | D-alanine transport;Amino-acid transport | PA1260, *braE*, *braD*, PA4913, PA3865 |

| | | | |
|---|---|---|---|
| 6 | 4 | Nicotinate and nicotinamide metabolism; Mixed, incl. NAD salvage, and Polyketide cyclase SnoaL-like | *kynB*, PA4918, PA4916, PA1602 |
| 7 | 4 | Mixed, incl. ACT domain , and B3/4 domain | PA4182, PA4364, PA4365, PA4181 |
| 8 | 3 | Tyrosine catabolism; Styrene degradation | *hmgA*, *fahA*, *maiA* |
| 9 | 2 | Transducer; Methyl-accepting chemotaxis-like domains (chemotaxis sensory transducer).; Cache domain, and Chemotaxis protein methyltransferase CheR | PA4633, *pctA* |

***Figure S3.*** **Markov clustering of transcripts displaying decreased abundance in $PAO1_{MW}$ during growth in poly-culture. The data clusters are representated as STRING maps and as Voronoi tesselations. The colour coding in the STRING maps and Voronoi tesselations is the same, and the identities of the modulated transcripts in each cluster are shown in the colour-coded table below the Figure.**

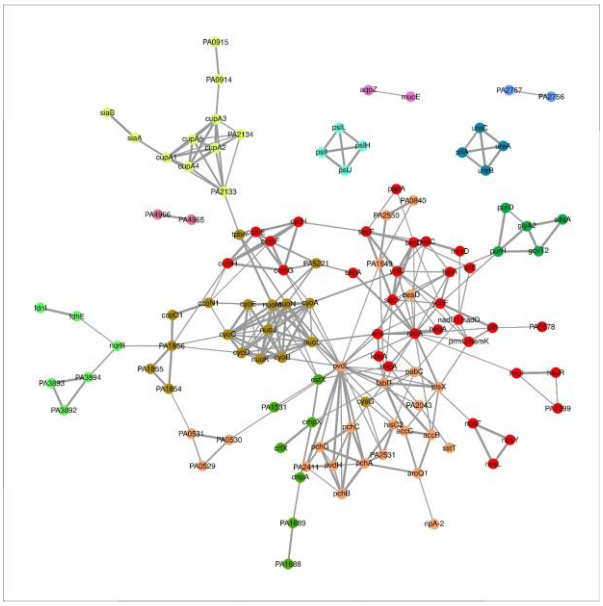

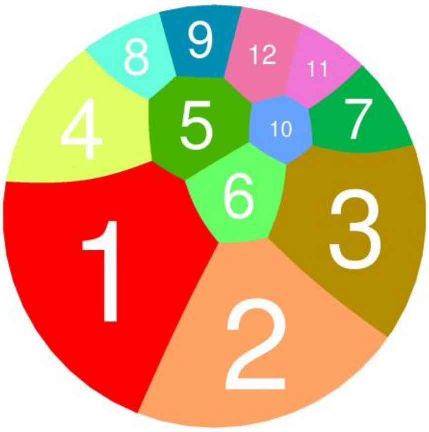


| Cluster | Gene Count | Description | Genes |
|---|---|---|---|
| 1 | 31 | Translation and Protein export | *sspA, rpoA, secY, yidC, typA, ychF, pth, hemK, nadD, mreD, mreC, secF, secD, lptE, pgpA, cycH, ccmH, kdsA, eno, nosF, nosY, nosL, hepA, ccmF, ccmG, ccmE, micA*, PA0578, PA1300, PA1301, PA1299 |
| 2 | 25 | Monocarboxylic acid biosynthesis; Mixed, incl. Siderophore metabolism, and Biosynthesis of siderophore group nonribosomal peptides; Vitamin binding | PA0531, PA0530, *pvdL, pchC, pchB, pchA, aroQ1, accB, accC, plsX, fabD, deaD, hisC2, pabC*, PA2531, PA2043, *sstT*, PA1649, PA2550, PA0840, *rlpA-2, pchG, pvdH*, PA2411, PA0529 |
| 3 | 18 | Oxidative phosphorylation | *nuoN, tpbA, nuoM*, PA5221, PA1856, PA1855, PA1854, *nuoL, nuoK, nuoJ, cyoD, cyoE, cyoC, ccoN1, ccoO1, cyoA, cyoB, cysG* |
| 4 | 11 | Cellular response to oxygen levels | PA2133, PA2134, *cupA5, cupA4, cupA3, cupA2, cupA1*, PA0172, PA0171, PA0914, PA0915 |

| | | | |
|---|---|---|---|
| 5 | 7 | Mixed, incl. Lipopolysaccharide kinase (Kdo/WaaP) family, and Lipopolysaccharide core region metabolism | *cmpX*, PA5002, PA1689, PA1688, *crfX, estX*, PA1531 |
| 6 | 6 | Para-hydroxybenzoic acid efflux pump subunit AaeB/fusaric acid resistance protein, and Membrane fusion protein, biotin-lipoyl like domain | *nqrB*, PA3894, PA3893, PA3892, *fdhE, fdnI* |
| 7 | 5 | One carbon pool by folate; Nucleoside monophosphate metabolism, and Glycine, serine and threonine metabolism; Hydroxymethyl-, formyl- and related transferase activity | *purH, purD, glyA2, sdaA, gcvT2* |
| 8 | 4 | Extracellular polysaccharide biosynthesis | *pslI, pslJ, pslL, pslH* |
| 9 | 4 | Urea catabolism | *pitA, ureB, ureC, ureA* |
| 10 | 2 | Mixed, incl. Annealing activity, and Chlorhexidine efflux transporter | PA2757, PA2756 |
| 11 | 2 | | PA4033, *aqpZ* |
| 12 | 2 | DNA topoisomerase IV complex, and Retropepsin-like domain, bacterial | PA4966, PA4965 |

***Figure S4.*** **Markov clustering of transcripts displaying increased abundance in the Δ*lasR*::Tc$^R$ mutant during growth in poly-culture. The data clusters are representated as STRING maps and as Voronoi tesselations. The colour coding in the STRING maps and Voronoi tesselations is the same, and the identities of the modulated transcripts in each cluster are shown in the colour-coded table below the Figure.**

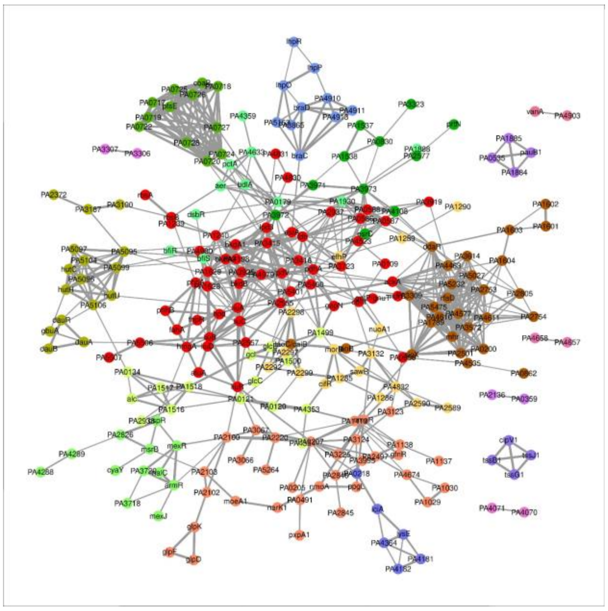

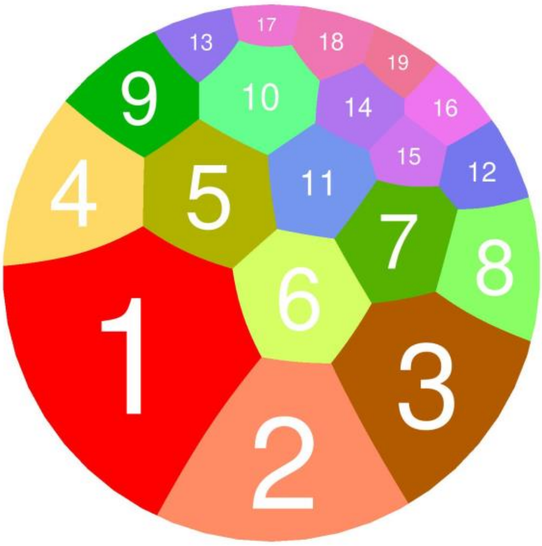


| Cluster | Gene Count | Description | Genes |
|---|---|---|---|
| 1 | 54 | Carboxylic acid catabolism | *colII, hmgA, atuA, liuR*, PA5401, PA5400, PA4979, PA4980, PA4198, PA2665, *liuA, bkdB, lpdV*, PA3416, *pdhA, lpd3*, PA4830, PA4831, *estA, bkdA1, bkdA2*, PA3415, PA3925, PA2555, PA2557, *liuB*, PA5506, PA5507, *rbsB, rbsA, ldh*, PA3723, PA2937, PA0588, PA3919, PA0656, *liuC, liuD*, PA1629, PA1628, PA1240, PA1239, *hpd, phhA, fahA, phhB*, PA2323, *gnuT, ackA*, PA0586, PA0587, PA4523, PA0109, *norC* |
| 2 | 33 | LysR, substrate-binding, and HTH-type transcriptional regulator AraC-type, N-terminal, mixed metabolism | *nmoR, ppgL, nmoA, narK1, moeA1*, PA2103, PA1413, PA3630, PA3995, PA1138, PA2497, PA4674, PA1030, PA1029, PA0207, PA3067, PA3066, PA2220, PA5264, PA2100, PA2102, *glpK, glpD, glpF*, PA3124, PA3123, PA2846, PA2845, PA0491, pxpA1, PA3225, PA0205, PA1137 |
| 3 | 28 | Mixed, incl. UspA, and Alcohol dehydrogenase, zinc-type, conserved site | PA4463, PA4611, PA4577, PA5475, PA5232, PA5027, PA4610, PA3572, PA3614, *rfaD*, PA3309, PA2753, PA2754, PA1789, PA2501, PA2805, PA1673, PA1414, PA4535, PA0862, *calB*, PA0365, PA1196, PA1603, PA1604, PA1602, PA1601, PA0200 |
| 4 | 17 | Mostly uncharacterized, incl. Nitrotoluene degradation, and Flavodoxin-like fold | *nuoA1*, PA3132, PA3133, PA4832, PA4107, *cifR, morB*, PA1285, PA2299, PA2298, PA2297, PA2292, PA1286, PA1289, PA1290, PA2590, PA2589 |
| 5 | 16 | Histidine catabolism, and Glycine betaine transport ATP-binding subunit | *dauA, dauR*, PA5099, *hutU*, PA5106, *hutC*, PA5104, *hutH*, PA5097, PA5096, PA5095, PA3187, PA3190, PA2372, *gbuA, dauB* |

| | | | |
|---|---|---|---|
| 6 | 15 | Purine nucleobase metabolism | *glcC, glcE*, PA1499, PA1500, *gcl*, PA1518, PA2938, PA1516, PA1517, *alc*, PA0134, PA0121, PA4353, PA4354, PA0120 |
| 7 | 12 | Mixed, incl. Virion | PA0724, PA0725, PA0726, PA0727, PA0728, *coaB*, PA0722, PA0721, PA0720, PA0719, PA0718, PA0717 |
| 8 | 12 | Negative regulation of transmembrane transport, and Positive regulation of transporter activity | *msrB, cyaY*, PA2826, PA4289, PA4288, *ospR, armR*, PA3720, *nalC, mexR*, PA3718, PA3677 |
| 9 | 11 | Mixed, incl. Predicted metal-dependent hydrolase, and MerR HTH family regulatory protein | PA3971, PA3972, PA3973, PA4108, PA4596, PA2577, *prtN*, PA1538, PA1537, PA0830, PA3323 |
| 10 | 11 | Bacterial chemotaxis;Transducer | *bfiS, bfiR, aer*, PA4633, PA4359, *pctA, bdlA*, PA0179, PA1930, PA1888, PA2479 |
| 11 | 10 | Amino acid transport, D-alanine transport, and Leucine-binding protein domain | *braC*, PA4911, PA4913, PA4910, PA1260, PA1261, PA1256, PA5153, *braD*, PA3865 |
| 12 | 6 | Mixed, incl. ACT domain , and B3/4 domain | PA0218, *iciA*, PA4365, PA4364, PA4182, PA4181 |
| 13 | 4 | Biofilm formation - Pseudomonas aeruginosa, and Tetratricopeptide repeat | PA0080, PA0089, *clpV1*, PA0083 |
| 14 | 4 | Mixed, incl. Helix-turn-helix XRE-family like proteins, and Bacterial extracellular solute-binding protein | PA0534, PA0535, PA1884, PA1885 |
| 15 | 2 | | PA0359, PA2136 |
| 16 | 2 | Mixed, incl. 2-thiouracil desulfurase, and Oxoglutarate/iron-dependent dioxygenase | PA3306, PA3307 |
| 17 | 2 | | PA4070, PA4071 |
| 18 | 2 | Mixed, incl. Helix_turn_helix, mercury resistance, and Chromophore | PA4657, PA4658 |
| 19 | 2 | | PA4903, *vanA* |

***Figure S5.*** **Markov clustering of transcripts displaying decreased abundance in the Δ*lasR*::Tc$^R$ mutant during growth in poly-culture. The data clusters are representated as STRING maps and as Voronoi tesselations. The colour coding in the STRING maps and Voronoi tesselations is the same, and the identities of the modulated transcripts in each cluster are shown in the colour-coded table below the Figure.**

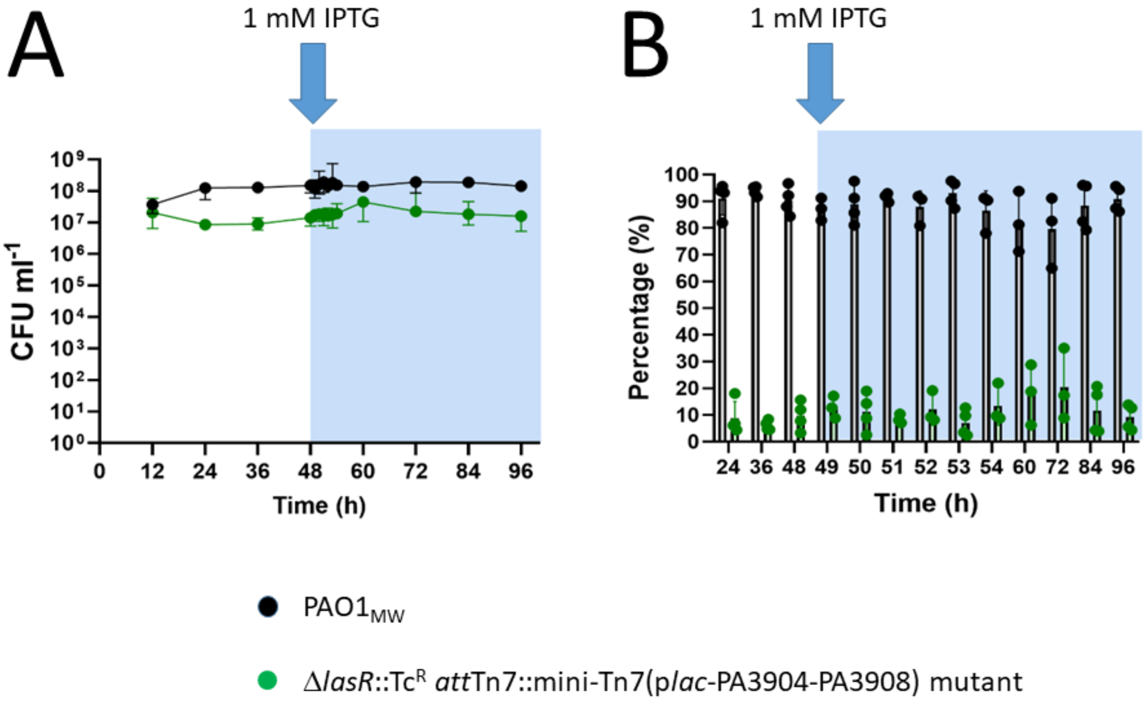


***Figure S6*. Externally-induced expression of the *tsi* cluster in the Δ*lasR*::Tc$^R$ *att*Tn7::mini-Tn7(p*lac*-PA3904-PA3908) mutant does not change competitiveness with respect to PAO1$_{MW}$ in mono-species co-cultures.** (**A**) Mono-species co-culture of PAO1$_{MW}$ (●) and the Δ*lasR*::Tc$^R$ *att*Tn7::mini-Tn7(p*lac*-PA3904-PA3908) mutant (●). IPTG (1 mM) was added to the co-cultures after 48 h, as indicated, and was continuously present in the input medium thereafter. CFU mL$^{-1}$ values are plotted on a log$_{10}$ scale and the data are presented as the median with 95% confidence intervals based on four independent experiments. Both strains were initially introduced (at t = 0 h) at an OD$_{600}$ of 0.05. ASM was supplied at a flow rate of 145 µL min$^{-1}$. Note the higher frqeuency of sampling shortly after the IPTG was added. (**B**) Detailed relative abundances of PAO1$_{MW}$ and the Δ*lasR*::Tc$^R$ *att*Tn7::mini-Tn7(p*lac*-PA3904-PA3908) mutant over time. The mean relative abundance of the Δ*lasR*::Tc$^R$ *att*Tn7::mini-Tn7(p*lac*-PA3904-PA3908) mutant was 8.2 ± 4.5% before induction of *tsi* cluster expression, and 12.0 ± 7.3% after induction, with no significant difference between the two datasets.

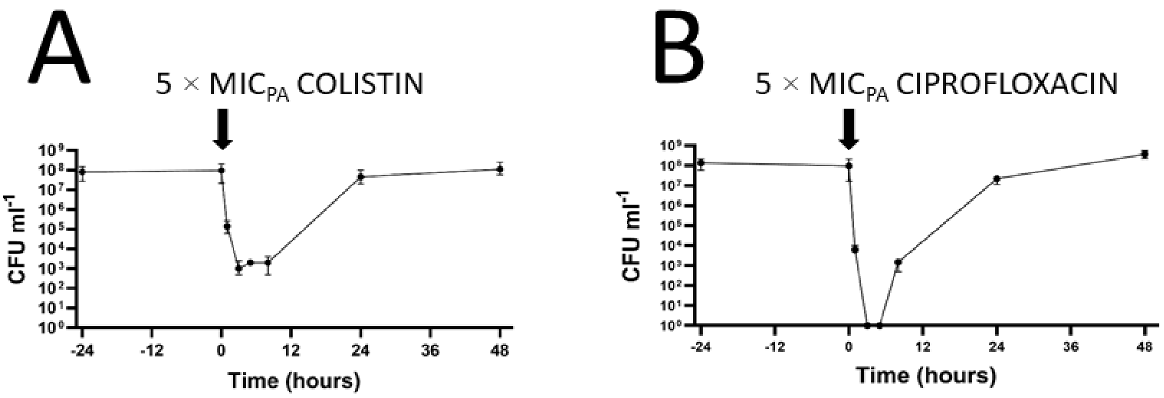


***Figure S7.*** **Effect of 5xMIC$_{PA}$ colistin and ciprofloxacin on *Pseudomonas aeruginosa* PAO1$_{MW}$ mono-cultures in the continuous-flow system.** Steady-state mono-species cultures containing PAO1$_{MW}$ (●) were grown for 48 h and then challenged (at "t = 0 h", indicated by the black arrow on the graph) with a pulse of 5 × MIC$_{P. aeruginosa}$ colistin (left panel) or 5 × MIC$_{P. aeruginosa}$ ciprofloxacin (right panel). Note that for clarity, we are only showing here the population dynamics in the 24 h period prior to antibiotic addition. Data are from six independent experiments and show the median CFU mL$^{-1}$ with a 95% confidence interval. Both cultures were grown in ASM with a fluid replacement rate of 145 µL min$^{-1}$.

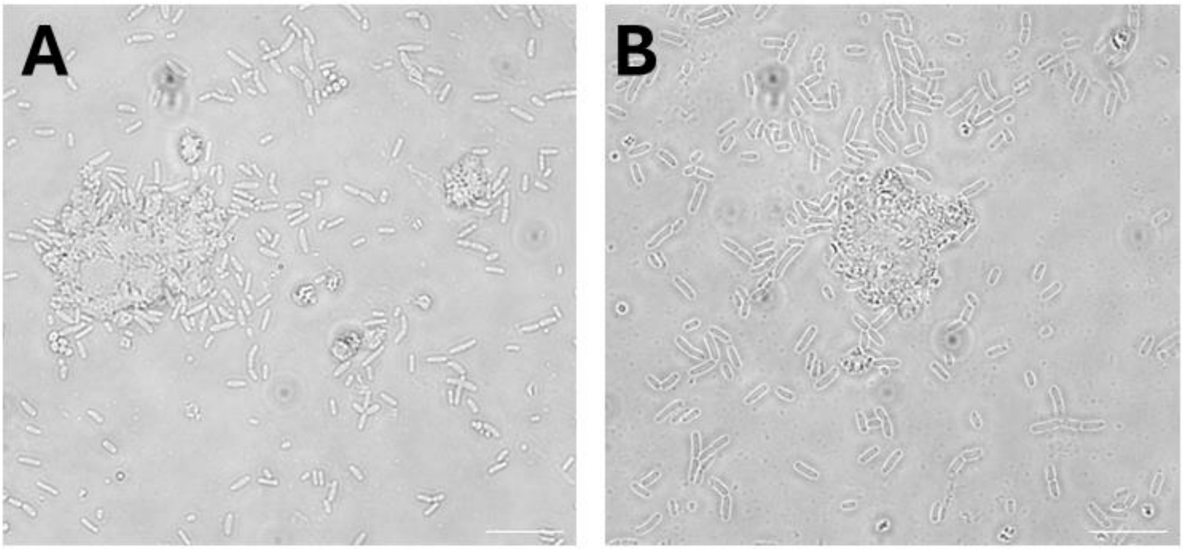


***Figure S8.*** **Phase-contrast microscopy images of aggregates formed during steady-state growth in the *in vitro* continuous-flow system.** Aggregate formation was observed in non-antibiotic perturbed cultures, with free-living cells typically surrounding the structures (A-B). Images were taken using an Olympus BX51 microscope equipped with a QICAM Fast 1394 camera. Scale bars represent 10 µm.

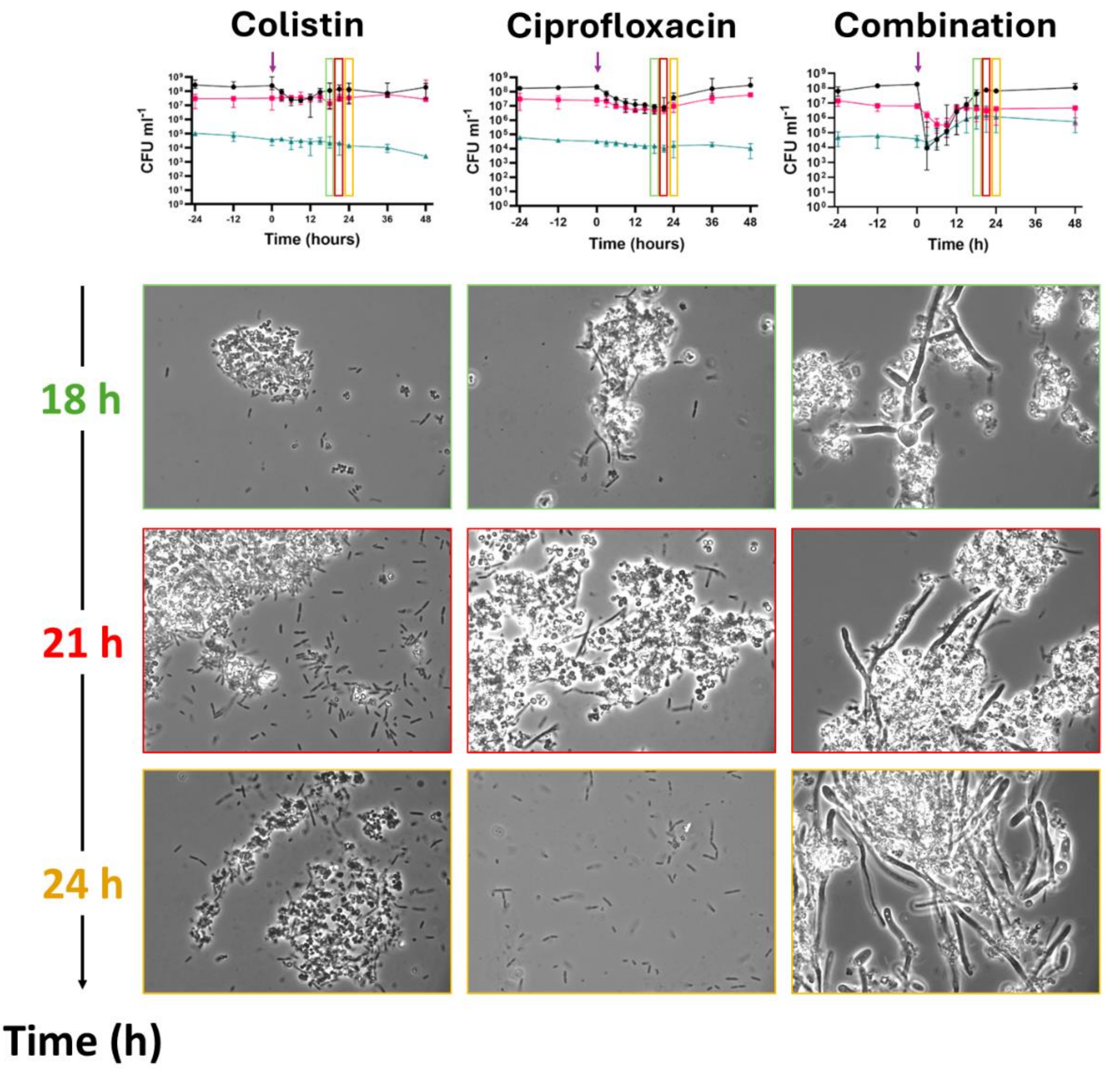


***Figure S9.*** **Phase contrast microscopy reveals blooming of *C. albicans* hyphae in poly-cultures following treatment with 5 × MIC$_{PA}$ [colistin + ciprofloxacin].** Upper panels show the steady state cultures dynamics shown in **Figure 5A-C**, with *S. aureus* (■), *C. albicans* (▲), and PAO1$_{MW}$ (●). The cultures were challenged with the indicated antibiotics, added at the time point shown by the arrow. Aliquots of cultures were then removed for phase contrast microscopy at 18 h (green box), 21 h (red box), and 24 h (yellow box) following the antibiotic challenge. Cell aggregation is observed in all treatment scenarios; however, the combined antibiotic treatment induces a visible hyphal blooming by the *C. albicans*.
