## Supplementary datasets for "*Pseudomonas aeruginosa lasR* is a keystone gene in polymicrobial cultures"

### *Supplementary Dataset A.*

**List of $PAO1_{MW}$ transcripts with significant differences in abundance in poly- (+) *versus* mono-cultures (-)**

| Gene | Name | Product | $log_2$ FC | Adjusted p-value |
|---|---|---|---|---|
| PA0075 | *pppA* | PppA | -1.046 | 4.13E-03 |
| PA0079 | *tssK1* | TssK1 | -1.120 | 7.48E-05 |
| PA0080 | *tssJ1* | TssJ1 | -1.299 | 1.94E-04 |
| PA0083 | *tssB1* | TssB1 | -1.284 | 3.58E-07 |
| PA0089 | *tssG1* | TssG1 | -1.179 | 4.75E-03 |
| PA0120 | probable transcriptional regulator | NA | -1.189 | 1.60E-04 |
| PA0121 | hypothetical protein | NA | -1.213 | 6.16E-07 |
| PA0534 | *pauB1* | NA | -1.237 | 2.78E-05 |
| PA0603 | *agtA* | AgtA | 1.523 | 4.19E-10 |
| PA0907 | *alpA* | NA | -1.038 | 3.32E-06 |
| PA1072 | *braE* | branched-chain amino acid transport protein BraE | -1.001 | 6.37E-05 |
| PA1073 | *braD* | branched-chain amino acid transport protein BraD | -1.080 | 1.74E-05 |
| PA1260 | *lhpP* | NA | -1.134 | 4.98E-05 |
| PA1320 | *cyoD* | cytochrome o ubiquinol oxidase subunit IV | 1.024 | 7.26E-03 |
| PA1429 | probable cation-transporting P-type ATPase | NA | -1.072 | 1.78E-07 |
| PA1500 | probable oxidoreductase | probable oxidoreductase | -1.319 | 2.29E-04 |
| PA1514 | ureidoglycolate hydrolaseYbbT | ureidoglycolate hydrolaseYbbT | -1.053 | 3.64E-07 |
| PA1515 | *alc* | allantoicase | -1.351 | 4.19E-10 |
| PA1516 | hypothetical protein | NA | -1.207 | 3.82E-10 |
| PA1517 | conserved hypothetical protein | NA | -1.286 | 2.86E-13 |
| PA1602 | probable oxidoreductase | probable oxidoreductase | -1.009 | 3.77E-06 |
| PA1854 | conserved hypothetical protein | NA | 1.675 | 9.82E-09 |

| Gene | Name | Product | $\log_2$ FC | Adjusted p-value |
|---|---|---|---|---|
| PA1856 | probable cytochrome oxidase subunit | probable cytochrome oxidase subunit | 1.854 | 4.03E-02 |
| PA1864 | probable transcriptional regulator | NA | -1.054 | 4.26E-04 |
| PA2007 | *maiA* | maleylacetoacetate isomerase | -1.421 | 7.13E-04 |
| PA2008 | *fahA* | fumarylacetoacetase | -1.413 | 1.22E-05 |
| PA2009 | *hmgA* | homogentisate 1 | -1.415 | 4.87E-02 |
| PA2013 | *liuC* | putative 3-methylglutaconyl-CoA hydratase | -1.104 | 4.84E-03 |
| PA2014 | *liuB* | methylcrotonyl-CoA carboxylase | -1.384 | 1.38E-06 |
| PA2015 | *liuA* | putative isovaleryl-CoA dehydrogenase | -1.418 | 4.74E-09 |
| PA2016 | *liuR* | NA | -1.197 | 5.69E-05 |
| PA2081 | *kynB* | kynurenine formamidase | -1.146 | 1.39E-04 |
| PA2247 | *bkdA1* | 2-oxoisovalerate dehydrogenase (alpha subunit) | -2.149 | 4.27E-18 |
| PA2248 | *bkdA2* | 2-oxoisovalerate dehydrogenase (beta subunit) | -2.209 | 6.45E-21 |
| PA2249 | *bkdB* | branched-chain alpha-keto acid dehydrogenase (lipoamide component) | -1.763 | 1.75E-10 |
| PA2250 | *lpdV* | lipoamide dehydrogenase-Val | -1.547 | 1.36E-09 |
| PA2442 | *gcvT2* | glycine cleavage system protein T2 | 1.159 | 6.59E-06 |
| PA2444 | *glyA2* | serine hydroxymethyltransferase | 1.069 | 1.26E-03 |
| PA2756 | hypothetical protein | NA | 1.145 | 3.96E-05 |
| PA2757 | hypothetical protein | NA | 1.405 | 2.53E-06 |
| PA2937 | hypothetical protein | NA | -1.003 | 1.32E-04 |

| Gene | Name | Product | $log_2$ FC | Adjusted p-value |
|---|---|---|---|---|
| PA3415 | probable dihydrolipoamide acetyltransferase | probable dihydrolipoamide acetyltransferase | -1.034 | 1.06E-02 |
| PA3416 | probable pyruvate dehydrogenase E1 component | probable pyruvate dehydrogenase E1 component | -1.212 | 1.01E-03 |
| PA3417 | probable pyruvate dehydrogenase E1 component | probable pyruvate dehydrogenase E1 component | -1.261 | 2.29E-04 |
| PA3608 | *potB* | polyamine transport protein PotB | 1.015 | 1.66E-04 |
| PA3609 | *potC* | polyamine transport protein PotC | 1.102 | 1.97E-05 |
| PA3865 | putative periplasmic lysine- | putative periplasmic lysine- | -1.106 | 1.40E-08 |
| PA3919 | conserved hypothetical protein | NA | -1.107 | 6.79E-07 |
| PA4033 | *mucE* | NA | 1.079 | 7.83E-06 |
| PA4034 | *aqpZ* | NA | 1.051 | 6.63E-06 |
| PA4181 | hypothetical protein | NA | -1.263 | 2.82E-04 |
| PA4182 | hypothetical protein | NA | -1.162 | 1.02E-05 |
| PA4288 | probable transcriptional regulator | NA | -1.363 | 7.37E-11 |
| PA4290 | probable chemotaxis transducer | probable chemotaxis transducer | -1.076 | 1.07E-03 |
| PA4309 | *pctA* | chemotactic transducer PctA | -1.104 | 2.30E-07 |
| PA4364 | hypothetical protein | NA | -2.044 | 5.48E-09 |
| PA4365 | *lysE* | NA | -1.861 | 5.48E-09 |
| PA4596 | *esrC* | NA | -1.225 | 2.76E-04 |
| PA4633 | probable chemotaxis transducer | probable chemotaxis transducer | -1.310 | 9.14E-08 |
| PA4828 | conserved hypothetical protein | NA | -1.163 | 2.02E-04 |
| PA4829 | *lpd3* | dihydrolipoamide dehydrogenase 3 | -1.072 | 1.02E-04 |

| Gene | Name | Product | $\log_2$ FC | Adjusted p-value |
|---|---|---|---|---|
| PA4913 | probable binding protein component of ABC transporter | probable binding protein component of ABC transporter | -1.049 | 5.31E-07 |
| PA4916 | *nrtR* | NA | -1.120 | 1.51E-04 |
| PA4918 | *pcnA* | nicotinamidase | -1.094 | 5.92E-09 |
| PA5098 | *hutH* | histidine ammonia-lyase | -1.155 | 9.81E-05 |
| PA5099 | probable transporter | NA | -1.342 | 1.46E-06 |
| PA5100 | *hutU* | urocanase | -1.430 | 5.48E-09 |
| PA5104 | conserved hypothetical protein | NA | -1.216 | 4.60E-04 |
| PA5105 | *hutC* | NA | -1.182 | 4.74E-09 |
| PA5106 | conserved hypothetical protein | conserved hypothetical protein | -1.646 | 3.63E-14 |
| PA5112 | *estA* | NA | -1.056 | 5.48E-09 |
| PA5401 | hypothetical protein | NA | -1.229 | 3.09E-05 |
| PA5507 | hypothetical protein | NA | -1.021 | 6.97E-07 |

### *Supplementary Dataset B.*

**List of transcripts from the Δ*lasR*::Tc$^R$ mutant showing significant differences in abundance in poly-(+) *versus* mono-cultures (-).**

| Gene | Name | Product | $\log_2$ FC | Adjusted p-value |
|---|---|---|---|---|
| PA0050 | hypothetical protein | NA | -1.385 | 1.74E-06 |
| PA0080 | *tssJ1* | TssJ1 | -1.551 | 2.35E-04 |
| PA0083 | *tssB1* | TssB1 | -1.327 | 1.74E-03 |
| PA0089 | *tssG1* | TssG1 | -1.354 | 2.10E-02 |
| PA0090 | *clpV1* | ClpV1 | -1.592 | 4.12E-03 |
| PA0108 | *coIII* | cytochrome c oxidase | -1.41 | 4.92E-04 |
| PA0109 | hypothetical protein | NA | -1.359 | 1.52E-05 |
| PA0120 | probable transcriptional regulator | NA | -1.542 | 9.19E-03 |
| PA0121 | hypothetical protein | NA | -1.749 | 1.23E-03 |
| PA0134 | probable guanine deaminase | probable guanine deaminase | -1.754 | 6.46E-05 |

| Gene | Name | Product | $log_2$ FC | Adjusted p-value |
|---|---|---|---|---|
| PA0160 | hypothetical protein | NA | 2.793 | 5.81E-06 |
| PA0171 | *siaB* | NA | 1.408 | 9.20E-05 |
| PA0172 | *siaA* | NA | 1.962 | 5.45E-08 |
| PA0179 | probable two-component response regulator | probable two-component response regulator | -1.714 | 1.13E-08 |
| PA0200 | hypothetical protein | NA | -1.898 | 2.81E-03 |
| PA0205 | probable permease of ABC transporter | probable permease of ABC transporter | -1.56 | 5.62E-04 |
| PA0207 | probable transcriptional regulator | NA | -1.308 | 1.26E-03 |
| PA0218 | probable transcriptional regulator | NA | -1.826 | 8.52E-06 |
| PA0359 | hypothetical protein | NA | -1.587 | 2.84E-11 |
| PA0365 | *laoB* | NA | -1.467 | 4.50E-08 |
| PA0366 | *laoC* | NA | -2.07 | 1.29E-13 |
| PA0382 | *micA* | NA | 1.65 | 2.12E-06 |
| PA0388 | hypothetical protein | NA | -1.406 | 4.69E-07 |
| PA0416 | *chpD* | NA | 1.471 | 5.01E-04 |
| PA0424 | *mexR* | multidrug resistance operon repressor MexR | -1.423 | 3.53E-04 |
| PA0491 | probable transcriptional regulator | NA | -1.724 | 3.37E-04 |
| PA0492 | conserved hypothetical protein | NA | -1.463 | 2.67E-03 |
| PA0523 | *norC* | nitric-oxide reductase subunit C | -1.385 | 1.45E-02 |
| PA0529 | conserved hypothetical protein | NA | 1.641 | 3.43E-03 |
| PA0530 | probable class III pyridoxal phosphate-dependent aminotransferase | probable class III pyridoxal phosphate-dependent aminotransferase | 1.757 | 1.87E-04 |

| Gene | Name | Product | $log_2$ FC | Adjusted p-value |
|---|---|---|---|---|
| PA0531 | probable glutamine amidotransferase | NA | 1.682 | 7.81E-05 |
| PA0534 | *pauB1* | NA | -2.997 | 1.57E-06 |
| PA0535 | transcriptional regulator | NA | -2.112 | 5.70E-06 |
| PA0578 | conserved hypothetical protein | NA | 1.469 | 7.24E-07 |
| PA0586 | conserved hypothetical protein | NA | -1.486 | 2.10E-16 |
| PA0587 | conserved hypothetical protein | NA | -1.704 | 8.51E-16 |
| PA0588 | conserved hypothetical protein | NA | -1.987 | 1.29E-13 |
| PA0596 | hypothetical protein | hypothetical protein | -1.415 | 2.01E-05 |
| PA0603 | *agtA* | AgtA | 1.92 | 3.55E-04 |
| PA0608 | probable phosphoglycolate phosphatase | probable phosphoglycolate phosphatase | 1.385 | 1.84E-06 |
| PA0610 | *prtN* | NA | -1.715 | 7.07E-07 |
| PA0656 | probable HIT family protein | NA | -1.442 | 2.05E-06 |
| PA0673 | hypothetical protein | NA | -1.495 | 3.95E-03 |
| PA0716.2 | *xisF4* | NA | -3.047 | 3.00E-02 |
| PA0717 | hypothetical protein of bacteriophage Pf1 | NA | -3.067 | 2.88E-02 |
| PA0718 | hypothetical protein of bacteriophage Pf1 | NA | -3.609 | 7.90E-03 |
| PA0719 | hypothetical protein of bacteriophage Pf1 | NA | -4.641 | 6.71E-04 |
| PA0720 | helix destabilizing protein of bacteriophage Pf1 | NA | -5.979 | 7.84E-05 |
| PA0721 | *pfsE* | NA | -4.546 | 4.45E-03 |
| PA0722 | hypothetical protein of | NA | -4.251 | 1.98E-03 |

| Gene | Name | Product | $log_2$ FC | Adjusted p-value |
|---|---|---|---|---|
| | bacteriophage Pf1 | | | |
| PA0723 | *coaB* | NA | -5.476 | 5.98E-04 |
| PA0724 | probable coat protein A of bacteriophage Pf1 | NA | -3.227 | 2.22E-03 |
| PA0725 | hypothetical protein of bacteriophage Pf1 | NA | -2.148 | 9.79E-04 |
| PA0726 | hypothetical protein of bacteriophage Pf1 | NA | -2.313 | 1.48E-04 |
| PA0727 | Pf replication initiator protein | NA | -2.284 | 3.32E-04 |
| PA0728 | probable bacteriophage integrase | NA | -1.577 | 2.43E-03 |
| PA0830 | hypothetical protein | NA | -1.764 | 5.95E-10 |
| PA0836 | *ackA* | acetate kinase | -1.847 | 5.44E-05 |
| PA0840 | probable oxidoreductase | NA | 1.431 | 5.66E-07 |
| PA0862 | hypothetical protein | NA | -1.437 | 4.00E-11 |
| PA0865 | *hpd* | 4-hydroxyphenylpyruvate dioxygenase | -2.171 | 1.85E-04 |
| PA0871 | *phhB* | NA | -1.475 | 3.19E-05 |
| PA0872 | *phhA* | phenylalanine-4-hydroxylase | -2.077 | 3.59E-11 |
| PA0907 | *alpA* | NA | -2.372 | 6.52E-05 |
| PA0914 | hypothetical protein | NA | 1.611 | 5.03E-05 |
| PA0915 | conserved hypothetical protein | NA | 1.337 | 3.15E-04 |
| PA0918 | cytochrome b561 | NA | -1.424 | 3.25E-07 |
| PA0942 | probable transcriptional regulator | NA | -1.384 | 3.75E-09 |
| PA0978 | conserved hypothetical protein | NA | 1.579 | 6.84E-04 |
| PA0984 | colicin immunity protein | NA | -1.439 | 1.12E-03 |

| Gene | Name | Product | $log_2$ FC | Adjusted p-value |
|---|---|---|---|---|
| PA0986 | conserved hypothetical protein | NA | -1.744 | 5.52E-10 |
| PA1003 | *mvfR* | Transcriptional regulator MvfR | -1.492 | 8.86E-12 |
| PA1029 | hypothetical protein | NA | -1.739 | 1.80E-04 |
| PA1030 | hypothetical protein | NA | -1.804 | 1.97E-05 |
| PA1073 | *braD* | branched-chain amino acid transport protein BraD | -1.318 | 8.30E-04 |
| PA1074 | *braC* | branched-chain amino acid transport protein BraC | -1.957 | 2.97E-08 |
| PA1137 | probable oxidoreductase | NA | -1.664 | 1.61E-04 |
| PA1138 | probable transcriptional regulator | NA | -1.352 | 3.26E-05 |
| PA1151 | *imm2* | NA | -1.544 | 8.68E-06 |
| PA1183 | *dctA* | C4-dicarboxylate transport protein | 1.778 | 1.37E-08 |
| PA1194 | probable amino acid permease | NA | 1.4 | 2.95E-05 |
| PA1196 | *ddaR* | NA | -1.695 | 2.03E-03 |
| PA1210 | conserved hypothetical protein | NA | -1.854 | 3.81E-07 |
| PA1223 | probable transcriptional regulator | NA | -1.323 | 6.30E-05 |
| PA1229 | probable transcriptional regulator | NA | -1.412 | 1.72E-04 |
| PA1239 | hypothetical protein | NA | -1.636 | 5.84E-05 |
| PA1240 | probable enoyl-CoA hydratase/isomerase | NA | -1.738 | 1.66E-04 |
| PA1256 | *lhpO* | NA | -1.565 | 5.02E-04 |
| PA1260 | *lhpP* | NA | -2.695 | 2.46E-09 |
| PA1261 | *lhpR* | NA | -1.564 | 1.76E-04 |
| PA1265 | hypothetical protein | NA | 1.782 | 4.08E-02 |

| Gene | Name | Product | $log_2$ FC | Adjusted p-value |
|---|---|---|---|---|
| PA1267 | *lhpB* | D-hydroxyproline dehydrogenase beta-subunit | 1.894 | 3.03E-02 |
| PA1285 | probable transcriptional regulator | NA | -2.64 | 1.04E-08 |
| PA1286 | probable major facilitator superfamily (MFS) transporter | NA | -2.44 | 2.09E-08 |
| PA1289 | hypothetical protein | NA | -1.907 | 1.00E-05 |
| PA1290 | probable transcriptional regulator | NA | -1.726 | 2.06E-03 |
| PA1299 | conserved hypothetical protein | NA | 1.493 | 1.57E-06 |
| PA1300 | *hxuI* | NA | 1.719 | 1.19E-05 |
| PA1301 | *hxuR* | NA | 1.598 | 3.21E-03 |
| PA1317 | *cyoA* | cytochrome o ubiquinol oxidase subunit II | 2.541 | 8.93E-07 |
| PA1318 | *cyoB* | cytochrome o ubiquinol oxidase subunit I | 2.494 | 9.54E-08 |
| PA1319 | *cyoC* | cytochrome o ubiquinol oxidase subunit III | 2.603 | 3.09E-08 |
| PA1320 | *cyoD* | cytochrome o ubiquinol oxidase subunit IV | 2.706 | 1.13E-07 |
| PA1321 | *cyoE* | cytochrome o ubiquinol oxidase protein CyoE | 2.693 | 3.75E-09 |
| PA1413 | probable transcriptional regulator | NA | -1.439 | 7.66E-04 |
| PA1414 | hypothetical protein | NA | -1.808 | 3.16E-04 |
| PA1421 | *gbuA* | guanidinobutyras e | -1.304 | 6.04E-06 |
| PA1423 | *bdlA* | BdlA | -2.094 | 1.73E-08 |
| PA1479 | *ccmE* | NA | 1.342 | 5.49E-04 |
| PA1480 | *ccmF* | NA | 1.708 | 4.30E-05 |
| PA1481 | *ccmG* | NA | 1.556 | 1.49E-04 |
| PA1482 | *ccmH* | NA | 1.769 | 3.11E-05 |
| PA1483 | *cycH* | NA | 1.876 | 8.83E-05 |

| Gene | Name | Product | $\log_2$ FC | Adjusted p-value |
|---|---|---|---|---|
| PA1499 | conserved hypothetical protein | conserved hypothetical protein | -1.375 | 1.86E-02 |
| PA1500 | probable oxidoreductase | probable oxidoreductase | -1.96 | 9.22E-06 |
| PA1502 | *gcl* | glyoxylate carboligase | -1.843 | 4.16E-06 |
| PA1515 | *alc* | allantoicase | -1.793 | 2.05E-05 |
| PA1516 | hypothetical protein | NA | -2.213 | 7.39E-05 |
| PA1517 | conserved hypothetical protein | NA | -2.535 | 4.63E-06 |
| PA1518 | conserved hypothetical protein | conserved hypothetical protein | -1.777 | 4.78E-05 |
| PA1519 | probable transporter | NA | -1.357 | 3.21E-08 |
| PA1531 | hypothetical protein | NA | 1.561 | 2.46E-09 |
| PA1537 | probable short-chain dehydrogenase | NA | -1.766 | 3.27E-06 |
| PA1538 | probable flavin-containing monooxygenase | NA | -1.927 | 2.75E-09 |
| PA1553 | *ccoO1* | Cytochrome c oxidase | 1.326 | 1.18E-03 |
| PA1554 | *ccoN1* | Cytochrome c oxidase | 1.435 | 5.95E-10 |
| PA1561 | *aer* | aerotaxis receptor Aer | -1.454 | 5.98E-04 |
| PA1601 | probable aldehyde dehydrogenase | NA | -1.815 | 9.72E-05 |
| PA1602 | probable oxidoreductase | probable oxidoreductase | -2.185 | 2.02E-07 |
| PA1603 | probable transcriptional regulator | NA | -2.183 | 1.25E-11 |
| PA1604 | hypothetical protein | NA | -2.035 | 7.62E-12 |
| PA1628 | probable 3-hydroxyacyl-CoA dehydrogenase | probable 3-hydroxyacyl-CoA dehydrogenase | -1.525 | 3.69E-05 |
| PA1629 | probable enoyl-CoA hydratase/isomerase | NA | -1.566 | 1.30E-06 |

| Gene | Name | Product | $log_2$ FC | Adjusted p-value |
|---|---|---|---|---|
| PA1639 | hypothetical protein | NA | 1.4 | 2.95E-04 |
| PA1649 | probable short-chain dehydrogenase | NA | 1.569 | 3.07E-03 |
| PA1673 | *mhr* | NA | -1.572 | 2.69E-03 |
| PA1688 | hypothetical protein | NA | 1.775 | 7.78E-06 |
| PA1689 | conserved hypothetical protein | NA | 1.464 | 2.02E-04 |
| PA1766 | hypothetical protein | NA | 1.451 | 1.07E-04 |
| PA1771 | *estX* | NA | 1.895 | 5.63E-06 |
| PA1774 | *crfX* | NA | 1.329 | 8.55E-07 |
| PA1775 | *cmpX* | NA | 1.47 | 4.50E-06 |
| PA1789 | hypothetical protein | NA | -1.689 | 4.08E-04 |
| PA1824 | conserved hypothetical protein | NA | 1.319 | 2.35E-03 |
| PA1835 | hypothetical protein | NA | -1.595 | 2.83E-06 |
| PA1848 | probable major facilitator superfamily (MFS) transporter | NA | 1.51 | 4.54E-03 |
| PA1853 | probable transcriptional regulator | NA | 2.22 | 5.20E-09 |
| PA1854 | conserved hypothetical protein | NA | 3.305 | 2.21E-21 |
| PA1855 | hypothetical protein | NA | 3.48 | 6.05E-06 |
| PA1856 | probable cytochrome oxidase subunit | probable cytochrome oxidase subunit | 2.219 | 6.70E-06 |
| PA1864 | probable transcriptional regulator | NA | -2.308 | 2.49E-05 |
| PA1883 | probable NADH-ubiquinone/plastoquinone oxidoreductase | probable NADH-ubiquinone/plastoquinone oxidoreductase | -1.509 | 1.85E-04 |
| PA1884 | transcriptional regulator | NA | -1.303 | 2.77E-02 |

| Gene | Name | Product | $log_2$ FC | Adjusted p-value |
|---|---|---|---|---|
| PA1885 | conserved hypothetical protein | NA | -2.436 | 8.10E-08 |
| PA1888 | hypothetical protein | NA | -1.431 | 1.78E-09 |
| PA1926 | Uncharacterized protein | NA | 1.326 | 1.73E-08 |
| PA1930 | probable chemotaxis transducer | probable chemotaxis transducer | -1.308 | 1.02E-05 |
| PA1946 | *rbsB* | binding protein component precursor of ABC ribose transporter | -2.024 | 1.08E-18 |
| PA1947 | *rbsA* | ribose transport protein RbsA | -2.355 | 8.12E-12 |
| PA1982 | *exaA* | quinoprotein ethanol dehydrogenase | 1.561 | 4.32E-02 |
| PA2008 | *fahA* | fumarylacetoacetase | -1.561 | 3.44E-03 |
| PA2009 | *hmgA* | homogentisate 1 | -2.619 | 2.40E-07 |
| PA2012 | *liuD* | methylcrotonyl-CoA carboxylase | -1.326 | 2.57E-04 |
| PA2013 | *liuC* | putative 3-methylglutaconyl-CoA hydratase | -1.631 | 1.19E-05 |
| PA2014 | *liuB* | methylcrotonyl-CoA carboxylase | -1.952 | 1.99E-07 |
| PA2015 | *liuA* | putative isovaleryl-CoA dehydrogenase | -2.6 | 6.32E-07 |
| PA2016 | *liuR* | NA | -2.618 | 1.53E-05 |
| PA2042 | probable transporter (membrane subunit) | NA | 1.916 | 1.19E-05 |
| PA2043 | hypothetical protein | NA | 1.635 | 5.62E-04 |
| PA2064 | *pcoB* | NA | 1.565 | 5.02E-03 |
| PA2099 | probable short-chain dehydrogenase | NA | -1.41 | 2.21E-03 |
| PA2100 | probable transcriptional regulator | NA | -1.502 | 4.02E-03 |
| PA2102 | hypothetical protein | NA | -1.559 | 2.54E-04 |

| Gene | Name | Product | $\log_2$ FC | Adjusted p-value |
|---|---|---|---|---|
| PA2103 | probable molybdopterin biosynthesis protein MoeB | probable molybdopterin biosynthesis protein MoeB | -1.404 | 4.23E-04 |
| PA2118 | *ada* | NA | 1.549 | 4.25E-04 |
| PA2128 | *cupA1* | NA | 3.605 | 5.39E-03 |
| PA2129 | *cupA2* | NA | 5.186 | 1.88E-04 |
| PA2130 | *cupA3* | NA | 3.85 | 3.26E-03 |
| PA2131 | *cupA4* | NA | 4.902 | 6.17E-04 |
| PA2132 | *cupA5* | NA | 5.203 | 2.35E-05 |
| PA2133 | Cyclic-guanylate-specific phosphodiesterase | NA | 4.303 | 1.68E-03 |
| PA2134 | hypothetical protein | NA | 3.191 | 7.34E-03 |
| PA2136 | hypothetical protein | NA | -1.854 | 1.40E-06 |
| PA2187 | hypothetical protein | NA | -1.362 | 2.34E-03 |
| PA2202 | probable amino acid permease | NA | 1.621 | 1.20E-04 |
| PA2220 | probable transcriptional regulator | NA | -1.378 | 3.02E-03 |
| PA2221 | conserved hypothetical protein | NA | -1.64 | 9.93E-04 |
| PA2238 | *pslH* | PslH | 1.645 | 8.63E-06 |
| PA2239 | *pslI* | PslI | 1.505 | 4.04E-05 |
| PA2240 | *pslJ* | PslJ | 1.758 | 5.95E-10 |
| PA2242 | *pslL* | hypothetical protein | 1.403 | 3.40E-07 |
| PA2245 | *pslO* | NA | -1.627 | 2.32E-03 |
| PA2247 | *bkdA1* | 2-oxoisovalerate dehydrogenase (alpha subunit) | -3.924 | 1.91E-21 |
| PA2248 | *bkdA2* | 2-oxoisovalerate dehydrogenase (beta subunit) | -3.636 | 3.64E-12 |
| PA2249 | *bkdB* | branched-chain alpha-keto acid dehydrogenase (lipoamide component) | -2.92 | 4.22E-10 |
| PA2250 | *lpdV* | lipoamide dehydrogenase-Val | -2.529 | 1.63E-12 |

| Gene | Name | Product | $log_2$ FC | Adjusted p-value |
|---|---|---|---|---|
| PA2277 | *arsR* | NA | -1.804 | 2.18E-06 |
| PA2292 | hypothetical protein | NA | -1.687 | 5.24E-05 |
| PA2297 | probable ferredoxin | NA | -1.306 | 2.47E-03 |
| PA2298 | probable oxidoreductase | NA | -1.702 | 1.05E-04 |
| PA2299 | probable transcriptional regulator | NA | -1.639 | 5.71E-05 |
| PA2322 | *gntP* | NA | -2.197 | 6.49E-04 |
| PA2323 | *gapN* | GapN | -2.236 | 4.49E-04 |
| PA2372 | hypothetical protein | NA | -1.336 | 6.43E-06 |
| PA2411 | probable thioesterase | NA | 2.592 | 4.91E-02 |
| PA2413 | *pvdH* | L-2 | 2.306 | 1.95E-02 |
| PA2424 | *pvdL* | NA | 1.601 | 3.76E-02 |
| PA2440 | hypothetical protein | NA | 1.347 | 4.00E-02 |
| PA2442 | *gcvT2* | glycine cleavage system protein T2 | 1.532 | 1.85E-03 |
| PA2443 | *sdaA* | L-serine dehydratase | 1.345 | 8.06E-03 |
| PA2444 | *glyA2* | serine hydroxymethyltr ansferase | 1.703 | 8.82E-04 |
| PA2479 | *dsbR* | NA | -1.343 | 4.50E-06 |
| PA2485 | hypothetical protein | NA | -1.355 | 2.28E-03 |
| PA2497 | probable transcriptional regulator | NA | -1.302 | 3.67E-06 |
| PA2501 | hypothetical protein | NA | -1.417 | 2.97E-02 |
| PA2521 | *czcB* | NA | -1.443 | 3.48E-02 |
| PA2531 | probable aminotransferase | probable aminotransferase | 1.445 | 5.53E-04 |
| PA2550 | probable acyl-CoA dehydrogenase | probable acyl-CoA dehydrogenase | 1.859 | 2.54E-06 |
| PA2555 | probable AMP-binding enzyme | probable AMP-binding enzyme | -1.357 | 6.10E-06 |
| PA2557 | probable AMP-binding enzyme | NA | -1.32 | 4.31E-05 |
| PA2566.1 | Uncharacterized protein | NA | -1.348 | 2.24E-04 |

| Gene | Name | Product | $log_2$ FC | Adjusted p-value |
|---|---|---|---|---|
| PA2577 | probable transcriptional regulator | NA | -1.407 | 1.07E-05 |
| PA2589 | hypothetical protein | NA | -1.574 | 1.15E-05 |
| PA2590 | hypothetical protein | NA | -1.6 | 4.51E-07 |
| PA2611 | *cysG* | siroheme synthase | 1.35 | 4.11E-05 |
| PA2645 | *nuoJ* | NADH dehydrogenase I chain J | 1.355 | 1.17E-05 |
| PA2646 | *nuoK* | NADH dehydrogenase I chain K | 1.321 | 1.50E-05 |
| PA2647 | *nuoL* | NADH dehydrogenase I chain L | 1.437 | 3.90E-06 |
| PA2648 | *nuoM* | NADH dehydrogenase I chain M | 1.506 | 3.51E-07 |
| PA2649 | *nuoN* | NADH dehydrogenase I chain N | 1.548 | 5.63E-06 |
| PA2665 | *fhpR* | NA | -1.853 | 3.18E-07 |
| PA2729 | hypothetical protein | NA | 1.44 | 4.92E-06 |
| PA2753 | hypothetical protein | NA | -1.605 | 3.60E-03 |
| PA2754 | conserved hypothetical protein | NA | -1.503 | 1.59E-03 |
| PA2756 | hypothetical protein | NA | 2.535 | 1.25E-11 |
| PA2757 | hypothetical protein | NA | 2.329 | 2.98E-08 |
| PA2769 | hypothetical protein | NA | 1.597 | 9.75E-05 |
| PA2805 | hypothetical protein | NA | -1.643 | 8.63E-06 |
| PA2825 | *ospR* | NA | -2.125 | 1.15E-04 |
| PA2826 | probable glutathione peroxidase | probable glutathione peroxidase | -2.463 | 3.34E-04 |
| PA2827 | conserved hypothetical protein | NA | -1.332 | 6.33E-04 |

| Gene | Name | Product | $\log_2$ FC | Adjusted p-value |
|---|---|---|---|---|
| PA2840 | probable ATP-dependent RNA helicase | probable ATP-dependent RNA helicase | 1.742 | 4.11E-05 |
| PA2845 | hypothetical protein | NA | -1.403 | 4.06E-02 |
| PA2846 | probable transcriptional regulator | NA | -1.517 | 2.08E-09 |
| PA2878 | hypothetical protein | NA | -1.335 | 4.12E-05 |
| PA2886 | *atuA* | NA | -1.319 | 1.85E-03 |
| PA2910 | conserved hypothetical protein | NA | 1.782 | 8.64E-05 |
| PA2931 | *cifR* | NA | -1.888 | 2.37E-05 |
| PA2932 | *morB* | morphinone reductase | -1.995 | 3.63E-03 |
| PA2937 | hypothetical protein | NA | -2.853 | 2.72E-14 |
| PA2938 | probable transporter | NA | -1.961 | 2.54E-10 |
| PA2941 | hypothetical protein | NA | -1.8 | 1.95E-05 |
| PA2964 | *pabC* | 4-amino-4-deoxychorismate lyase | 1.612 | 2.75E-04 |
| PA2968 | *fabD* | malonyl-CoA-[acyl-carrier-protein] transacylase | 1.481 | 4.63E-04 |
| PA2969 | *plsX* | fatty acid biosynthesis protein PlsX | 1.7 | 4.15E-11 |
| PA2998 | *nqrB* | NA | 1.427 | 4.48E-05 |
| PA3063 | *pelB* | PelB | 1.589 | 8.40E-04 |
| PA3066 | hypothetical protein | NA | -1.335 | 2.04E-04 |
| PA3067 | probable transcriptional regulator | NA | -1.56 | 4.17E-04 |
| PA3123 | RidA subfamily protein | NA | -1.539 | 1.05E-03 |
| PA3124 | probable transcriptional regulator | NA | -1.371 | 1.39E-03 |
| PA3132 | probable hydrolase | NA | -1.641 | 3.02E-05 |
| PA3133 | *sawR* | NA | -1.584 | 4.47E-05 |

| Gene | Name | Product | $\log_2$ FC | Adjusted p-value |
|---|---|---|---|---|
| PA3165 | *hisC2* | histidinol-phosphate aminotransferase | 1.439 | 7.40E-06 |
| PA3187 | probable ATP-binding component of ABC transporter | probable ATP-binding component of ABC transporter | -1.564 | 4.32E-03 |
| PA3190 | probable binding protein component of ABC sugar transporter | probable binding protein component of ABC sugar transporter | -2.1 | 2.53E-03 |
| PA3225 | transcriptional regulator | NA | -1.371 | 2.04E-06 |
| PA3306 | hypothetical protein | NA | -1.39 | 9.64E-08 |
| PA3307 | hypothetical protein | NA | -1.546 | 5.88E-08 |
| PA3308 | *hepA* | NA | 1.319 | 2.88E-11 |
| PA3309 | conserved hypothetical protein | NA | -1.665 | 9.93E-05 |
| PA3310 | conserved hypothetical protein | conserved hypothetical protein | -1.359 | 1.48E-04 |
| PA3321 | probable transcriptional regulator | NA | -1.331 | 1.26E-04 |
| PA3323 | conserved hypothetical protein | NA | -1.333 | 6.03E-06 |
| PA3337 | *rfaD* | ADP-L-glycero-D-mannoheptose 6-epimerase | -1.382 | 4.35E-03 |
| PA3367 | hypothetical protein | NA | 1.337 | 9.93E-05 |
| PA3394 | *nosF* | NosF protein | 1.318 | 1.05E-02 |
| PA3395 | *nosY* | NosY protein | 1.752 | 4.51E-04 |
| PA3396 | *nosL* | NA | 1.992 | 9.91E-04 |
| PA3415 | probable dihydrolipoamide acetyltransferase | probable dihydrolipoamide acetyltransferase | -1.864 | 5.06E-09 |
| PA3416 | probable pyruvate dehydrogenase E1 component | probable pyruvate dehydrogenase E1 component | -2.244 | 8.84E-14 |
| PA3417 | probable pyruvate | probable pyruvate | -2.477 | 1.29E-13 |

| Gene | Name | Product | $log_2$ FC | Adjusted p-value |
|---|---|---|---|---|
| | dehydrogenase E1 component | dehydrogenase E1 component | | |
| PA3418 | *ldh* | leucine dehydrogenase | -2.519 | 4.22E-22 |
| PA3424 | hypothetical protein | NA | 1.384 | 3.32E-02 |
| PA3572 | hypothetical protein | NA | -1.778 | 5.83E-06 |
| PA3581 | *glpF* | NA | -1.352 | 3.35E-07 |
| PA3582 | *glpK* | glycerol kinase | -1.864 | 1.19E-08 |
| PA3584 | *glpD* | glycerol-3-phosphate dehydrogenase | -2.202 | 3.65E-12 |
| PA3614 | hypothetical protein | NA | -1.536 | 7.25E-05 |
| PA3630 | *gfnR* | NA | -1.349 | 1.47E-05 |
| PA3635 | *eno* | enolase | 1.476 | 4.17E-04 |
| PA3636 | *kdsA* | 2-dehydro-3-deoxyphosphooc tonate aldolase | 1.469 | 1.62E-06 |
| PA3677 | *mexJ* | NA | -2.069 | 1.19E-16 |
| PA3693 | conserved hypothetical protein | NA | 1.345 | 1.88E-04 |
| PA3718 | probable major facilitator superfamily (MFS) transporter | NA | -1.6 | 2.09E-09 |
| PA3719 | *armR* | antirepressor for MexR | -2.95 | 1.42E-12 |
| PA3720 | hypothetical protein | NA | -2.701 | 2.92E-45 |
| PA3721 | *nalC* | NalC | -1.48 | 2.33E-08 |
| PA3723 | probable FMN oxidoreductase | NA | -1.449 | 1.64E-06 |
| PA3820 | *secF* | secretion protein SecF | 1.509 | 3.55E-05 |
| PA3821 | *secD* | secretion protein SecD | 1.4 | 2.38E-07 |
| PA3862 | *dauB* | NAD(P)H-dependent anabolic L-arginine dehydrogenase | -1.5 | 2.99E-04 |
| PA3863 | *dauA* | FAD-dependent catabolic D-arginine dehydrogenase | -1.603 | 2.43E-06 |

| Gene | Name | Product | $\log_2$ FC | Adjusted p-value |
|---|---|---|---|---|
| PA3864 | *dauR* | NA | -1.632 | 4.88E-05 |
| PA3865 | putative periplasmic lysine- | putative periplasmic lysine- | -2.762 | 4.71E-09 |
| PA3865.1 | pyocin S4 immunity protein | NA | -1.371 | 3.67E-05 |
| PA3877 | *narK1* | nitrite extrusion protein 1 | -1.663 | 3.90E-04 |
| PA3885 | *tpbA* | NA | 1.399 | 1.53E-02 |
| PA3892 | conserved hypothetical protein | NA | 1.582 | 5.48E-04 |
| PA3893 | conserved hypothetical protein | NA | 1.793 | 1.94E-05 |
| PA3894 | probable outer membrane protein precursor | NA | 1.74 | 7.05E-04 |
| PA3904 | PAAR4 | NA | 2.426 | 2.12E-06 |
| PA3914 | *moeA1* | molybdenum cofactor biosynthetic protein A1 | -1.48 | 1.10E-02 |
| PA3919 | conserved hypothetical protein | NA | -2.073 | 2.83E-06 |
| PA3925 | probable acyl-CoA thiolase | probable acyl-CoA thiolase | -1.5 | 1.44E-11 |
| PA3971 | hypothetical protein | NA | -1.554 | 3.25E-04 |
| PA3972 | probable acyl-CoA dehydrogenase | NA | -1.677 | 5.15E-07 |
| PA3973 | probable transcriptional regulator | NA | -1.874 | 4.68E-06 |
| PA3988 | *lptE* | NA | 1.372 | 1.20E-04 |
| PA3993 | probable transposase | NA | 1.543 | 1.63E-02 |
| PA3995 | probable transcriptional regulator | NA | -1.68 | 1.07E-06 |
| PA4006 | *nadD1* | nicotinate mononucleotide adenylyltransferase NadD1 | 1.309 | 2.62E-05 |
| PA4033 | *mucE* | NA | 2.155 | 5.95E-10 |
| PA4034 | *aqpZ* | NA | 2.23 | 8.58E-08 |

| Gene | Name | Product | $log_2$ FC | Adjusted p-value |
|---|---|---|---|---|
| PA4050 | *pgpA* | phosphatidylglycerophosphatase A | 1.4 | 7.91E-08 |
| PA4070 | probable transcriptional regulator | NA | -1.735 | 1.57E-12 |
| PA4071 | hypothetical protein | NA | -1.482 | 3.17E-03 |
| PA4090 | hypothetical protein | NA | -1.384 | 9.13E-03 |
| PA4107 | *efhP* | NA | -1.34 | 8.31E-03 |
| PA4108 | cyclic di-GMP phosphodiesterase | NA | -1.998 | 5.93E-08 |
| PA4139 | hypothetical protein | NA | 2.098 | 9.46E-06 |
| PA4181 | hypothetical protein | NA | -2.15 | 2.40E-09 |
| PA4182 | hypothetical protein | NA | -1.997 | 4.59E-13 |
| PA4196 | *bfiR* | NA | -1.318 | 7.01E-03 |
| PA4197 | *bfiS* | NA | -1.553 | 1.18E-03 |
| PA4198 | probable AMP-binding enzyme | NA | -1.904 | 2.14E-04 |
| PA4202 | *nmoA* | nitronate monooxygenase NmoA | -1.582 | 5.51E-09 |
| PA4203 | *nmoR* | NA | -1.706 | 2.01E-04 |
| PA4204 | *ppgL* | NA | -1.319 | 8.37E-03 |
| PA4224 | *pchG* | pyochelin biosynthetic protein PchG | 2.369 | 1.87E-02 |
| PA4229 | *pchC* | NA | 2.786 | 9.98E-03 |
| PA4230 | *pchB* | salicylate biosynthesis protein PchB | 2.915 | 1.41E-02 |
| PA4231 | *pchA* | salicylate biosynthesis isochorismate synthase | 1.667 | 1.01E-02 |
| PA4238 | *rpoA* | DNA-directed RNA polymerase alpha chain | 1.337 | 1.83E-04 |
| PA4243 | *secY* | secretion protein SecY | 1.405 | 1.13E-04 |
| PA4288 | probable transcriptional regulator | NA | -2.898 | 4.04E-08 |

| Gene | Name | Product | $log_2$ FC | Adjusted p-value |
|---|---|---|---|---|
| PA4289 | probable transporter | NA | -2.189 | 1.39E-12 |
| PA4309 | *pctA* | chemotactic transducer PctA | -2.05 | 7.11E-14 |
| PA4353 | conserved hypothetical protein | NA | -1.313 | 9.57E-06 |
| PA4354 | conserved hypothetical protein | NA | -2.001 | 3.87E-03 |
| PA4359 | conserved hypothetical protein | NA | -1.692 | 3.26E-05 |
| PA4363 | *iciA* | NA | -1.489 | 1.13E-04 |
| PA4364 | hypothetical protein | NA | -4.71 | 4.20E-24 |
| PA4365 | *lysE* | NA | -3.258 | 3.12E-20 |
| PA4368 | hypothetical protein | NA | -1.45 | 1.16E-10 |
| PA4428 | *sspA* | NA | 1.444 | 4.51E-06 |
| PA4463 | conserved hypothetical protein | NA | -1.54 | 3.97E-04 |
| PA4479 | *mreD* | NA | 1.57 | 7.93E-05 |
| PA4480 | *mreC* | NA | 1.563 | 5.59E-11 |
| PA4485 | conserved hypothetical protein | NA | 1.332 | 2.67E-03 |
| PA4523 | hypothetical protein | NA | -1.893 | 7.83E-08 |
| PA4535 | hypothetical protein | NA | -1.34 | 2.64E-06 |
| PA4577 | hypothetical protein | NA | -1.498 | 3.90E-04 |
| PA4596 | *esrC* | NA | -3.021 | 2.64E-11 |
| PA4610 | hypothetical protein | NA | -1.797 | 7.61E-04 |
| PA4611 | hypothetical protein | NA | -2.017 | 9.49E-04 |
| PA4633 | probable chemotaxis transducer | probable chemotaxis transducer | -2.122 | 5.19E-20 |
| PA4657 | hypothetical protein | NA | -1.649 | 5.13E-05 |
| PA4658 | hypothetical protein | NA | -1.978 | 9.17E-05 |
| PA4664 | *prmC* | NA | 1.634 | 5.55E-05 |
| PA4672 | peptidyl-tRNA hydrolase | NA | 1.63 | 1.60E-14 |

| Gene | Name | Product | $\log_2$ FC | Adjusted p-value |
|---|---|---|---|---|
| PA4673 | conserved hypothetical protein | NA | 1.404 | 2.52E-07 |
| PA4674 | Antitoxin HigA | NA | -1.883 | 6.08E-04 |
| PA4674.1 | *higB* | NA | -1.802 | 1.19E-04 |
| PA4683 | hypothetical protein | NA | 1.748 | 5.85E-08 |
| PA4739 | conserved hypothetical protein | NA | 1.64 | 5.98E-04 |
| PA4809 | *fdhE* | NA | 1.347 | 1.09E-03 |
| PA4810 | *fdnI* | nitrate-inducible formate dehydrogenase | 1.485 | 1.18E-04 |
| PA4828 | conserved hypothetical protein | NA | -2.289 | 5.23E-04 |
| PA4829 | *lpd3* | dihydrolipoamide dehydrogenase 3 | -2.479 | 4.16E-06 |
| PA4830 | hypothetical protein | NA | -1.973 | 9.82E-04 |
| PA4831 | probable transcriptional regulator | NA | -1.387 | 2.39E-02 |
| PA4832 | probable short-chain dehydrogenase | NA | -1.883 | 6.02E-04 |
| PA4837 | *cntO* | NA | -1.315 | 5.78E-05 |
| PA4846 | *aroQ1* | 3-dehydroquinate dehydratase | 1.594 | 3.54E-12 |
| PA4847 | *accB* | biotin carboxyl carrier protein (BCCP) | 1.496 | 7.65E-04 |
| PA4848 | *accC* | biotin carboxylase | 1.338 | 2.06E-03 |
| PA4854 | *purH* | phosphoribosylaminoimidazolecarboxamide formyltransferase | 1.323 | 5.81E-06 |
| PA4855 | *purD* | phosphoribosylamine--glycine ligase | 1.386 | 6.25E-04 |
| PA4865 | *ureA* | urease gamma subunit | 1.4 | 1.03E-02 |
| PA4866 | putative phosphinothricin acetyltransferase | putative phosphinothricin acetyltransferase | 2.15 | 3.05E-05 |

| Gene | Name | Product | $log_2$ FC | Adjusted p-value |
|---|---|---|---|---|
| PA4867 | *ureB* | urease beta subunit | 2.105 | 7.91E-06 |
| PA4868 | *ureC* | urease alpha subunit | 1.681 | 1.62E-04 |
| PA4898 | *opdK* | NA | -2.269 | 3.45E-11 |
| PA4902 | probable transcriptional regulator | NA | -1.906 | 3.62E-12 |
| PA4903 | probable major facilitator superfamily (MFS) transporter | NA | -2.028 | 4.01E-05 |
| PA4904 | *vanA* | vanillate O-demethylase oxygenase subunit | -1.332 | 4.16E-03 |
| PA4910 | branched chain amino acid ABC transporter ATP binding protein | branched chain amino acid ABC transporter ATP binding protein | -1.46 | 5.09E-06 |
| PA4911 | probable permease of ABC branched-chain amino acid transporter | probable permease of ABC branched-chain amino acid transporter | -1.599 | 5.96E-05 |
| PA4913 | probable binding protein component of ABC transporter | probable binding protein component of ABC transporter | -2.196 | 5.72E-13 |
| PA4916 | *nrtR* | NA | -1.314 | 9.98E-03 |
| PA4965 | hypothetical protein | NA | 1.339 | 4.37E-04 |
| PA4966 | hypothetical protein | NA | 1.681 | 1.69E-04 |
| PA4979 | probable acyl-CoA dehydrogenase | NA | -1.766 | 4.13E-05 |
| PA4980 | probable enoyl-CoA hydratase/isomerase | probable enoyl-CoA hydratase/isomerase | -1.961 | 1.52E-05 |
| PA5002 | *dnpA* | NA | 1.364 | 2.85E-05 |
| PA5027 | hypothetical protein | NA | -1.854 | 9.15E-04 |
| PA5071 | conserved hypothetical protein | NA | 1.418 | 1.02E-08 |

| Gene | Name | Product | $log_2$ FC | Adjusted p-value |
|---|---|---|---|---|
| PA5095 | probable permease of ABC transporter | probable permease of ABC transporter | -1.471 | 5.82E-07 |
| PA5096 | probable binding protein component of ABC transporter | probable binding protein component of ABC transporter | -1.556 | 5.48E-08 |
| PA5097 | probable amino acid permease | NA | -1.676 | 1.64E-09 |
| PA5098 | *hutH* | histidine ammonia-lyase | -2.104 | 4.51E-14 |
| PA5099 | probable transporter | NA | -3.097 | 1.32E-17 |
| PA5100 | *hutU* | urocanase | -3.415 | 4.89E-35 |
| PA5104 | conserved hypothetical protein | NA | -2.011 | 5.92E-07 |
| PA5105 | *hutC* | NA | -2.492 | 5.50E-22 |
| PA5106 | conserved hypothetical protein | conserved hypothetical protein | -3.476 | 1.51E-42 |
| PA5112 | *estA* | NA | -1.636 | 2.83E-06 |
| PA5117 | *typA* | NA | 1.371 | 1.08E-06 |
| PA5153 | amino acid (lysine/arginine/ ornithine/histidin e/octopine) ABC transporter periplasmic binding protein | NA | -1.563 | 4.31E-06 |
| PA5183.1 | *rsmN* | NA | -1.419 | 1.20E-03 |
| PA5220 | hypothetical protein | NA | 1.469 | 2.27E-04 |
| PA5221 | probable FAD-dependent monooxygenase | probable FAD-dependent monooxygenase | 1.44 | 1.16E-03 |
| PA5232 | conserved hypothetical protein | NA | -1.681 | 1.88E-03 |
| PA5264 | hypothetical protein | NA | -1.417 | 5.91E-07 |
| PA5275 | conserved hypothetical protein | NA | -1.431 | 1.40E-05 |
| PA5354 | *glcE* | glycolate oxidase subunit GlcE | -1.658 | 3.67E-02 |
| PA5356 | *glcC* | NA | -1.377 | 2.09E-04 |
| PA5400 | probable electron transfer | NA | -1.682 | 4.62E-02 |

| Gene | Name | Product | $\log_2$ FC | Adjusted p-value |
|---|---|---|---|---|
| | flavoprotein alpha subunit | | | |
| PA5401 | hypothetical protein | NA | -1.954 | 8.01E-05 |
| PA5475 | hypothetical protein | NA | -1.817 | 4.22E-05 |
| PA5506 | hypothetical protein | NA | -1.338 | 5.23E-03 |
| PA5507 | hypothetical protein | NA | -1.662 | 4.92E-05 |
| PA5568 | conserved hypothetical protein | conserved hypothetical protein | 1.498 | 3.00E-12 |

### *Supplementary Dataset C.*

**List of transcripts with significant differences in abundance in PAO1$_{MW}$ poly-cultures (+) *versus* Δ*lasR*::Tc$^R$ mutant poly-cultures (-)**

| Gene | Name | Product | $\log_2$ FC | Adjusted p-value |
|---|---|---|---|---|
| PA0039 | hypothetical protein | NA | -1.071 | 3.82E-03 |
| PA0046 | hypothetical protein | NA | -1.022 | 3.98E-05 |
| PA0085 | *hcp1* | Hcp1 | -1.367 | 7.35E-03 |
| PA0119 | probable dicarboxylate transporter | probable dicarboxylate transporter | -1.279 | 1.63E-02 |
| PA0136 | probable ATP-binding component of ABC transporter | NA | 1.232 | 1.31E-02 |
| PA0155 | *pcaR* | NA | 1.412 | 1.07E-05 |
| PA0160 | hypothetical protein | NA | -2.585 | 7.99E-07 |
| PA0161 | hypothetical protein | NA | -2.048 | 6.36E-12 |
| PA0162 | *opdC* | NA | -1.518 | 6.27E-06 |
| PA0169 | *siaD* | SiaD | -1.260 | 8.23E-06 |
| PA0170 | *siaC* | NA | -1.050 | 1.20E-03 |
| PA0171 | *siaB* | NA | -1.493 | 7.29E-08 |
| PA0173 | probable methylesterase | probable methylesterase | 1.478 | 1.76E-02 |
| PA0227 | probable CoA transferase | probable CoA transferase | 1.483 | 1.97E-02 |
| PA0359 | hypothetical protein | NA | 1.003 | 1.33E-05 |

| Gene | Name | Product | $log_2$ FC | Adjusted p-value |
|---|---|---|---|---|
| PA0451 | conserved hypothetical protein | NA | 1.103 | 2.58E-02 |
| PA0503 | probable biotin synthesis protein BioC | probable biotin synthesis protein BioC | 1.715 | 2.33E-07 |
| PA0506 | probable acyl-CoA dehydrogenase | NA | 1.125 | 7.01E-03 |
| PA0529 | conserved hypothetical protein | NA | -1.062 | 4.14E-04 |
| PA0532 | hypothetical protein | NA | -1.278 | 3.02E-04 |
| PA0535 | transcriptional regulator | NA | 1.636 | 1.97E-03 |
| PA0563 | conserved hypothetical protein | NA | -1.118 | 2.67E-07 |
| PA0604 | *agtB* | AgtB | -1.081 | 6.25E-04 |
| PA0672 | *hemO* | NA | -1.225 | 3.94E-02 |
| PA0717 | hypothetical protein of bacteriophage Pf1 | NA | -1.014 | 1.55E-02 |
| PA0826 | hypothetical protein | NA | -1.227 | 4.68E-08 |
| PA0839 | probable transcriptional regulator | NA | -1.052 | 9.67E-05 |
| PA0859 | hypothetical protein | NA | 1.171 | 9.67E-05 |
| PA0874 | hypothetical protein | NA | -2.284 | 6.27E-06 |
| PA0887 | *acsA* | acetyl-coenzyme A synthetase | -1.533 | 5.22E-03 |
| PA0913 | *mgtE* | NA | -1.063 | 3.80E-03 |
| PA0922 | hypothetical protein | NA | -1.197 | 4.59E-07 |
| PA0978 | conserved hypothetical protein | NA | -1.755 | 8.05E-06 |
| PA1051 | probable transporter | NA | -1.012 | 7.69E-03 |
| PA1159 | probable cold-shock protein | NA | -1.041 | 2.55E-05 |
| PA1183 | *dctA* | C4-dicarboxylate transport protein | -1.131 | 5.15E-04 |

| Gene | Name | Product | $log_2$ FC | Adjusted p-value |
|---|---|---|---|---|
| PA1235 | probable transcriptional regulator | NA | 1.246 | 3.00E-02 |
| PA1239 | hypothetical protein | NA | 1.618 | 2.11E-05 |
| PA1240 | probable enoyl-CoA hydratase/isomerase | NA | 1.567 | 9.22E-04 |
| PA1250 | aprI | NA | 1.190 | 2.08E-03 |
| PA1251 | probable chemotaxis transducer | NA | 1.048 | 1.44E-02 |
| PA1317 | *cyoA* | cytochrome o ubiquinol oxidase subunit II | -1.917 | 5.83E-06 |
| PA1318 | *cyoB* | cytochrome o ubiquinol oxidase subunit I | -1.758 | 2.12E-05 |
| PA1319 | *cyoC* | cytochrome o ubiquinol oxidase subunit III | -1.660 | 2.33E-04 |
| PA1320 | *cyoD* | cytochrome o ubiquinol oxidase subunit IV | -1.683 | 4.03E-04 |
| PA1321 | *cyoE* | cytochrome o ubiquinol oxidase protein CyoE | -1.230 | 1.52E-02 |
| PA1342 | *aatJ* | putative acidic amino acid ABC transporter substrate-binding protein | -1.072 | 1.86E-03 |
| PA1369 | hypothetical protein | NA | -1.126 | 4.91E-07 |
| PA1413 | probable transcriptional regulator | NA | 1.111 | 6.65E-04 |
| PA1430 | *lasR* | transcriptional regulator LasR | 5.295 | 5.61E-90 |
| PA1431 | *rsaL* | NA | 1.493 | 1.37E-03 |
| PA1432 | *lasI* | autoinducer synthesis protein LasI | 3.607 | 4.33E-17 |
| PA1433 | conserved hypothetical protein | NA | 2.338 | 1.27E-09 |
| PA1445 | *fliO* | flagellar protein FliO | 1.039 | 2.73E-03 |

| Gene | Name | Product | $log_2$ FC | Adjusted p-value |
|---|---|---|---|---|
| PA1494 | *muiA* | NA | 1.289 | 2.61E-03 |
| PA1539 | hypothetical protein | NA | -1.239 | 6.33E-04 |
| PA1540 | conserved hypothetical protein | NA | -1.048 | 1.34E-03 |
| PA1541 | probable drug efflux transporter | NA | -1.729 | 1.58E-09 |
| PA1545 | hypothetical protein | NA | -1.180 | 5.00E-03 |
| PA1601 | probable aldehyde dehydrogenase | NA | 1.031 | 1.93E-02 |
| PA1610 | *fabA* | beta-hydroxydecanoyl-ACP dehydrase | -1.412 | 7.10E-13 |
| PA1626 | probable major facilitator superfamily (MFS) transporter | NA | 1.165 | 1.85E-03 |
| PA1713 | *exsA* | transcriptional regulator ExsA | -1.198 | 1.79E-03 |
| PA1785 | *nasT* | NA | 1.035 | 2.95E-02 |
| PA1835 | hypothetical protein | NA | 1.082 | 9.91E-07 |
| PA1856 | probable cytochrome oxidase subunit | probable cytochrome oxidase subunit | -1.012 | 2.33E-04 |
| PA1884 | transcriptional regulator | NA | 1.396 | 1.92E-02 |
| PA1885 | conserved hypothetical protein | NA | 1.314 | 6.28E-03 |
| PA1942 | hypothetical protein | NA | 1.032 | 2.02E-05 |
| PA1947 | *rbsA* | ribose transport protein RbsA | 1.091 | 8.06E-04 |
| PA1970 | hypothetical protein | NA | 1.195 | 7.44E-09 |
| PA1982 | *exaA* | quinoprotein ethanol dehydrogenase | -1.917 | 4.79E-03 |
| PA1983 | *exaB* | NA | -1.872 | 3.53E-02 |
| PA2109 | hypothetical protein | NA | 1.833 | 1.85E-02 |
| PA2217 | probable aldehyde dehydrogenase | probable aldehyde dehydrogenase | 1.044 | 3.14E-03 |

| Gene | Name | Product | $log_2$ FC | Adjusted p-value |
|---|---|---|---|---|
| PA2268 | hypothetical protein | NA | 1.995 | 4.52E-04 |
| PA2277 | *arsR* | NA | 1.246 | 6.85E-03 |
| PA2302 | *ambE* | NA | 1.293 | 1.62E-04 |
| PA2303 | *ambD* | NA | 2.227 | 1.09E-04 |
| PA2304 | *ambC* | NA | 1.452 | 6.85E-03 |
| PA2305 | *ambB* | NA | 1.464 | 5.12E-04 |
| PA2355 | probable FMNH2-dependent monooxygenase | NA | 1.083 | 2.91E-02 |
| PA2386 | *pvdA* | NA | -2.763 | 1.80E-02 |
| PA2392 | *pvdP* | NA | -1.320 | 1.85E-02 |
| PA2393 | putative dipeptidase | NA | -1.873 | 3.89E-03 |
| PA2394 | *pvdN* | NA | -1.476 | 1.80E-02 |
| PA2396 | *pvdF* | NA | -1.456 | 3.38E-03 |
| PA2398 | *fpvA* | NA | -1.717 | 1.86E-03 |
| PA2402 | *pvdI* | NA | -1.106 | 4.19E-02 |
| PA2412 | conserved hypothetical protein | conserved hypothetical protein | -3.471 | 2.25E-02 |
| PA2425 | *pvdG* | NA | -1.616 | 3.88E-02 |
| PA2426 | *pvdS* | NA | -2.459 | 1.45E-03 |
| PA2462 | hypothetical protein | NA | -1.037 | 4.80E-04 |
| PA2478 | *dsbD* | NA | 1.027 | 6.01E-03 |
| PA2485 | hypothetical protein | NA | 1.004 | 5.03E-04 |
| PA2487 | hypothetical protein | NA | 1.196 | 1.25E-05 |
| PA2488 | probable transcriptional regulator | NA | 1.085 | 2.19E-04 |
| PA2521 | *czcB* | NA | 1.614 | 2.86E-02 |
| PA2539 | conserved hypothetical protein | NA | -1.131 | 3.50E-03 |
| PA2540 | conserved hypothetical protein | NA | -1.022 | 1.72E-04 |
| PA2570 | *lecA* | LecA | -1.548 | 1.24E-06 |
| PA2587 | *pqsH* | probable FAD-dependent monooxygenase | 1.881 | 2.31E-06 |
| PA2605 | conserved hypothetical protein | conserved hypothetical protein | 1.212 | 2.84E-02 |

| Gene | Name | Product | $log_2$ FC | Adjusted p-value |
|---|---|---|---|---|
| PA2667 | *mvaU* | NA | -1.137 | 1.07E-05 |
| PA2668 | hypothetical protein | NA | -1.108 | 3.98E-05 |
| PA2716 | probable FMN oxidoreductase | NA | 1.031 | 1.14E-02 |
| PA2729 | hypothetical protein | NA | -1.056 | 1.44E-04 |
| PA2730 | hypothetical protein | NA | -1.021 | 1.75E-06 |
| PA2731 | Uncharacterized protein | NA | -1.053 | 1.42E-04 |
| PA2756 | hypothetical protein | NA | -1.216 | 4.33E-04 |
| PA2762 | hypothetical protein | NA | -1.076 | 8.58E-04 |
| PA2819 | hypothetical protein | NA | -1.961 | 3.01E-06 |
| PA2825 | *ospR* | NA | 1.224 | 1.29E-02 |
| PA2880 | hypothetical protein | NA | -1.060 | 5.07E-04 |
| PA2891 | *atuF* | geranyl-CoA carboxylase | 1.270 | 1.78E-05 |
| PA2903 | *cobJ* | precorrin-3 methylase CobJ | 1.088 | 3.36E-02 |
| PA2904 | *cobI* | precorrin-2 methyltransferase CobI | 1.188 | 1.97E-02 |
| PA2910 | conserved hypothetical protein | NA | -1.571 | 1.96E-04 |
| PA2933 | probable major facilitator superfamily (MFS) transporter | NA | 1.671 | 1.90E-02 |
| PA2935 | hypothetical protein | NA | 1.477 | 2.86E-02 |
| PA2937 | hypothetical protein | NA | 1.086 | 2.31E-02 |
| PA2947 | *cobE* | CobE | 1.343 | 2.77E-02 |
| PA3035 | probable glutathione S-transferase | probable glutathione S-transferase | 1.579 | 5.51E-04 |
| PA3037 | hypothetical protein | NA | 1.020 | 9.45E-04 |
| PA3063 | *pelB* | PelB | -1.150 | 1.29E-02 |
| PA3067 | probable transcriptional regulator | NA | 1.081 | 6.61E-03 |

| Gene | Name | Product | $log_2$ FC | Adjusted p-value |
|---|---|---|---|---|
| PA3132 | probable hydrolase | NA | 1.309 | 3.55E-05 |
| PA3133 | *sawR* | NA | 1.182 | 9.67E-05 |
| PA3140 | hypothetical protein | NA | -1.156 | 4.90E-09 |
| PA3143 | transposase | NA | -1.091 | 3.18E-08 |
| PA3145 | *wbpL* | NA | -1.134 | 4.54E-09 |
| PA3146 | *wbpK* | NA | -1.075 | 6.97E-07 |
| PA3147 | *wbpJ* | NA | -1.096 | 4.41E-06 |
| PA3148 | *wbpI* | UDP-N-acetylglucosamine 2-epimerase WbpI | -1.086 | 2.42E-05 |
| PA3149 | *wbpH* | NA | -1.042 | 6.77E-04 |
| PA3181 | 2-keto-3-deoxy-6-phosphogluconate aldolase | 2-keto-3-deoxy-6-phosphogluconate aldolase | 1.404 | 2.30E-03 |
| PA3182 | *pgl* | 6-phosphogluconolactonase | 1.711 | 8.06E-04 |
| PA3193 | *glk* | glucokinase | 1.146 | 1.43E-03 |
| PA3291 | *tli1* | NA | -1.013 | 1.60E-02 |
| PA3309 | conserved hypothetical protein | NA | 1.038 | 2.14E-02 |
| PA3314 | probable ATP-binding component of ABC transporter | probable ATP-binding component of ABC transporter | 1.265 | 5.83E-03 |
| PA3321 | probable transcriptional regulator | NA | 1.421 | 2.24E-03 |
| PA3410 | *hasI* | NA | -1.373 | 2.92E-04 |
| PA3415 | probable dihydrolipoamide acetyltransferase | probable dihydrolipoamide acetyltransferase | 1.130 | 4.95E-02 |
| PA3436 | hypothetical protein | NA | 2.278 | 1.92E-17 |
| PA3476 | *rhlI* | autoinducer synthesis protein RhlI | 1.553 | 2.46E-05 |
| PA3493 | conserved hypothetical protein | NA | 1.082 | 3.19E-02 |
| PA3530 | *bfd* | NA | -1.245 | 2.08E-03 |
| PA3575 | hypothetical protein | NA | 1.059 | 1.04E-03 |

| Gene | Name | Product | $log_2$ FC | Adjusted p-value |
|---|---|---|---|---|
| PA3677 | *mexJ* | NA | 1.370 | 5.45E-09 |
| PA3681 | hypothetical protein | NA | 1.001 | 2.08E-03 |
| PA3718 | probable major facilitator superfamily (MFS) transporter | NA | 1.251 | 2.51E-07 |
| PA3719 | *armR* | antirepressor for MexR | 2.271 | 2.14E-09 |
| PA3720 | hypothetical protein | NA | 1.385 | 4.37E-09 |
| PA3727 | hypothetical protein | NA | -1.056 | 2.29E-02 |
| PA3729 | conserved hypothetical protein | NA | -1.270 | 4.03E-06 |
| PA3730 | hypothetical protein | NA | -1.019 | 5.15E-03 |
| PA3765 | hypothetical protein | NA | 1.344 | 2.03E-02 |
| PA3862 | *dauB* | NAD(P)H-dependent anabolic L-arginine dehydrogenase | 1.670 | 1.24E-06 |
| PA3863 | *dauA* | FAD-dependent catabolic D-arginine dehydrogenase | 1.672 | 1.35E-08 |
| PA3864 | *dauR* | NA | 1.139 | 9.67E-05 |
| PA3865 | putative periplasmic lysine- | putative periplasmic lysine- | 1.817 | 8.18E-09 |
| PA3905 | *tecT* | NA | 1.967 | 2.09E-04 |
| PA3907 | *tseT* | NA | 1.308 | 3.19E-02 |
| PA3908 | *tsiT* | NA | 1.661 | 9.22E-04 |
| PA3925 | probable acyl-CoA thiolase | probable acyl-CoA thiolase | 1.254 | 1.80E-08 |
| PA3928 | hypothetical protein | NA | 1.079 | 3.06E-02 |
| PA3933 | *betT3* | NA | -1.064 | 1.63E-02 |
| PA4033 | *mucE* | NA | -1.053 | 9.88E-04 |
| PA4139 | hypothetical protein | NA | -1.065 | 3.26E-02 |
| PA4181 | hypothetical protein | NA | 2.365 | 6.36E-12 |
| PA4182 | hypothetical protein | NA | 2.170 | 2.87E-16 |

| Gene | Name | Product | $log_2$ FC | Adjusted p-value |
|---|---|---|---|---|
| PA4183 | hypothetical protein | NA | 1.026 | 5.36E-03 |
| PA4197 | *bfiS* | NA | 1.117 | 1.28E-02 |
| PA4202 | *nmoA* | nitronate monooxygenase NmoA | 1.515 | 8.91E-10 |
| PA4203 | *nmoR* | NA | 1.334 | 1.95E-03 |
| PA4219 | *ampO* | NA | -5.000 | 1.71E-02 |
| PA4229 | *pchC* | NA | -3.392 | 2.54E-04 |
| PA4230 | *pchB* | salicylate biosynthesis protein PchB | -3.801 | 1.53E-02 |
| PA4231 | *pchA* | salicylate biosynthesis isochorismate synthase | -1.387 | 1.75E-02 |
| PA4357 | conserved hypothetical protein | NA | 1.538 | 5.70E-03 |
| PA4362 | hypothetical protein | NA | 1.066 | 4.86E-02 |
| PA4363 | *iciA* | NA | 1.414 | 4.58E-03 |
| PA4364 | hypothetical protein | NA | 3.270 | 1.03E-08 |
| PA4365 | *lysE* | NA | 2.440 | 3.81E-08 |
| PA4467 | hypothetical protein | NA | -1.021 | 3.01E-03 |
| PA4468 | *sodM* | NA | -1.138 | 2.10E-02 |
| PA4495 | hypothetical protein | NA | 1.451 | 8.91E-10 |
| PA4581 | *rtcR* | NA | 1.040 | 1.56E-05 |
| PA4623 | hypothetical protein | NA | 1.324 | 1.96E-05 |
| PA4746 | conserved hypothetical protein | NA | -1.074 | 9.17E-07 |
| PA4829 | *lpd3* | dihydrolipoamide dehydrogenase 3 | 1.364 | 1.64E-02 |
| PA4830 | hypothetical protein | NA | 1.210 | 3.97E-02 |
| PA4832 | probable short-chain dehydrogenase | NA | 1.195 | 1.44E-02 |
| PA4846 | *aroQ1* | 3-dehydroquinate dehydratase | -1.057 | 1.24E-06 |

| Gene | Name | Product | $log_2$ FC | Adjusted p-value |
|---|---|---|---|---|
| PA4859 | probable permease of ABC transporter | probable permease of ABC transporter | 1.099 | 1.16E-02 |
| PA4988 | *waaA* | 3-deoxy-D-manno-octulosonic-acid (KDO) transferase | 1.195 | 1.86E-03 |
| PA5020 | probable acyl-CoA dehydrogenase | probable acyl-CoA dehydrogenase | -1.065 | 4.52E-04 |
| PA5042 | *pilO* | NA | -1.133 | 1.27E-03 |
| PA5092 | *hutI* | imidazolone-5-propionate hydrolase HutI | 1.008 | 6.06E-03 |
| PA5098 | *hutH* | histidine ammonia-lyase | 1.299 | 9.22E-04 |
| PA5125 | *ntrC* | two-component response regulator NtrC | 1.123 | 2.14E-09 |
| PA5157 | probable transcriptional regulator | NA | 1.146 | 1.37E-03 |
| PA5158 | probable outer membrane protein precursor | NA | 1.186 | 3.02E-03 |
| PA5291 | *betT2* | NA | -1.337 | 3.42E-05 |
| PA5374 | *betI* | NA | -1.193 | 2.09E-04 |
| PA5375 | *betT1* | NA | -1.094 | 2.03E-07 |
| PA5408 | hypothetical protein | NA | 1.342 | 2.71E-02 |
| PA5553 | *atpC* | ATP synthase epsilon chain | -1.053 | 4.16E-02 |

### *Supplementary Dataset D.*

**List of transcripts with significant differences in abundance in PAO1$_{MW}$ mono-cultures (+) *versus* *mexT*$^{I129F}$ mutant mono-cultures (-)**

| Gene | Name | Product | $log_2$ FC | Adjusted p-value |
|---|---|---|---|---|
| PA0029 | probable sulfate transporter | NA | -1.124 | 3.08E-03 |
| PA0056 | *osaR* | NA | -1.147 | 5.67E-03 |
| PA0057 | hypothetical protein | NA | -1.143 | 1.63E-02 |
| PA0058 | *dsbM* | NA | -2.419 | 2.04e-06 |
| PA0070 | *tagQ1* | NA | 1.042 | 2.14E-03 |
| PA0072 | *tagS1* | NA | -1.888 | 3.77E-03 |

| Gene | Name | Product | $log_2$ FC | Adjusted p-value |
|---|---|---|---|---|
| PA0073 | *tagT1* | NA | -3.006 | 4.51E-03 |
| PA0112 | hypothetical protein | hypothetical protein | -1.383 | 7.45E-03 |
| PA0121 | hypothetical protein | NA | 1.098 | 4.79E-03 |
| PA0137 | probable permease of ABC transporter | NA | -1.229 | 2.00E-03 |
| PA0144 | hypothetical protein | NA | -1.240 | 7.79E-04 |
| PA0149 | probable sigma-70 factor | NA | -1.622 | 1.02E-03 |
| PA0152 | *pcaQ* | NA | -1.116 | 4.43E-04 |
| PA0157 | *triB* | NA | -1.158 | 3.43E-04 |
| PA0163 | probable transcriptional regulator | NA | -1.683 | 1.13E-04 |
| PA0166 | probable transporter | NA | -1.075 | 4.90E-02 |
| PA0168 | conserved hypothetical protein | NA | -1.849 | 2.28E-04 |
| PA0171 | *siaB* | NA | -1.379 | 1.37E-04 |
| PA0172 | *siaA* | NA | -1.753 | 3.15e-08 |
| PA0179 | probable two-component response regulator | probable two-component response regulator | 1.391 | 4.11E-04 |
| PA0182 | probable short-chain dehydrogenase | probable short-chain dehydrogenase | -3.359 | 1.15e-10 |
| PA0183 | *atsA* | arylsulfatase | -1.074 | 4.03E-02 |
| PA0188 | hypothetical protein | NA | -1.643 | 2.03E-03 |
| PA0209 | conserved hypothetical protein | NA | -1.713 | 2.58E-02 |
| PA0210 | *mdcC* | NA | -2.291 | 4.78E-02 |
| PA0219 | probable aldehyde dehydrogenase | probable aldehyde dehydrogenase | -1.174 | 4.98E-02 |
| PA0220 | amino acid APC family transporter | NA | -1.296 | 1.24E-02 |
| PA0227 | probable CoA transferase | probable CoA transferase | -1.160 | 1.25E-02 |
| PA0233 | probable transcriptional regulator | NA | -1.005 | 5.41E-03 |
| PA0237 | probable oxidoreductase | NA | -1.064 | 2.22E-02 |
| PA0238 | hypothetical protein | NA | -1.520 | 7.89E-03 |

| Gene | Name | Product | $log_2$ FC | Adjusted p-value |
|---|---|---|---|---|
| PA0255 | conserved hypothetical protein | NA | -1.069 | 2.78E-03 |
| PA0263 | *hcpC* | secreted protein Hcp | -3.817 | 5.21e-09 |
| PA0274 | hypothetical protein | NA | -1.353 | 1.00E-02 |
| PA0278 | hypothetical protein | NA | -1.090 | 8.07E-03 |
| PA0282 | *cysT* | sulfate transport protein CysT | -1.443 | 5.31E-04 |
| PA0283 | *sbp* | sulfate-binding protein precursor | -1.935 | 4.03E-04 |
| PA0284 | hypothetical protein | NA | -1.485 | 9.40E-03 |
| PA0349 | hypothetical protein | NA | -1.247 | 2.79E-02 |
| PA0369 | Uncharacterized protein | NA | -1.228 | 2.28E-03 |
| PA0380 | conserved hypothetical protein | conserved hypothetical protein | -2.060 | 5.36e-09 |
| PA0388 | hypothetical protein | NA | 1.430 | 1.11E-04 |
| PA0392 | conserved hypothetical protein | NA | 1.046 | 1.88E-03 |
| PA0409 | *pilH* | twitching motility protein PilH | 1.013 | 3.93E-02 |
| PA0411 | *pilJ* | twitching motility protein PilJ | 1.117 | 1.06E-02 |
| PA0424 | *mexR* | multidrug resistance operon repressor MexR | 1.011 | 2.61E-03 |
| PA0451 | conserved hypothetical protein | NA | -1.268 | 6.66E-03 |
| PA0471 | *fiuR* | NA | -1.283 | 1.91E-02 |
| PA0472 | *fiuI* | NA | -1.058 | 4.91E-02 |
| PA0480 | probable hydrolase | probable hydrolase | -1.055 | 2.56E-02 |
| PA0491 | probable transcriptional regulator | NA | 1.329 | 2.23E-04 |
| PA0500 | *bioB* | biotin synthase | 1.334 | 2.84E-04 |
| PA0506 | probable acyl-CoA dehydrogenase | NA | 1.478 | 1.75e-05 |
| PA0534 | pauB1 | NA | 1.156 | 1.63E-02 |

| Gene | Name | Product | $log_2$ FC | Adjusted p-value |
|---|---|---|---|---|
| PA0535 | transcriptional regulator | NA | 1.236 | 4.61E-03 |
| PA0545 | hypothetical protein | NA | 1.467 | 1.16E-04 |
| PA0563 | conserved hypothetical protein | NA | 1.038 | 2.09E-03 |
| PA0582 | *folB* | dihydroneopterin aldolase | -1.232 | 1.56E-03 |
| PA0586 | conserved hypothetical protein | NA | 1.098 | 7.52E-03 |
| PA0587 | conserved hypothetical protein | NA | 1.214 | 1.04E-02 |
| PA0596 | hypothetical protein | hypothetical protein | 1.122 | 3.61E-04 |
| PA0603 | *agtA* | AgtA | -1.678 | 1.26e-05 |
| PA0614 | hypothetical protein | NA | 1.105 | 1.85E-02 |
| PA0629 | conserved hypothetical protein | NA | -1.408 | 3.42E-03 |
| PA0630 | hypothetical protein | NA | -1.487 | 4.74E-03 |
| PA0633 | hypothetical protein | NA | 1.107 | 1.24E-03 |
| PA0634 | hypothetical protein | NA | 1.160 | 5.54E-03 |
| PA0670 | hypothetical protein | NA | -1.533 | 6.29e-06 |
| PA0671 | hypothetical protein | NA | -1.942 | 4.33e-08 |
| PA0674 | *vreA* | NA | -2.083 | 7.67E-04 |
| PA0675 | *vreI* | NA | -1.240 | 3.72E-02 |
| PA0677 | *hxcW* | HxcW | -1.372 | 2.91E-02 |
| PA0678 | *hxcU* | HxcU | -2.516 | 3.14E-02 |
| PA0683 | *hxcY* | HxcY | -2.028 | 1.50E-02 |
| PA0684 | *hxcZ* | HxcZ | -2.142 | 5.21E-03 |
| PA0692 | *pdtB* | NA | -1.127 | 3.85E-02 |
| PA0715 | hypothetical protein | NA | -2.705 | 4.49E-02 |
| PA0716.1 | *pf4r* | NA | -2.244 | 2.08E-02 |
| PA0716.2 | *xisF4* | NA | -10.493 | 1.26e-189 |
| PA0717 | hypothetical protein of bacteriophage Pf1 | NA | -11.006 | 1.13e-155 |
| PA0718 | hypothetical protein of bacteriophage Pf1 | NA | -11.098 | 8.21e-138 |

| Gene | Name | Product | $log_2$ FC | Adjusted p-value |
|---|---|---|---|---|
| PA0719 | hypothetical protein of bacteriophage Pf1 | NA | -11.307 | 1.09e-188 |
| PA0720 | helix destabilizing protein of bacteriophage Pf1 | NA | -12.100 | 1.35e-257 |
| PA0721 | *pfsE* | NA | -10.795 | 3.07e-135 |
| PA0724 | probable coat protein A of bacteriophage Pf1 | NA | -9.063 | 7.6e-185 |
| PA0726 | hypothetical protein of bacteriophage Pf1 | NA | -8.253 | 6.71e-152 |
| PA0727 | Pf replication initiator protein | NA | -8.548 | 5.9e-133 |
| PA0728 | probable bacteriophage integrase | NA | -7.460 | 7.91e-104 |
| PA0729 | *pfiT* | NA | -3.233 | 8.06E-03 |
| PA0758 | hypothetical protein | NA | 1.267 | 2.95e-05 |
| PA0777 | hypothetical protein | NA | -2.167 | 3.37e-06 |
| PA0779 | *asrA* | NA | 1.443 | 3.91E-04 |
| PA0780 | *pruR* | NA | -1.463 | 1.57e-06 |
| PA0790 | hypothetical protein | NA | -1.315 | 1.01E-02 |
| PA0806 | hypothetical protein | NA | -1.762 | 3.12E-04 |
| PA0809 | probable transporter | NA | -1.031 | 2.82E-03 |
| PA0813 | hypothetical protein | NA | -1.443 | 4.46E-04 |
| PA0828 | probable transcriptional regulator | NA | -1.456 | 8.67E-04 |
| PA0830 | hypothetical protein | NA | 1.019 | 5.35E-03 |
| PA0834 | conserved hypothetical protein | NA | 1.105 | 2.55E-04 |
| PA0835 | *pta* | phosphate acetyltransferas e | 1.169 | 1.09E-02 |
| PA0855 | hypothetical protein | NA | -1.046 | 1.59E-03 |
| PA0861 | *rbdA* | NA | 1.084 | 1.30E-04 |

| Gene | Name | Product | $log_2$ FC | Adjusted p-value |
|---|---|---|---|---|
| PA0865 | *hpd* | 4-hydroxyphenylpyruvate dioxygenase | 1.081 | 3.00E-02 |
| PA0870 | *phhC* | aromatic amino acid aminotransferase | 1.221 | 1.71E-04 |
| PA0871 | *phhB* | NA | 1.380 | 7.28e-05 |
| PA0872 | *phhA* | phenylalanine-4-hydroxylase | 1.306 | 7.45E-03 |
| PA0880 | probable ring-cleaving dioxygenase | NA | -1.183 | 4.49E-02 |
| PA0896 | *aruF* | subunit I of arginine N2-succinyltransferase = ornithine N2-succinyltransferase | 1.152 | 4.2e-05 |
| PA0897 | *aruG* | subunit II of arginine N2-succinyltransferase = ornithine N2-succinyltransferase | 1.151 | 6.12E-04 |
| PA0899 | *aruB* | N2-Succinylarginine dihydrolase | 1.032 | 9.74E-03 |
| PA0902 | hypothetical protein | hypothetical protein | -1.015 | 7.23E-04 |
| PA0907 | *alpA* | NA | 1.447 | 4.95E-04 |
| PA0910 | *alpD* | NA | 1.016 | 1.91E-03 |
| PA0921 | hypothetical protein | NA | -1.114 | 2.51E-03 |
| PA0952 | hypothetical protein | NA | 1.475 | 1.38E-03 |
| PA0962 | *dps* | NA | 1.548 | 7.67E-04 |
| PA0978 | conserved hypothetical protein | NA | -1.687 | 1.37E-03 |
| PA0979 | conserved hypothetical protein | NA | -1.237 | 2.68E-04 |
| PA0993 | *cupC2* | NA | -1.215 | 1.06E-02 |
| PA0996 | *pqsA* | PqsA | -1.335 | 1.53E-04 |
| PA0997 | *pqsB* | PqsB | -1.188 | 1.09E-02 |
| PA1030 | hypothetical protein | NA | 1.136 | 4.98E-03 |
| PA1043 | hypothetical protein | NA | -1.116 | 8.98E-04 |

| Gene | Name | Product | $log_2$ FC | Adjusted p-value |
|---|---|---|---|---|
| PA1058 | *shaE* | NA | -1.098 | 1.83E-02 |
| PA1070 | *braG* | branched-chain amino acid transport protein BraG | 1.582 | 4.17e-08 |
| PA1071 | *braF* | branched-chain amino acid transport protein BraF | 1.761 | 3.34e-08 |
| PA1072 | *braE* | branched-chain amino acid transport protein BraE | 1.720 | 1.93e-06 |
| PA1073 | *braD* | branched-chain amino acid transport protein BraD | 1.430 | 2.58E-04 |
| PA1074 | *braC* | branched-chain amino acid transport protein BraC | 1.300 | 8.09E-04 |
| PA1076 | hypothetical protein | NA | 1.980 | 6.21e-05 |
| PA1089 | conserved hypothetical protein | NA | 1.085 | 8.90E-03 |
| PA1095 | hypothetical protein | hypothetical protein | 1.061 | 7.67E-04 |
| PA1096 | hypothetical protein | NA | 1.101 | 3.57E-04 |
| PA1103 | probable flagellar assembly protein | probable flagellar assembly protein | 1.122 | 1.03E-02 |
| PA1108 | probable major facilitator superfamily (MFS) transporter | NA | -1.621 | 2.48E-03 |
| PA1125 | probable cobalamin biosynthetic protein | NA | -1.116 | 3.20E-04 |
| PA1146 | probable iron-containing alcohol dehydrogenase | probable iron-containing alcohol dehydrogenase | -1.734 | 9.32e-06 |
| PA1155 | *nrdB* | NrdB | -1.029 | 3.07E-03 |
| PA1156 | *nrdA* | NrdA | -1.388 | 3.21e-05 |
| PA1172 | *napC* | NA | 1.025 | 7.74E-03 |
| PA1178 | *oprH* | NA | -3.807 | 2.85e-12 |

| Gene | Name | Product | $log_2$ FC | Adjusted p-value |
|---|---|---|---|---|
| PA1179 | *phoP* | two-component response regulator PhoP | -2.108 | 4.05E-04 |
| PA1180 | *phoQ* | two-component sensor PhoQ | -1.918 | 5.58e-05 |
| PA1197 | hypothetical protein | NA | 1.069 | 1.12E-03 |
| PA1202 | probable hydrolase | NA | -1.363 | 2.98E-02 |
| PA1211 | hypothetical protein | NA | -1.216 | 4.58E-02 |
| PA1213 | hypothetical protein | NA | -1.428 | 5.28E-03 |
| PA1214 | hypothetical protein | NA | -1.455 | 1.12E-02 |
| PA1215 | hypothetical protein | NA | -1.015 | 1.47E-02 |
| PA1220 | hypothetical protein | NA | -1.634 | 2.27E-03 |
| PA1221 | nonribosomal peptide synthetase | NA | -1.901 | 7.1e-09 |
| PA1237 | probable multidrug resistance efflux pump | NA | -2.002 | 8.70E-03 |
| PA1246 | *aprD* | alkaline protease secretion protein AprD | -1.476 | 9.29e-05 |
| PA1251 | probable chemotaxis transducer | NA | -1.052 | 9.52E-03 |
| PA1265 | hypothetical protein | NA | -1.605 | 5.44E-03 |
| PA1266 | *lhpE* | NA | -1.468 | 1.26E-02 |
| PA1267 | *lhpB* | D-hydroxyproline dehydrogenase beta-subunit | -1.505 | 2.83E-02 |
| PA1275 | *cobD* | cobalamin biosynthetic protein CobD | -1.453 | 2.34E-02 |
| PA1276 | *cobC* | cobalamin biosynthetic protein CobC | -1.296 | 7.82E-03 |
| PA1281 | *cobV* | cobalamin (5'-phosphate) synthase | -1.877 | 2.10E-04 |
| PA1282 | probable major facilitator superfamily | NA | -1.131 | 4.77E-03 |

| Gene | Name | Product | $log_2$ FC | Adjusted p-value |
|---|---|---|---|---|
| | (MFS) transporter | | | |
| PA1290 | probable transcriptional regulator | NA | 1.102 | 1.90E-03 |
| PA1297 | probable metal transporter | NA | -1.046 | 1.27E-02 |
| PA1298 | conserved hypothetical protein | NA | -1.826 | 2.58E-04 |
| PA1301 | *hxuR* | NA | -1.334 | 7.28E-03 |
| PA1326 | *ilvA2* | threonine dehydratase | -2.090 | 3.74E-02 |
| PA1329 | conserved hypothetical protein | NA | -1.612 | 1.69E-02 |
| PA1331 | conserved hypothetical protein | NA | -1.241 | 1.61E-02 |
| PA1332 | hypothetical protein | NA | 1.745 | 3.18e-11 |
| PA1333 | hypothetical protein | NA | 1.360 | 7.21E-04 |
| PA1334 | probable oxidoreductase | probable oxidoreductase | -1.259 | 1.58E-04 |
| PA1340 | *aatM* | AatM | 1.274 | 4.70E-04 |
| PA1341 | *aatQ* | AatQ | 1.026 | 1.62E-02 |
| PA1342 | *aatJ* | putative acidic amino acid ABC transporter substrate-binding protein | 1.038 | 1.24E-02 |
| PA1343 | hypothetical protein | NA | -2.667 | 1.09e-05 |
| PA1344 | probable short-chain dehydrogenase | NA | -1.145 | 7.89E-03 |
| PA1351 | probable sigma-70 factor | NA | -1.191 | 1.01E-02 |
| PA1356 | hypothetical protein | NA | -1.183 | 7.89E-03 |
| PA1359 | transcriptional regulator | NA | -1.019 | 3.91E-04 |
| PA1364 | probable transmembrane sensor | NA | -1.153 | 4.61E-02 |
| PA1399 | probable transcriptional regulator | NA | -1.267 | 5.17E-04 |
| PA1400 | probable pyruvate carboxylase | NA | -1.284 | 5.66e-05 |

| Gene | Name | Product | $log_2$ FC | Adjusted p-value |
|---|---|---|---|---|
| PA1412 | hypothetical protein | NA | -1.143 | 3.93E-02 |
| PA1414 | hypothetical protein | NA | 1.204 | 1.06E-02 |
| PA1419 | probable transporter | NA | -1.049 | 1.46E-02 |
| PA1435 | probable Resistance-Nodulation-Cell Division (RND) efflux membrane fusion protein precursor | NA | -1.372 | 6.05E-04 |
| PA1443 | *fliM* | flagellar motor switch protein FliM | 1.100 | 1.00E-02 |
| PA1451 | conserved hypothetical protein | NA | -2.207 | 2.81E-02 |
| PA1456 | *cheY* | two-component response regulator CheY | 1.017 | 7.43E-03 |
| PA1457 | *cheZ* | chemotaxis protein CheZ | 1.287 | 4.02E-04 |
| PA1470 | probable short-chain dehydrogenase | probable short-chain dehydrogenase | -1.164 | 7.67E-04 |
| PA1475 | *ccmA* | heme exporter protein CcmA | -1.304 | 5.85E-04 |
| PA1476 | *ccmB* | heme exporter protein CcmB | -1.097 | 7.88E-03 |
| PA1483 | *cycH* | NA | -1.374 | 4.05E-04 |
| PA1486 | *bapF* | NA | -1.326 | 3.43E-03 |
| PA1488 | hypothetical protein | NA | -1.738 | 1.11E-03 |
| PA1495 | hypothetical protein | NA | -1.084 | 4.28E-02 |
| PA1497 | probable transporter | NA | -1.388 | 2.04E-02 |
| PA1512 | *hcpA* | secreted protein Hcp | -2.017 | 1.21E-03 |
| PA1513 | hypothetical protein | NA | 1.559 | 7.33e-05 |
| PA1514 | ureidoglycolate hydrolaseYbbT | ureidoglycolate hydrolaseYbbT | 1.898 | 1.02e-05 |
| PA1515 | *alc* | allantoicase | 1.252 | 8.86E-03 |
| PA1516 | hypothetical protein | NA | 1.259 | 1.35E-02 |
| PA1517 | conserved hypothetical protein | NA | 1.425 | 1.89E-03 |

| Gene | Name | Product | $log_2$ FC | Adjusted p-value |
|---|---|---|---|---|
| PA1518 | conserved hypothetical protein | conserved hypothetical protein | 1.210 | 7.26E-04 |
| PA1537 | probable short-chain dehydrogenase | NA | 1.228 | 1.27E-03 |
| PA1538 | probable flavin-containing monooxygenase | NA | 1.140 | 1.21E-03 |
| PA1546 | *hemN* | NA | 1.273 | 6.17E-04 |
| PA1555 | *ccoP2* | Cytochrome c oxidase | 1.021 | 7.38E-03 |
| PA1561 | *aer* | aerotaxis receptor Aer | 1.182 | 3.52E-03 |
| PA1566 | *pauA3* | Glutamylpolyamine synthetase | -1.464 | 8.93e-05 |
| PA1578 | hypothetical protein | NA | -1.103 | 3.24E-03 |
| PA1596 | *htpG* | NA | 1.473 | 1.22e-05 |
| PA1600 | probable cytochrome c | NA | 1.190 | 7.79E-04 |
| PA1602 | probable oxidoreductase | probable oxidoreductase | 1.137 | 6.28E-03 |
| PA1617 | probable AMP-binding enzyme | NA | -1.071 | 8.61E-04 |
| PA1629 | probable enoyl-CoA hydratase/isomerase | NA | 1.025 | 7.18E-03 |
| PA1630 | probable transcriptional regulator | NA | 1.002 | 2.45E-03 |
| PA1631 | probable acyl-CoA dehydrogenase | probable acyl-CoA dehydrogenase | 1.046 | 4.21E-03 |
| PA1645 | hypothetical protein | NA | -1.640 | 5.04e-05 |
| PA1655 | probable glutathione S-transferase | probable glutathione S-transferase | 1.121 | 1.45e-05 |
| PA1670 | *stp1* | Stp1 | -1.472 | 1.21E-03 |
| PA1698 | *popN* | Type III secretion outer membrane protein PopN precursor | -1.953 | 4.32E-04 |
| PA1704 | *pcrR* | NA | -1.982 | 3.28E-02 |
| PA1705 | *pcrG* | NA | -1.974 | 1.58E-02 |
| PA1742 | *pauD2* | NA | 1.212 | 2.03E-03 |
| PA1743 | hypothetical protein | NA | 1.232 | 2.85E-02 |

| Gene | Name | Product | $log_2$ FC | Adjusted p-value |
|---|---|---|---|---|
| PA1744 | hypothetical protein | NA | 2.321 | 1.8e-11 |
| PA1747 | hypothetical protein | NA | 1.469 | 8.52E-04 |
| PA1748 | probable enoyl-CoA hydratase/isomerase | probable enoyl-CoA hydratase/isomerase | 1.202 | 5.25E-04 |
| PA1761 | hypothetical protein | NA | 1.015 | 4.57E-04 |
| PA1782 | probable serine/threonine-protein kinase | NA | -1.866 | 2.47E-04 |
| PA1788 | hypothetical protein | NA | 1.320 | 9.67E-04 |
| PA1789 | hypothetical protein | NA | 1.248 | 1.82E-03 |
| PA1818 | *cadA* | lysine decarboxylase | 1.015 | 9.23e-05 |
| PA1830 | hypothetical protein | NA | 1.027 | 3.23E-03 |
| PA1839 | hypothetical protein | NA | -1.151 | 2.31E-04 |
| PA1850 | probable transcriptional regulator | NA | -1.075 | 4.19E-04 |
| PA1854 | conserved hypothetical protein | NA | -1.363 | 3.22E-03 |
| PA1855 | hypothetical protein | NA | -3.961 | 2.15e-12 |
| PA1856 | probable cytochrome oxidase subunit | probable cytochrome oxidase subunit | -2.036 | 6.05E-04 |
| PA1858 | *str* | NA | -1.054 | 2.86E-03 |
| PA1863 | *modA* | molybdate-binding periplasmic protein precursor ModA | 1.416 | 5.22e-05 |
| PA1870 | hypothetical protein | NA | -1.035 | 1.76E-02 |
| PA1879 | hypothetical protein | NA | -1.265 | 1.79E-03 |
| PA1884 | transcriptional regulator | NA | -1.194 | 4.55E-03 |
| PA1891 | hypothetical protein | NA | -2.673 | 2.18E-03 |
| PA1893 | hypothetical protein | hypothetical protein | -1.292 | 1.16E-03 |
| PA1908 | probable major facilitator superfamily | NA | -1.729 | 4.77E-03 |

| Gene | Name | Product | $log_2$ FC | Adjusted p-value |
|---|---|---|---|---|
| | (MFS) transporter | | | |
| PA1911 | *femR* | NA | -2.494 | 1.54E-03 |
| PA1918 | hypothetical protein | NA | -1.371 | 8.62E-04 |
| PA1919 | *nrdG* | NA | -1.100 | 1.29E-02 |
| PA1921 | hypothetical protein | NA | -1.373 | 2.73E-02 |
| PA1923 | hypothetical protein | hypothetical protein | -2.004 | 8.25e-05 |
| PA1929 | hypothetical protein | NA | -1.212 | 7.99E-03 |
| PA1931 | probable ferredoxin | probable ferredoxin | -1.025 | 7.82E-03 |
| PA1932 | probable hydroxylase molybdopterin-containing subunit | probable hydroxylase molybdopterin-containing subunit | -1.184 | 2.82E-04 |
| PA1941 | hypothetical protein | NA | 1.807 | 1.41e-09 |
| PA1942 | hypothetical protein | NA | 6.316 | 2.53e-70 |
| PA1948 | *rbsC* | membrane protein component of ABC ribose transporter | 1.237 | 2.62E-04 |
| PA1957 | hypothetical protein | NA | -1.287 | 1.85E-03 |
| PA1970 | hypothetical protein | NA | 3.601 | 2.56e-28 |
| PA1971 | *braZ* | NA | -1.224 | 2.43E-03 |
| PA1973 | *pqqF* | NA | -1.238 | 1.24E-03 |
| PA1977 | hypothetical protein | NA | -1.677 | 3.37E-02 |
| PA1990 | *pqqH* | NA | -1.468 | 1.07E-03 |
| PA1992 | *ercS* | NA | -1.099 | 1.58E-02 |
| PA2009 | *hmgA* | homogentisate 1 | 1.363 | 2.98E-03 |
| PA2014 | *liuB* | methylcrotonyl-CoA carboxylase | 1.061 | 3.25E-02 |
| PA2015 | *liuA* | putative isovaleryl-CoA dehydrogenase | 1.191 | 1.99E-02 |
| PA2019 | *mexX* | Resistance-Nodulation-Cell Division (RND) multidrug efflux membrane fusion protein MexX precursor | -1.715 | 1.20E-04 |

| Gene | Name | Product | $log_2$ FC | Adjusted p-value |
|---|---|---|---|---|
| PA2034 | hypothetical protein | NA | -1.298 | 1.91E-02 |
| PA2040 | *pauA4* | Glutamylpolyamine synthetase | -1.215 | 2.79E-04 |
| PA2046 | hypothetical protein | NA | -1.866 | 2.99E-04 |
| PA2047 | *cmrA* | NA | -1.088 | 9.11E-03 |
| PA2048 | hypothetical protein | NA | -1.479 | 3.85E-03 |
| PA2050 | probable sigma-70 factor | NA | -1.444 | 2.49E-02 |
| PA2062 | probable pyridoxal-phosphate dependent enzyme | probable pyridoxal-phosphate dependent enzyme | -1.557 | 2.52E-04 |
| PA2065 | *pcoA* | NA | -1.195 | 1.95E-02 |
| PA2069 | probable carbamoyl transferase | NA | -1.564 | 2.62E-04 |
| PA2082 | *kynR* | NA | 1.184 | 2.42E-04 |
| PA2084 | probable asparagine synthetase | probable asparagine synthetase | -1.008 | 3.44E-03 |
| PA2086 | probable epoxide hydrolase | probable epoxide hydrolase | -1.553 | 1.29E-03 |
| PA2088 | hypothetical protein | NA | -1.267 | 2.07E-02 |
| PA2091 | hypothetical protein | NA | -1.508 | 3.50E-02 |
| PA2092 | probable major facilitator superfamily (MFS) transporter | NA | -1.505 | 3.95E-02 |
| PA2099 | probable short-chain dehydrogenase | NA | 1.287 | 2.34E-03 |
| PA2100 | probable transcriptional regulator | NA | 1.775 | 1.09E-04 |
| PA2102 | hypothetical protein | NA | 1.295 | 7.18E-03 |
| PA2103 | probable molybdopterin biosynthesis protein MoeB | probable molybdopterin biosynthesis protein MoeB | 1.436 | 2.17E-04 |
| PA2104 | probable cysteine synthase | probable cysteine synthase | 1.296 | 3.90E-03 |

| Gene | Name | Product | $log_2$ FC | Adjusted p-value |
|---|---|---|---|---|
| PA2105 | probable acetyltransferase | probable acetyltransferase | 1.188 | 2.48E-03 |
| PA2113 | *opdO* | NA | -1.148 | 3.59E-02 |
| PA2116 | conserved hypothetical protein | NA | -1.100 | 1.52E-02 |
| PA2118 | *ada* | NA | -1.036 | 2.88E-03 |
| PA2124 | probable dehydrogenase | NA | -1.309 | 2.42E-03 |
| PA2127 | *cgrA* | NA | 1.672 | 4.01E-03 |
| PA2135 | probable transporter | NA | -1.903 | 1.80E-03 |
| PA2162 | probable glycosyl hydrolase | probable glycosyl hydrolase | -1.458 | 1.64E-02 |
| PA2163 | hypothetical protein | hypothetical protein | -2.694 | 8.34e-05 |
| PA2164 | probable glycosyl hydrolase | probable glycosyl hydrolase | -1.578 | 2.42E-04 |
| PA2172 | hypothetical protein | NA | -1.274 | 7.94E-03 |
| PA2193 | *hcnA* | hydrogen cyanide synthase HcnA | -1.622 | 5.57E-03 |
| PA2202 | probable amino acid permease | NA | -1.383 | 3.36E-04 |
| PA2203 | probable amino acid permease | NA | -1.589 | 1.34E-03 |
| PA2204 | putative binding protein component of ABC transporter | NA | -1.555 | 3.14E-03 |
| PA2207 | hypothetical protein | NA | -1.480 | 2.86E-03 |
| PA2208 | hypothetical protein | NA | -1.401 | 3.16E-02 |
| PA2211 | conserved hypothetical protein | NA | -1.168 | 4.11E-02 |
| PA2212 | conserved hypothetical protein | conserved hypothetical protein | -1.145 | 8.57E-03 |
| PA2213 | probable porin | NA | -1.172 | 1.04E-02 |
| PA2214 | putative L-lyxonate transporter | NA | -1.321 | 3.14E-03 |
| PA2238 | *pslH* | PslH | -1.241 | 1.31E-04 |
| PA2241 | *pslK* | PslL | -1.993 | 3.15e-06 |
| PA2243 | *pslM* | NA | -1.059 | 1.94E-02 |
| PA2245 | *pslO* | NA | -1.494 | 5.67E-03 |

| Gene | Name | Product | $log_2$ FC | Adjusted p-value |
|---|---|---|---|---|
| PA2247 | *bkdA1* | 2-oxoisovalerate dehydrogenase (alpha subunit) | 1.699 | 2.06E-03 |
| PA2248 | *bkdA2* | 2-oxoisovalerate dehydrogenase (beta subunit) | 1.821 | 2.62E-04 |
| PA2249 | *bkdB* | branched-chain alpha-keto acid dehydrogenase (lipoamide component) | 1.609 | 4.46E-04 |
| PA2250 | *lpdV* | lipoamide dehydrogenase -Val | 1.382 | 5.28E-04 |
| PA2252 | probable AGCS sodium/alanine/ glycine symporter | NA | -1.162 | 8.03E-04 |
| PA2253 | *ansA* | L-asparaginase I | -1.389 | 3.38E-04 |
| PA2268 | hypothetical protein | NA | -1.829 | 9.22e-06 |
| PA2270 | probable transcriptional regulator | NA | -1.026 | 2.17E-03 |
| PA2271 | probable acetyltransferase | NA | -1.019 | 8.58E-03 |
| PA2274 | hypothetical protein | NA | -1.162 | 2.26E-03 |
| PA2275 | probable alcohol dehydrogenase (Zn-dependent) | NA | -1.508 | 3.9e-07 |
| PA2283 | hypothetical protein | NA | -2.079 | 5.45e-07 |
| PA2284 | hypothetical protein | NA | -2.065 | 1.04e-07 |
| PA2285 | hypothetical protein | NA | -1.165 | 3.92E-03 |
| PA2286 | hypothetical protein | NA | -1.598 | 3.44e-06 |
| PA2287 | hypothetical protein | NA | -1.979 | 3.01e-08 |
| PA2288 | hypothetical protein | NA | -2.174 | 6.05e-10 |
| PA2292 | hypothetical protein | NA | 1.361 | 8.47E-03 |
| PA2297 | probable ferredoxin | NA | 1.012 | 1.36E-02 |
| PA2302 | *ambE* | NA | -1.136 | 1.52E-04 |

| Gene | Name | Product | $log_2$ FC | Adjusted p-value |
| --- | --- | --- | --- | --- |
| PA2305 | *ambB* | NA | -1.297 | 8.99E-04 |
| PA2307 | probable permease of ABC transporter | NA | -1.061 | 3.14E-02 |
| PA2308 | probable ATP-binding component of ABC transporter | NA | -1.585 | 4.34E-03 |
| PA2310 | hypothetical protein | hypothetical protein | -1.271 | 8.96E-03 |
| PA2311 | hypothetical protein | NA | -2.165 | 7.84E-03 |
| PA2312 | probable transcriptional regulator | NA | -1.678 | 4.82E-03 |
| PA2314 | probable major facilitator superfamily (MFS) transporter | NA | -2.106 | 4.11E-04 |
| PA2315 | hypothetical protein | NA | -1.710 | 7.52E-04 |
| PA2316 | probable transcriptional regulator | NA | -1.180 | 9.72E-03 |
| PA2323 | *gapN* | GapN | 1.126 | 1.41E-02 |
| PA2324 | hypothetical protein | NA | -1.604 | 1.73E-02 |
| PA2330 | hypothetical protein | NA | -1.239 | 2.32E-02 |
| PA2334 | probable transcriptional regulator | NA | -1.650 | 1.71E-03 |
| PA2336 | hypothetical protein | NA | -1.061 | 2.73E-02 |
| PA2342 | *mtlD* | mannitol dehydrogenase | -1.119 | 2.02E-02 |
| PA2344 | *mtlZ* | fructokinase | -1.822 | 7.67E-04 |
| PA2346 | conserved hypothetical protein | NA | -1.466 | 1.80E-03 |
| PA2347 | hypothetical protein | NA | -1.646 | 7.38E-03 |
| PA2354 | *sfnR1* | NA | -1.044 | 1.62E-02 |
| PA2357 | *msuE* | NADH-dependent FMN reductase MsuE | -2.026 | 5.63E-03 |
| PA2358 | hypothetical protein | NA | -2.263 | 7.39e-05 |
| PA2359 | probable transcriptional regulator | NA | -1.788 | 7.35e-05 |

| Gene | Name | Product | $log_2$ FC | Adjusted p-value |
|---|---|---|---|---|
| PA2369 | *hsiG3* | NA | -1.615 | 8.52e-06 |
| PA2370 | *hsiH3* | HsiH3 | -1.370 | 1.66E-02 |
| PA2375 | hypothetical protein | NA | -1.245 | 1.01E-02 |
| PA2378 | probable aldehyde dehydrogenase | NA | -1.023 | 6.70E-03 |
| PA2383 | probable transcriptional regulator | NA | -1.377 | 4.88E-04 |
| PA2385 | *pvdQ* | NA | -1.142 | 2.85E-02 |
| PA2411 | probable thioesterase | NA | -1.760 | 3.10E-02 |
| PA2437 | hypothetical protein | NA | -1.295 | 1.55E-02 |
| PA2450 | hypothetical protein | NA | -1.156 | 1.31E-03 |
| PA2452 | hypothetical protein | NA | -1.434 | 1.16E-03 |
| PA2465 | hypothetical protein | NA | -1.471 | 5.90E-03 |
| PA2467 | *foxR* | NA | -1.615 | 2.76E-03 |
| PA2477 | *dsbE* | NA | -1.055 | 1.14E-02 |
| PA2485 | hypothetical protein | NA | 2.644 | 4.77e-13 |
| PA2486 | *ptrC* | NA | 3.505 | 4.47e-28 |
| PA2487 | hypothetical protein | NA | 2.371 | 4.04e-14 |
| PA2490 | conserved hypothetical protein | NA | 1.737 | 4.47e-06 |
| PA2491 | *mexS* | NA | 3.578 | 3.15e-31 |
| PA2493 | *mexE* | NA | 5.364 | 4.09e-83 |
| PA2494 | *mexF* | NA | 5.907 | 4.97e-60 |
| PA2495 | *oprN* | NA | 5.197 | 1.23e-55 |
| PA2498 | conserved hypothetical protein | NA | -1.075 | 2.94E-02 |
| PA2501 | hypothetical protein | NA | 1.074 | 1.26E-02 |
| PA2502 | hypothetical protein | NA | -1.094 | 3.38E-04 |
| PA2521 | *czcB* | NA | -1.591 | 9.06E-04 |
| PA2522 | *czcC* | NA | -1.865 | 2.36e-05 |
| PA2523 | *czcR* | CzcR | -1.851 | 1.57e-06 |
| PA2524 | *czcS* | CzcS | -1.304 | 5.08E-04 |
| PA2525 | *opmB* | NA | -1.107 | 3.26E-03 |
| PA2526 | *muxC* | MuxC | -1.087 | 2.76E-03 |
| PA2549 | conserved hypothetical protein | NA | -2.145 | 2.76e-05 |

| Gene | Name | Product | $log_2$ FC | Adjusted p-value |
|---|---|---|---|---|
| PA2566.1 | Uncharacterized protein | NA | 1.058 | 3.09E-02 |
| PA2567 | hypothetical protein | NA | 1.142 | 2.77E-03 |
| PA2596 | conserved hypothetical protein | conserved hypothetical protein | -1.038 | 1.51E-02 |
| PA2605 | conserved hypothetical protein | conserved hypothetical protein | -1.611 | 1.05E-04 |
| PA2634 | *aceA* | isocitrate lyase AceA | 1.177 | 1.07E-03 |
| PA2638 | *nuoB* | NADH dehydrogenase I chain B | 1.081 | 1.35E-02 |
| PA2660 | hypothetical protein | NA | -1.233 | 7.42e-05 |
| PA2680 | probable quinone oxidoreductase | NA | -1.177 | 1.24E-02 |
| PA2681 | probable transcriptional regulator | NA | -1.040 | 2.86E-02 |
| PA2685 | *vgrG4* | VgrG4 | -1.284 | 7.89E-03 |
| PA2691 | conserved hypothetical protein | conserved hypothetical protein | -1.363 | 3.85E-03 |
| PA2697 | hypothetical protein | NA | -1.762 | 7.32e-05 |
| PA2698 | probable hydrolase | NA | -1.398 | 1.83E-02 |
| PA2701 | probable major facilitator superfamily (MFS) transporter | NA | -1.285 | 1.27E-03 |
| PA2714 | probable molybdopterin oxidoreductase | NA | -1.020 | 1.09E-03 |
| PA2753 | hypothetical protein | NA | 1.810 | 5.62E-04 |
| PA2756 | hypothetical protein | NA | -1.829 | 1.36E-04 |
| PA2757 | hypothetical protein | NA | -2.106 | 1.49e-06 |
| PA2758 | probable transcriptional regulator | NA | 1.882 | 4.51e-08 |
| PA2760 | *oprQ* | NA | 1.026 | 7.86E-03 |
| PA2804 | hypothetical protein | NA | -1.166 | 2.29E-02 |
| PA2805 | hypothetical protein | NA | 1.291 | 1.43E-03 |

| Gene | Name | Product | $log_2$ FC | Adjusted p-value |
|---|---|---|---|---|
| PA2811 | probable permease of ABC-2 transporter | NA | 2.602 | 2.02e-21 |
| PA2812 | probable ATP-binding component of ABC transporter | NA | 2.433 | 3.01e-18 |
| PA2813 | probable glutathione S-transferase | probable glutathione S-transferase | 2.962 | 1.13e-33 |
| PA2825 | *ospR* | NA | 1.615 | 2.29E-04 |
| PA2826 | probable glutathione peroxidase | probable glutathione peroxidase | 1.731 | 6.02e-05 |
| PA2839 | conserved hypothetical protein | NA | -1.100 | 1.72E-04 |
| PA2857 | probable ATP-binding component of ABC transporter | NA | -1.284 | 3.70E-04 |
| PA2872 | hypothetical protein | NA | -1.584 | 1.82E-03 |
| PA2874 | hypothetical protein | NA | -1.529 | 2.86E-03 |
| PA2889 | *atuD* | citronellyl-CoA dehydrogenase | 1.647 | 4.2e-05 |
| PA2898 | hypothetical protein | NA | -1.774 | 3.18E-04 |
| PA2903 | *cobJ* | precorrin-3 methylase CobJ | -1.108 | 1.95E-03 |
| PA2908 | *cbiD* | cobalamin biosynthetic protein CbiD | -1.119 | 2.82E-03 |
| PA2909 | hypothetical protein | hypothetical protein | -1.681 | 3.98E-02 |
| PA2912 | probable ATP-binding component of ABC transporter | probable ATP-binding component of ABC transporter | -1.720 | 4.00E-03 |
| PA2914 | probable permease of ABC transporter | probable permease of ABC transporter | -2.143 | 6.74e-09 |
| PA2915 | hypothetical protein | NA | -1.317 | 4.42E-04 |
| PA2919 | hypothetical protein | NA | -1.266 | 3.02E-02 |
| PA2922 | probable hydrolase | probable hydrolase | -1.233 | 7.20E-03 |
| PA2936 | hypothetical protein | NA | -1.182 | 2.91E-03 |
| PA2947 | *cobE* | CobE | -1.187 | 1.73E-02 |

| Gene | Name | Product | $log_2$ FC | Adjusted p-value |
|---|---|---|---|---|
| PA2953 | electron transfer flavoprotein-ubiquinone oxidoreductase | NA | 1.267 | 5.08e-05 |
| PA3006 | *psrA* | NA | 1.633 | 3.81e-07 |
| PA3007 | *lexA* | NA | -1.023 | 2.62E-04 |
| PA3013 | *faoB* | fatty-acid oxidation complex beta-subunit | 1.107 | 7.20E-03 |
| PA3014 | *faoA* | fatty-acid oxidation complex alpha-subunit | 1.371 | 1.95E-03 |
| PA3025 | probable FAD-dependent glycerol-3-phosphate dehydrogenase | probable FAD-dependent glycerol-3-phosphate dehydrogenase | -1.479 | 5.81E-04 |
| PA3041 | hypothetical protein | NA | 1.249 | 1.18E-04 |
| PA3060 | *pelE* | PelE | -2.051 | 7.73e-06 |
| PA3063 | *pelB* | PelB | -1.360 | 3.38E-04 |
| PA3067 | probable transcriptional regulator | NA | 1.217 | 1.85E-03 |
| PA3068 | *gdhB* | NAD-dependent glutamate dehydrogenase | 1.403 | 2.68e-07 |
| PA3095 | *xcpZ* | general secretion pathway protein M | -1.990 | 4.06E-04 |
| PA3114 | *truA* | NA | 1.046 | 6.91E-03 |
| PA3115 | *fimV* | NA | 1.066 | 2.92E-02 |
| PA3118 | *leuB* | 3-isopropylmalate dehydrogenase | 1.076 | 7.39E-04 |
| PA3120 | *leuD* | 3-isopropylmalate dehydratase small subunit | 1.335 | 1.26e-05 |
| PA3122 | probable transcriptional regulator | NA | 1.150 | 1.56E-03 |
| PA3123 | RidA subfamily protein | NA | 1.558 | 8.95E-04 |
| PA3124 | probable transcriptional regulator | NA | 1.461 | 3.75E-04 |
| PA3126 | *ibpA* | NA | 1.688 | 7.04e-05 |

| Gene | Name | Product | $\log_2$ FC | Adjusted p-value |
|---|---|---|---|---|
| PA3144 | transposase with Helix-turn-helix Hin domain | NA | -1.697 | 2.3e-05 |
| PA3183 | *zwf* | glucose-6-phosphate 1-dehydrogenase | 1.166 | 4.31E-04 |
| PA3195 | *gapA* | glyceraldehyde 3-phosphate dehydrogenase | 1.222 | 4.03E-03 |
| PA3205 | hypothetical protein | NA | -1.408 | 8.60E-03 |
| PA3206 | *cpxS* | NA | -1.583 | 1.30E-03 |
| PA3207 | hypothetical protein | NA | -1.619 | 1.37E-04 |
| PA3208 | conserved hypothetical protein | NA | -1.160 | 1.85E-03 |
| PA3226 | probable hydrolase | NA | 1.391 | 6.89E-04 |
| PA3227 | *ppiA* | peptidyl-prolyl cis-trans isomerase A | 1.772 | 6.8e-06 |
| PA3228 | probable ATP-binding/permease fusion ABC transporter | probable ATP-binding/permease fusion ABC transporter | 1.364 | 2.95e-05 |
| PA3229 | hypothetical protein | NA | 5.650 | 1.26e-81 |
| PA3230 | conserved hypothetical protein | NA | 3.816 | 1.85e-39 |
| PA3231 | hypothetical protein | NA | 3.752 | 2.41e-19 |
| PA3232 | probable nuclease | probable nuclease | 1.870 | 7.85e-08 |
| PA3233 | hypothetical protein | NA | 1.361 | 2.04E-04 |
| PA3278 | hypothetical protein | NA | 1.587 | 8.54e-05 |
| PA3309 | conserved hypothetical protein | NA | 1.150 | 3.62E-02 |
| PA3310 | conserved hypothetical protein | conserved hypothetical protein | 1.226 | 1.52E-04 |
| PA3314 | probable ATP-binding component of ABC transporter | probable ATP-binding component of ABC transporter | -1.885 | 9.19e-06 |
| PA3327 | probable non-ribosomal | NA | -1.499 | 1.59e-06 |

| Gene | Name | Product | $log_2$ FC | Adjusted p-value |
|---|---|---|---|---|
| | peptide synthetase | | | |
| PA3328 | probable FAD-dependent monooxygenase | NA | -2.033 | 5.85E-04 |
| PA3330 | probable short chain dehydrogenase | NA | -1.961 | 2.01E-03 |
| PA3331 | cytochrome P450 | NA | -1.300 | 3.44E-03 |
| PA3337 | *rfaD* | ADP-L-glycero-D-mannoheptose 6-epimerase | 1.294 | 1.46E-02 |
| PA3348 | *cheR1* | CheR1 | 1.048 | 1.24E-02 |
| PA3360 | probable secretion protein | NA | -1.324 | 6.33E-04 |
| PA3374 | conserved hypothetical protein | conserved hypothetical protein | -1.557 | 5.84e-05 |
| PA3375 | probable ATP-binding component of ABC transporter | probable ATP-binding component of ABC transporter | -1.291 | 3.28E-02 |
| PA3376 | probable ATP-binding component of ABC transporter | NA | -1.711 | 2.07E-03 |
| PA3406 | *hasD* | transport protein HasD | -2.208 | 5.22E-04 |
| PA3409 | *hasS* | NA | -1.878 | 7.94E-03 |
| PA3412 | hypothetical protein | NA | -1.169 | 1.79E-02 |
| PA3414 | hypothetical protein | NA | -1.780 | 1.51e-08 |
| PA3419 | hypothetical protein | NA | -1.599 | 4.33E-04 |
| PA3420 | probable transcriptional regulator | NA | -2.346 | 1.51e-08 |
| PA3424 | hypothetical protein | NA | -1.168 | 1.87E-03 |
| PA3426 | probable enoyl CoA-hydratase/isomerase | probable enoyl CoA-hydratase/isomerase | -1.539 | 1.63e-05 |
| PA3442 | probable ATP-binding component of ABC transporter | probable ATP-binding component of ABC transporter | -1.177 | 3.07E-03 |

| Gene | Name | Product | $log_2$ FC | Adjusted p-value |
|---|---|---|---|---|
| PA3444 | conserved hypothetical protein | conserved hypothetical protein | -1.840 | 5.39e-06 |
| PA3445 | conserved hypothetical protein | conserved hypothetical protein | -1.526 | 1.35E-04 |
| PA3446 | conserved hypothetical protein | conserved hypothetical protein | -2.409 | 1.23e-06 |
| PA3447 | probable ATP-binding component of ABC transporter | probable ATP-binding component of ABC transporter | -1.414 | 4.01E-03 |
| PA3448 | probable permease of ABC transporter | probable permease of ABC transporter | -1.462 | 2.78E-02 |
| PA3449 | conserved hypothetical protein | conserved hypothetical protein | -1.767 | 9.06E-04 |
| PA3450 | *lsfA* | NA | -1.136 | 4.76E-02 |
| PA3458 | probable transcriptional regulator | NA | 1.014 | 3.47E-02 |
| PA3459 | probable glutamine amidotransferas e | probable glutamine amidotransferas e | 1.227 | 3.61E-03 |
| PA3468 | conserved hypothetical protein | NA | -1.043 | 1.51E-02 |
| PA3470 | NUDIX hydrolase | NA | -1.667 | 1.25e-06 |
| PA3483 | hypothetical protein | NA | -1.055 | 1.69E-03 |
| PA3493 | conserved hypothetical protein | NA | -1.349 | 2.91E-03 |
| PA3510 | hypothetical protein | NA | 1.406 | 4.46E-04 |
| PA3521 | *opmE* | NA | -2.020 | 2.12e-06 |
| PA3529 | alkylhydroperox ide reductase C | NA | 1.199 | 9.40E-03 |
| PA3535 | probable serine protease | NA | -1.036 | 2.66E-03 |
| PA3543 | *algK* | NA | -1.247 | 1.02E-02 |
| PA3552 | *arnB* | ArnB | -4.114 | 1.28e-16 |
| PA3553 | *arnC* | ArnC | -3.709 | 1.45e-14 |
| PA3554 | *arnA* | ArnA | -2.231 | 2.22e-06 |
| PA3555 | *arnD* | ArnD | -3.389 | 1.07e-06 |
| PA3556 | *arnT* | inner membrane L-Ara4N | -2.182 | 2.19e-09 |

| Gene | Name | Product | $log_2$ FC | Adjusted p-value |
|---|---|---|---|---|
| | | transferase ArnT | | |
| PA3557 | *arnE* | ArnE | -3.180 | 5.78E-03 |
| PA3558 | *arnF* | ArnF | -2.564 | 2.05E-03 |
| PA3559 | probable nucleotide sugar dehydrogenase | probable nucleotide sugar dehydrogenase | -1.551 | 3.04E-04 |
| PA3564 | conserved hypothetical protein | NA | -1.221 | 1.71E-02 |
| PA3575 | hypothetical protein | NA | -1.346 | 9.87E-04 |
| PA3582 | *glpK* | glycerol kinase | 1.086 | 7.54E-03 |
| PA3589 | probable acyl-CoA thiolase | probable acyl-CoA thiolase | -1.319 | 3.32E-02 |
| PA3591 | probable enoyl-CoA hydratase/isomerase | NA | -1.664 | 9.83E-03 |
| PA3592 | conserved hypothetical protein | NA | -1.289 | 2.67E-02 |
| PA3613 | hypothetical protein | NA | 1.061 | 3.08E-02 |
| PA3629 | *adhC* | alcohol dehydrogenase class III | 1.027 | 1.32E-03 |
| PA3632 | conserved hypothetical protein | NA | 1.109 | 1.31E-02 |
| PA3669 | hypothetical protein | NA | -1.409 | 2.84E-04 |
| PA3670 | hypothetical protein | NA | -1.431 | 4.19E-04 |
| PA3672 | probable ATP-binding component of ABC transporter | NA | -1.733 | 1.74E-04 |
| PA3676 | *mexK* | NA | 1.125 | 6.02e-05 |
| PA3680 | conserved hypothetical protein | NA | -1.106 | 6.74E-04 |
| PA3711 | probable transcriptional regulator | NA | -1.197 | 9.97E-04 |
| PA3712 | hypothetical protein | NA | -2.119 | 9.24e-07 |
| PA3765 | hypothetical protein | NA | -1.382 | 3.38E-04 |
| PA3768 | probable metallo-oxidoreductase | NA | 1.039 | 9.99E-04 |

| Gene | Name | Product | $log_2$ FC | Adjusted p-value |
|---|---|---|---|---|
| PA3773 | hypothetical protein | NA | -2.681 | 1.58E-04 |
| PA3775 | hypothetical protein | NA | -1.571 | 1.28E-02 |
| PA3787 | putative peptidase | NA | -1.219 | 8.61E-03 |
| PA3839 | probable sodium:sulfate symporter | NA | 1.578 | 7.82e-06 |
| PA3864 | *dauR* | NA | 1.097 | 6.96E-04 |
| PA3870 | *moaA1* | molybdopterin biosynthetic protein A1 | -1.686 | 1.02E-03 |
| PA3880 | conserved hypothetical protein | NA | -1.238 | 3.42E-03 |
| PA3883 | probable short-chain dehydrogenase | NA | -1.238 | 3.24E-03 |
| PA3904 | *PAAR4* | NA | -1.650 | 1.45E-03 |
| PA3905 | *tecT* | NA | -1.094 | 1.19E-02 |
| PA3919 | conserved hypothetical protein | NA | 1.354 | 2.72E-03 |
| PA3928 | hypothetical protein | NA | -1.799 | 3.01e-08 |
| PA3931 | putative methionine-binding protein | putative methionine-binding protein | -1.431 | 7.00E-03 |
| PA3936 | probable permease of ABC taurine transporter | probable permease of ABC taurine transporter | -1.374 | 8.70E-03 |
| PA3937 | probable ATP-binding component of ABC taurine transporter | probable ATP-binding component of ABC taurine transporter | -1.563 | 4.53E-04 |
| PA3938 | probable periplasmic taurine-binding protein precursor | probable periplasmic taurine-binding protein precursor | -1.227 | 6.30E-03 |
| PA3964 | hypothetical protein | NA | -2.205 | 8.95e-09 |
| PA3985 | conserved hypothetical protein | NA | -1.017 | 2.65E-03 |
| PA4014 | hypothetical protein | NA | -1.141 | 6.38E-03 |
| PA4059 | hypothetical protein | NA | -1.370 | 3.25E-02 |
| PA4067 | *oprG* | NA | 1.229 | 2.20E-03 |

| Gene | Name | Product | $\log_2$ FC | Adjusted p-value |
|---|---|---|---|---|
| PA4074 | probable transcriptional regulator | NA | -1.130 | 5.40E-03 |
| PA4075 | hypothetical protein | NA | -1.337 | 1.41E-04 |
| PA4076 | hypothetical protein | NA | -1.502 | 2.97E-04 |
| PA4093 | hypothetical protein | NA | -1.141 | 2.92E-02 |
| PA4096 | probable major facilitator superfamily (MFS) transporter | NA | -1.877 | 2.48E-03 |
| PA4097 | probable alcohol dehydrogenase (Zn-dependent) | probable alcohol dehydrogenase (Zn-dependent) | -1.708 | 3.28E-02 |
| PA4098 | probable short-chain dehydrogenase | NA | -2.167 | 1.59E-04 |
| PA4129 | hypothetical protein | NA | -1.101 | 4.50E-02 |
| PA4130 | *nirA* | ferredoxin-dependent nitrite reductase | -1.496 | 3.84e-06 |
| PA4131 | probable iron-sulfur protein | NA | -1.889 | 1.9e-06 |
| PA4139 | hypothetical protein | NA | -1.863 | 3.07E-03 |
| PA4140 | hypothetical protein | NA | -1.681 | 2.29E-04 |
| PA4142 | probable secretion protein | NA | -1.171 | 3.12E-03 |
| PA4144 | probable outer membrane protein precursor | probable outer membrane protein precursor | -1.142 | 4.56E-03 |
| PA4148 | probable short-chain dehydrogenase | probable short-chain dehydrogenase | -1.564 | 4.37E-03 |
| PA4159 | *fepB* | ferrienterobactin-binding periplasmic protein precursor FepB | -1.701 | 3.48E-03 |
| PA4160 | *fepD* | ferric enterobactin transport protein FepD | -1.955 | 2.83E-03 |
| PA4161 | *fepG* | ferric enterobactin | -1.568 | 5.25E-04 |

| Gene | Name | Product | $log_2$ FC | Adjusted p-value |
|---|---|---|---|---|
| | | transport protein FepG | | |
| PA4177 | hypothetical protein | NA | -1.368 | 5.49E-03 |
| PA4181 | hypothetical protein | NA | 1.103 | 4.55E-02 |
| PA4182 | hypothetical protein | NA | 1.807 | 1.05E-04 |
| PA4183 | hypothetical protein | NA | 1.584 | 1.33e-06 |
| PA4196 | *bfiR* | NA | 1.227 | 7.67E-04 |
| PA4198 | probable AMP-binding enzyme | NA | 1.150 | 6.43E-04 |
| PA4205 | *mexG* | NA | -1.379 | 4.31e-05 |
| PA4206 | *mexH* | probable Resistance-Nodulation-Cell Division (RND) efflux membrane fusion protein precursor | -1.525 | 5.27e-07 |
| PA4208 | *opmD* | NA | -2.193 | 3.48e-07 |
| PA4210 | *phzA1* | NA | -2.458 | 3.89E-04 |
| PA4211 | *phzB1* | probable phenazine biosynthesis protein | -1.849 | 3.61E-04 |
| PA4218 | *ampP* | NA | -1.160 | 4.99E-02 |
| PA4226 | *pchE* | dihydroaeruginoic acid synthetase | -1.464 | 2.52E-04 |
| PA4236 | *katA* | catalase | 1.366 | 1.44E-04 |
| PA4288 | probable transcriptional regulator | NA | 1.134 | 2.08E-02 |
| PA4299 | *tadD* | NA | -2.141 | 1.75E-03 |
| PA4305 | *rcpC* | NA | -1.058 | 9.79E-03 |
| PA4325 | hypothetical protein | NA | -1.028 | 7.79E-04 |
| PA4331 | probable ferredoxin reductase | NA | -1.466 | 5.62E-04 |
| PA4343 | probable major facilitator superfamily (MFS) transporter | NA | -1.351 | 7.94E-03 |
| PA4353 | conserved hypothetical protein | NA | 1.608 | 2.75E-04 |

| Gene | Name | Product | $log_2$ FC | Adjusted p-value |
|---|---|---|---|---|
| PA4354 | conserved hypothetical protein | NA | 2.990 | 4.9e-19 |
| PA4355 | *pyeM* | NA | 1.738 | 1.89e-06 |
| PA4356 | *xenB* | NA | 3.222 | 1.01e-31 |
| PA4358 | *feoB* | NA | 1.091 | 8.34e-05 |
| PA4364 | hypothetical protein | NA | 1.871 | 2.61E-04 |
| PA4365 | *lysE* | NA | 1.967 | 3.81E-04 |
| PA4371 | hypothetical protein | NA | -1.139 | 1.60E-03 |
| PA4385 | *groEL* | GroEL protein | 1.047 | 3.38E-04 |
| PA4386 | *groES* | NA | 1.045 | 8.25e-05 |
| PA4420 | conserved hypothetical protein | NA | 1.029 | 4.62E-02 |
| PA4463 | conserved hypothetical protein | NA | 1.352 | 9.76E-04 |
| PA4464 | *ptsN* | nitrogen regulatory IIA protein | 1.115 | 1.10E-03 |
| PA4472 | *pmbA* | NA | 1.089 | 7.28e-05 |
| PA4474 | conserved hypothetical protein | NA | 1.256 | 1.14E-04 |
| PA4507 | hypothetical protein | NA | -1.328 | 7.07E-03 |
| PA4523 | hypothetical protein | NA | 1.286 | 1.79E-02 |
| PA4525 | *pilA* | type 4 fimbrial precursor PilA | 1.530 | 9.87E-04 |
| PA4542 | *clpB* | NA | 1.429 | 6.62E-04 |
| PA4550 | *fimU* | NA | 1.849 | 1.7e-08 |
| PA4551 | *pilV* | NA | 1.511 | 6.44e-07 |
| PA4552 | *pilW* | NA | 1.792 | 1.7e-08 |
| PA4553 | *pilX* | NA | 1.119 | 5.15E-03 |
| PA4554 | *pilY1* | NA | 1.323 | 5.2e-06 |
| PA4555 | *pilY2* | NA | 1.638 | 6.44e-07 |
| PA4556 | *pilE* | NA | 1.544 | 2.93e-07 |
| PA4571 | probable cytochrome c | NA | 1.322 | 1.44E-04 |
| PA4593 | probable permease of ABC transporter | NA | -1.040 | 2.33E-02 |
| PA4599 | *mexC* | NA | -1.457 | 2.31E-04 |
| PA4604 | conserved hypothetical protein | NA | 1.168 | 2.48e-05 |
| PA4611 | hypothetical protein | NA | 1.493 | 2.84E-03 |
| PA4615 | *fprB* | NA | 1.020 | 3.36E-03 |

| Gene | Name | Product | $log_2$ FC | Adjusted p-value |
|---|---|---|---|---|
| PA4620 | hypothetical protein | NA | 1.106 | 5.80E-04 |
| PA4622 | probable major facilitator superfamily (MFS) transporter | NA | 1.991 | 2.85e-12 |
| PA4623 | hypothetical protein | NA | 4.746 | 1.45e-64 |
| PA4635 | conserved hypothetical protein | NA | -1.144 | 1.10E-03 |
| PA4651 | *cupE4* | NA | -1.694 | 1.59E-03 |
| PA4658 | hypothetical protein | NA | 1.185 | 2.33E-03 |
| PA4659 | probable transcriptional regulator | NA | 1.435 | 7.23E-04 |
| PA4660 | *phr* | NA | 1.145 | 9.87E-04 |
| PA4661 | *pagL* | NA | 1.778 | 1.2e-08 |
| PA4679 | *holD* | NA | -1.157 | 3.78E-04 |
| PA4682 | hypothetical protein | NA | 1.076 | 3.30E-03 |
| PA4683 | hypothetical protein | NA | 1.326 | 4.42E-04 |
| PA4689 | hypothetical protein | NA | 1.416 | 4.4e-08 |
| PA4714 | conserved hypothetical protein | NA | -1.811 | 2.27E-04 |
| PA4752 | *ftsJ* | NA | 1.091 | 1.16E-04 |
| PA4759 | *dapB* | dihydrodipicolinate reductase | 1.050 | 3.20E-04 |
| PA4761 | *dnaK* | DnaK protein | 1.307 | 8.54e-05 |
| PA4762 | *grpE* | NA | 1.071 | 4.07E-03 |
| PA4763 | *recN* | NA | -1.169 | 1.00E-04 |
| PA4773 | *speD2* | SpeD2 | -1.164 | 3.12E-03 |
| PA4774 | *speE2* | SpeE2 | -1.340 | 7.55e-05 |
| PA4779 | hypothetical protein | NA | -1.290 | 9.29e-05 |
| PA4816 | hypothetical protein | NA | -1.947 | 4.62E-04 |
| PA4818 | conserved hypothetical protein | NA | -2.312 | 6.1e-05 |
| PA4819 | probable glycosyl transferase | NA | -1.401 | 1.53E-02 |
| PA4820 | hypothetical protein | NA | -1.239 | 3.42E-03 |
| PA4824 | hypothetical protein | NA | -1.877 | 4.31E-04 |

| Gene | Name | Product | $log_2$ FC | Adjusted p-value |
|---|---|---|---|---|
| PA4871 | hypothetical protein | NA | -2.287 | 1.23e-06 |
| PA4881 | hypothetical protein | NA | 3.790 | 8.89e-34 |
| PA4882 | hypothetical protein | NA | 1.078 | 7.89E-03 |
| PA4884 | hypothetical protein | NA | -1.843 | 9.03E-04 |
| PA4890 | *desT* | NA | -1.190 | 1.37E-03 |
| PA4893 | *ureG* | NA | 1.050 | 2.22E-02 |
| PA4918 | *pcnA* | nicotinamidase | 1.423 | 1.78e-06 |
| PA4919 | *pncB1* | nicotinate phosphoribosyltransferase | 1.972 | 4.35e-14 |
| PA4920 | *nadE* | NH3-dependent NAD synthetase | 1.327 | 3.02e-05 |
| PA4944 | *hfq* | Hfq | 1.185 | 2.09E-04 |
| PA4985 | Uncharacterized protein | Uncharacterized protein | -1.411 | 3.30E-04 |
| PA4986 | probable oxidoreductase | NA | -1.334 | 1.27E-03 |
| PA4988 | *waaA* | 3-deoxy-D-manno-octulosonic-acid (KDO) transferase | -1.290 | 4.33E-04 |
| PA5024 | conserved hypothetical protein | NA | -1.227 | 1.00E-02 |
| PA5031 | probable short chain dehydrogenase | NA | -1.566 | 3.92E-03 |
| PA5032 | probable transcriptional regulator | NA | -1.286 | 1.00E-02 |
| PA5040 | *pilQ* | NA | 1.554 | 1.02E-03 |
| PA5041 | *pilP* | NA | 1.441 | 9.65E-03 |
| PA5042 | *pilO* | NA | 1.574 | 3.42E-03 |
| PA5043 | *pilN* | NA | 1.400 | 1.20E-02 |
| PA5044 | *pilM* | NA | 1.012 | 5.00E-02 |
| PA5072 | *mcpK* | McpK | -1.046 | 9.17E-03 |
| PA5111 | *gloA3* | lactoylglutathione lyase | 1.124 | 4.43E-04 |
| PA5112 | *estA* | NA | 1.096 | 7.40E-03 |
| PA5119 | *glnA* | glutamine synthetase | 1.124 | 3.88E-02 |
| PA5135 | conserved hypothetical protein | NA | -1.077 | 2.01E-03 |
| PA5145 | hypothetical protein | NA | -1.031 | 5.96E-03 |

| Gene | Name | Product | $log_2$ FC | Adjusted p-value |
|---|---|---|---|---|
| PA5153 | amino acid (lysine/arginine/ ornithine/histidine/octopine) ABC transporter periplasmic binding protein | NA | 1.136 | 8.19E-04 |
| PA5154 | probable permease of ABC transporter | NA | 1.184 | 2.86E-03 |
| PA5155 | amino acid (lysine/arginine/ ornithine/histidine/octopine) ABC transporter membrane protein | NA | 1.276 | 1.27E-03 |
| PA5157 | probable transcriptional regulator | NA | 1.609 | 2.03E-04 |
| PA5159 | multidrug resistance protein | NA | 1.187 | 7.94E-03 |
| PA5170 | *arcD* | NA | 2.064 | 1.56E-04 |
| PA5171 | *arcA* | arginine deiminase | 1.406 | 7.86E-03 |
| PA5172 | *arcB* | ornithine carbamoyltransferase | 1.593 | 4.66E-04 |
| PA5173 | *arcC* | carbamate kinase | 1.048 | 3.85E-02 |
| PA5183.1 | *rsmN* | NA | 1.333 | 6.57E-04 |
| PA5200 | *amgR* | AmgR | -1.054 | 2.18E-02 |
| PA5202 | thiolester hydrolase | thiolester hydrolase | -1.273 | 2.26E-03 |
| PA5208 | conserved hypothetical protein | NA | 1.074 | 3.76E-03 |
| PA5219 | hypothetical protein | NA | -1.194 | 1.82E-03 |
| PA5229 | conserved hypothetical protein | NA | 1.135 | 5.84e-05 |
| PA5230 | probable permease of ABC transporter | NA | 1.751 | 7.2e-13 |
| PA5231 | probable ATP-binding/permease fusion ABC transporter | NA | 1.456 | 2.16E-04 |
| PA5328 | *sphB* | NA | -1.277 | 3.30E-02 |
| PA5341 | hypothetical protein | NA | -1.927 | 1.33e-05 |

| Gene | Name | Product | $log_2$ FC | Adjusted p-value |
|---|---|---|---|---|
| PA5356 | *glcC* | NA | -1.322 | 4.57E-03 |
| PA5357 | hypothetical protein | hypothetical protein | -1.105 | 1.27E-03 |
| PA5372 | *betA* | choline dehydrogenase | 1.059 | 7.19E-04 |
| PA5376 | *cbcV* | CbcV | 1.346 | 2.45E-04 |
| PA5378 | *cbcX* | CbcX | 1.041 | 5.49E-03 |
| PA5386 | *cdhA* | NA | -1.858 | 5.39E-04 |
| PA5388 | *caiX* | CaiX | -1.096 | 3.72E-02 |
| PA5391 | hypothetical protein | NA | -2.523 | 5.77E-04 |
| PA5431 | probable transcriptional regulator | NA | -1.089 | 1.25E-02 |
| PA5453 | *gmd* | GDP-mannose 4 | 1.483 | 2.32e-09 |
| PA5475 | hypothetical protein | NA | 1.519 | 8.62E-04 |
| PA5507 | hypothetical protein | NA | 1.214 | 1.39E-02 |
| PA5527 | hypothetical protein | NA | -1.017 | 3.38E-02 |
| PA5528 | hypothetical protein | NA | -1.745 | 4.46E-04 |
| PA5530 | C5-dicarboxylate transporter | NA | -1.267 | 1.10E-02 |

### *Supplementary Dataset E.*

**List of transcripts with significant differences in abundance in PAO1$_{MW}$ poly-cultures (+) *versus* *mexT*$^{I129F}$ mutant poly-cultures (-)**

| Gene | Name | Product | $log_2$ FC | Adjusted p-value |
|---|---|---|---|---|
| PA0049 | hypothetical protein | NA | -1.158 | 4.86E-02 |
| PA0058 | *dsbM* | NA | -2.003 | 3.82E-04 |
| PA0125 | ParD antitoxin | NA | -1.398 | 4.89E-02 |
| PA0152 | *pcaQ* | NA | -1.229 | 7.83E-04 |
| PA0157 | *triB* | NA | -1.011 | 7.82E-03 |
| PA0165 | hypothetical protein | NA | -1.171 | 8.11e-05 |
| PA0182 | probable short-chain dehydrogenase | probable short-chain dehydrogenase | -2.401 | 9.99E-04 |
| PA0213 | hypothetical protein | NA | -1.929 | 2.14E-02 |
| PA0263 | *hcpC* | secreted protein Hcp | -4.077 | 4.21e-09 |
| PA0282 | *cysT* | sulfate transport protein CysT | -1.851 | 2.43E-04 |
| PA0283 | *sbp* | sulfate-binding protein precursor | -2.536 | 3.62e-05 |
| PA0284 | hypothetical protein | NA | -2.593 | 1.26E-04 |

| | | | | |
|---|---|---|---|---|
| PA0380 | conserved hypothetical protein | conserved hypothetical protein | -1.742 | 1.47E-03 |
| PA0422 | conserved hypothetical protein | NA | -1.443 | 1.19E-03 |
| PA0451 | conserved hypothetical protein | NA | -1.883 | 1.27E-03 |
| PA0456 | probable cold-shock protein | NA | 1.039 | 1.98E-02 |
| PA0468 | hypothetical protein | NA | -1.144 | 2.14E-02 |
| PA0489 | probable phosphoribosyl transferase | NA | -1.186 | 4.36E-02 |
| PA0514 | *nirL* | NA | -1.477 | 4.63E-02 |
| PA0522 | hypothetical protein | NA | -1.841 | 4.27E-03 |
| PA0529 | conserved hypothetical protein | NA | 1.521 | 1.44E-02 |
| PA0531 | probable glutamine amidotransferase | NA | 1.275 | 6.36e-05 |
| PA0615 | hypothetical protein | NA | 1.295 | 3.40E-03 |
| PA0617 | probable bacteriophage protein | NA | 1.444 | 1.73E-02 |
| PA0618 | probable bacteriophage protein | NA | 1.603 | 9.35E-04 |
| PA0619 | probable bacteriophage protein | NA | 1.543 | 5.30E-03 |
| PA0633 | hypothetical protein | NA | 1.889 | 7.67E-04 |
| PA0634 | hypothetical protein | NA | 1.902 | 3.51E-03 |
| PA0636 | hypothetical protein | NA | 1.472 | 8.54E-03 |
| PA0637 | conserved hypothetical protein | NA | 1.167 | 3.81E-02 |
| PA0670 | hypothetical protein | NA | -1.492 | 5.84E-04 |
| PA0671 | hypothetical protein | NA | -2.060 | 1.55e-06 |
| PA0695 | hypothetical protein | NA | -1.792 | 4.66E-02 |
| PA0715 | hypothetical protein | NA | -4.546 | 1.74E-02 |
| PA0716.2 | *xisF4* | NA | -10.648 | 1.82e-13 |
| PA0717 | hypothetical protein of bacteriophage Pf1 | NA | -10.675 | 1.76e-15 |
| PA0718 | hypothetical protein of bacteriophage Pf1 | NA | -10.632 | 4.3e-16 |
| PA0719 | hypothetical protein of bacteriophage Pf1 | NA | -11.492 | 3.62e-20 |
| PA0720 | helix destabilizing protein of bacteriophage Pf1 | NA | -13.267 | 1.93e-35 |
| PA0721 | *pfsE* | NA | -11.014 | 1.44e-20 |
| PA0724 | probable coat protein A of bacteriophage Pf1 | NA | -9.796 | 4.05e-14 |
| PA0726 | hypothetical protein of bacteriophage Pf1 | NA | -8.619 | 1.03e-10 |

| | | | | |
|---|---|---|---|---|
| PA0727 | Pf replication initiator protein | NA | -9.001 | 4.62e-11 |
| PA0728 | probable bacteriophage integrase | NA | -7.800 | 5.62e-08 |
| PA0728.1 | *pfiA* | NA | -4.102 | 1.16E-02 |
| PA0729 | *pfiT* | NA | -3.841 | 1.54E-02 |
| PA0758 | hypothetical protein | NA | 1.190 | 6.75E-03 |
| PA0780 | *pruR* | NA | -1.310 | 5.27E-03 |
| PA0801 | hypothetical protein | NA | -1.224 | 4.88E-03 |
| PA0806 | hypothetical protein | NA | -1.395 | 4.86E-02 |
| PA0813 | hypothetical protein | NA | -1.130 | 4.92E-02 |
| PA0816 | probable transcriptional regulator | NA | -1.381 | 2.43E-02 |
| PA0855 | hypothetical protein | NA | -1.078 | 8.83E-03 |
| PA0952 | hypothetical protein | NA | 1.290 | 2.48E-02 |
| PA0958 | *oprD* | Basic amino acid | -1.242 | 5.03e-08 |
| PA0979 | conserved hypothetical protein | NA | -1.626 | 3.12E-02 |
| PA0996 | *pqsA* | PqsA | -1.203 | 4.19E-02 |
| PA1063 | hypothetical protein | NA | 1.063 | 1.22E-02 |
| PA1089 | conserved hypothetical protein | NA | 1.095 | 1.03E-03 |
| PA1103 | probable flagellar assembly protein | probable flagellar assembly protein | 1.129 | 1.19E-03 |
| PA1109 | probable transcriptional regulator | NA | -1.057 | 4.19E-02 |
| PA1125 | probable cobalamin biosynthetic protein | NA | -1.239 | 5.71E-03 |
| PA1146 | probable iron-containing alcohol dehydrogenase | probable iron-containing alcohol dehydrogenase | -1.263 | 1.39E-02 |
| PA1176 | *napF* | NA | -1.938 | 2.91E-02 |
| PA1178 | *oprH* | NA | -4.388 | 3.98E-02 |
| PA1180 | *phoQ* | two-component sensor PhoQ | -2.173 | 2.96E-02 |
| PA1221 | nonribosomal peptide synthetase | NA | -2.157 | 4.59E-03 |
| PA1223 | probable transcriptional regulator | NA | -1.048 | 4.00E-02 |
| PA1246 | *aprD* | alkaline protease secretion protein AprD | -1.672 | 2.43E-04 |
| PA1248 | *aprF* | NA | -1.076 | 1.39E-02 |
| PA1275 | *cobD* | cobalamin biosynthetic protein CobD | -1.547 | 1.97E-02 |
| PA1281 | *cobV* | cobalamin (5'-phosphate) synthase | -2.924 | 1.49e-06 |

| | | | | |
|---|---|---|---|---|
| PA1282 | probable major facilitator superfamily (MFS) transporter | NA | -1.335 | 1.28E-03 |
| PA1285 | probable transcriptional regulator | NA | -1.865 | 9.25e-08 |
| PA1286 | probable major facilitator superfamily (MFS) transporter | NA | -2.568 | 1.63e-11 |
| PA1298 | conserved hypothetical protein | NA | -1.510 | 3.90E-03 |
| PA1317 | *cyoA* | cytochrome o ubiquinol oxidase subunit II | 1.226 | 3.24E-04 |
| PA1318 | *cyoB* | cytochrome o ubiquinol oxidase subunit I | 1.036 | 1.22E-02 |
| PA1319 | *cyoC* | cytochrome o ubiquinol oxidase subunit III | 1.222 | 1.91E-02 |
| PA1320 | *cyoD* | cytochrome o ubiquinol oxidase subunit IV | 1.484 | 3.78E-03 |
| PA1328 | probable transcriptional regulator | NA | -1.437 | 3.74E-02 |
| PA1329 | conserved hypothetical protein | NA | -2.115 | 3.40E-03 |
| PA1332 | hypothetical protein | NA | 2.303 | 1.38e-10 |
| PA1333 | hypothetical protein | NA | 1.400 | 2.51E-02 |
| PA1334 | probable oxidoreductase | probable oxidoreductase | -1.147 | 4.59E-03 |
| PA1379 | probable short-chain dehydrogenase | NA | -1.041 | 2.14E-02 |
| PA1512 | *hcpA* | secreted protein Hcp | -2.372 | 2.29E-03 |
| PA1544 | *anr* | NA | 1.021 | 7.83E-04 |
| PA1566 | *pauA3* | Glutamylpolyamine synthetase | -1.096 | 1.12E-02 |
| PA1585 | *sucA* | 2-oxoglutarate dehydrogenase (E1 subunit) | 1.001 | 1.74E-02 |
| PA1592 | hypothetical protein | NA | -1.175 | 3.83E-02 |
| PA1617 | probable AMP-binding enzyme | NA | -1.009 | 1.90E-02 |
| PA1670 | stp1 | Stp1 | -1.263 | 1.22E-02 |
| PA1744 | hypothetical protein | NA | 2.887 | 1.77e-14 |
| PA1849 | conserved hypothetical protein | NA | -2.164 | 4.37E-02 |
| PA1881 | probable oxidoreductase | NA | -1.067 | 2.43E-02 |
| PA1884 | transcriptional regulator | NA | -1.544 | 3.53E-03 |

| | | | | |
|---|---|---|---|---|
| PA1885 | conserved hypothetical protein | NA | -1.671 | 2.30E-02 |
| PA1891 | hypothetical protein | NA | -2.353 | 2.69E-02 |
| PA1911 | *femR* | NA | -2.160 | 3.27E-02 |
| PA1919 | *nrdG* | NA | -1.356 | 1.41E-02 |
| PA1932 | probable hydroxylase molybdopterin-containing subunit | probable hydroxylase molybdopterin-containing subunit | -1.042 | 4.19E-02 |
| PA1937 | conserved hypothetical protein | NA | -1.256 | 7.96E-03 |
| PA1941 | hypothetical protein | NA | 2.278 | 2.21e-17 |
| PA1942 | hypothetical protein | NA | 6.263 | 1.66e-90 |
| PA1970 | hypothetical protein | NA | 4.029 | 1.25e-13 |
| PA2019 | *mexX* | Resistance-Nodulation-Cell Division (RND) multidrug efflux membrane fusion protein MexX precursor | -1.615 | 3.03E-02 |
| PA2040 | *pauA4* | Glutamylpolyamine synthetase | -1.406 | 6.31e-06 |
| PA2048 | hypothetical protein | NA | -1.577 | 2.07E-02 |
| PA2062 | probable pyridoxal-phosphate dependent enzyme | probable pyridoxal-phosphate dependent enzyme | -1.494 | 3.18E-02 |
| PA2069 | probable carbamoyl transferase | NA | -1.253 | 1.57E-02 |
| PA2081 | *kynB* | kynurenine formamidase | -1.247 | 1.19E-02 |
| PA2143 | hypothetical protein | NA | -1.474 | 3.17E-02 |
| PA2190 | conserved hypothetical protein | NA | -1.289 | 2.07E-02 |
| PA2204 | putative binding protein component of ABC transporter | NA | -2.307 | 1.36E-03 |
| PA2241 | *pslK* | PslL | -1.233 | 8.65E-03 |
| PA2243 | *pslM* | NA | -1.070 | 1.38E-02 |
| PA2245 | *pslO* | NA | -2.451 | 3.47E-03 |
| PA2253 | *ansA* | L-asparaginase I | -1.240 | 1.43E-02 |
| PA2268 | hypothetical protein | NA | -1.774 | 8.98e-05 |
| PA2275 | probable alcohol dehydrogenase (Zn-dependent) | NA | -1.416 | 4.22E-03 |
| PA2277 | *arsR* | NA | -1.457 | 8.87E-03 |
| PA2279 | *arsC* | NA | -1.101 | 1.19E-03 |
| PA2283 | hypothetical protein | NA | -1.468 | 2.63E-03 |
| PA2284 | hypothetical protein | NA | -1.663 | 4.12E-03 |
| PA2287 | hypothetical protein | NA | -1.637 | 4.12E-03 |
| PA2288 | hypothetical protein | NA | -1.901 | 4.78E-04 |
| PA2305 | *ambB* | NA | -1.391 | 1.72E-02 |
| PA2310 | hypothetical protein | hypothetical protein | -1.217 | 2.14E-02 |

| | | | | |
|---|---|---|---|---|
| PA2311 | hypothetical protein | NA | -2.356 | 1.90E-03 |
| PA2312 | probable transcriptional regulator | NA | -1.760 | 2.51E-02 |
| PA2316 | probable transcriptional regulator | NA | -1.116 | 2.18E-02 |
| PA2344 | *mtlZ* | fructokinase | -1.258 | 2.88E-02 |
| PA2356 | *msuD* | methanesulfonate sulfonatase MsuD | -1.393 | 3.22E-02 |
| PA2358 | hypothetical protein | NA | -1.989 | 2.84E-02 |
| PA2359 | probable transcriptional regulator | NA | -1.623 | 3.12E-02 |
| PA2375 | hypothetical protein | NA | -1.170 | 2.45E-02 |
| PA2378 | probable aldehyde dehydrogenase | NA | -1.193 | 1.39E-02 |
| PA2477 | *dsbE* | NA | -1.136 | 2.43E-02 |
| PA2480 | *dsbS* | NA | -1.107 | 2.23E-03 |
| PA2485 | hypothetical protein | NA | 2.923 | 4.09e-16 |
| PA2486 | *ptrC* | NA | 3.939 | 1.07e-39 |
| PA2487 | hypothetical protein | NA | 2.475 | 4.88e-15 |
| PA2490 | conserved hypothetical protein | NA | 2.869 | 2.66e-21 |
| PA2491 | *mexS* | NA | 4.001 | 9.19e-61 |
| PA2493 | *mexE* | NA | 5.979 | 1.36e-88 |
| PA2494 | *mexF* | NA | 6.910 | 5.1e-95 |
| PA2495 | *oprN* | NA | 6.234 | 2.92e-84 |
| PA2496 | hypothetical protein | NA | 1.540 | 1.41E-04 |
| PA2522 | *czcC* | NA | -2.088 | 4.45E-02 |
| PA2523 | *czcR* | CzcR | -2.430 | 1.62E-02 |
| PA2605 | conserved hypothetical protein | conserved hypothetical protein | -2.044 | 4.09e-08 |
| PA2758 | probable transcriptional regulator | NA | 2.174 | 2.43e-06 |
| PA2786 | hypothetical protein | NA | -1.343 | 6.66E-03 |
| PA2810 | *copS* | two-component sensor | 1.087 | 6.10E-03 |
| PA2811 | probable permease of ABC-2 transporter | NA | 3.013 | 1.48e-20 |
| PA2812 | probable ATP-binding component of ABC transporter | NA | 2.719 | 8.72e-15 |
| PA2813 | probable glutathione S-transferase | probable glutathione S-transferase | 3.183 | 7.24e-21 |
| PA2857 | probable ATP-binding component of ABC transporter | NA | -1.213 | 3.22E-02 |
| PA2869 | hypothetical protein | NA | -1.188 | 1.41E-03 |
| PA2914 | probable permease of ABC transporter | probable permease of ABC transporter | -2.050 | 2.79e-06 |
| PA2915 | hypothetical protein | NA | -1.376 | 3.43E-03 |

| | | | | |
|---|---|---|---|---|
| PA2919 | hypothetical protein | NA | -1.636 | 4.86E-02 |
| PA2937 | hypothetical protein | NA | -1.708 | 4.77E-02 |
| PA2995 | *nqrE* | NA | 1.003 | 3.67E-02 |
| PA2996 | *nqrD* | NA | 1.210 | 3.43E-03 |
| PA3060 | *pelE* | PelE | -1.709 | 1.06E-03 |
| PA3095 | *xcpZ* | general secretion pathway protein M | -1.461 | 4.31E-02 |
| PA3104 | *xcpP* | secretion protein XcpP | -1.103 | 9.59e-06 |
| PA3144 | transposase with Helix-turn-helix Hin domain | NA | -1.577 | 3.66e-06 |
| PA3193 | *glk* | glucokinase | -1.068 | 4.97E-02 |
| PA3205 | hypothetical protein | NA | -2.027 | 2.96E-02 |
| PA3206 | *cpxS* | NA | -1.753 | 1.56E-02 |
| PA3207 | hypothetical protein | NA | -1.364 | 1.39E-02 |
| PA3208 | conserved hypothetical protein | NA | -1.213 | 1.03E-02 |
| PA3214 | hypothetical protein | NA | -1.053 | 4.34E-02 |
| PA3229 | hypothetical protein | NA | 6.009 | 4.58e-86 |
| PA3230 | conserved hypothetical protein | NA | 4.264 | 2e-63 |
| PA3231 | hypothetical protein | NA | 3.658 | 9.5e-21 |
| PA3232 | probable nuclease | probable nuclease | 2.568 | 7.57e-16 |
| PA3233 | hypothetical protein | NA | 2.144 | 9.89e-12 |
| PA3266 | *capB* | NA | 1.059 | 1.74E-02 |
| PA3306 | hypothetical protein | NA | -1.355 | 1.98E-02 |
| PA3321 | probable transcriptional regulator | NA | -1.219 | 2.47E-02 |
| PA3327 | probable non-ribosomal peptide synthetase | NA | -1.090 | 1.16E-02 |
| PA3330 | probable short chain dehydrogenase | NA | -1.800 | 2.42E-02 |
| PA3354 | hypothetical protein | NA | -1.176 | 3.51E-03 |
| PA3406 | *hasD* | transport protein HasD | -1.511 | 4.95E-02 |
| PA3414 | hypothetical protein | NA | -2.001 | 3.99e-12 |
| PA3419 | hypothetical protein | NA | -1.223 | 1.43E-02 |
| PA3444 | conserved hypothetical protein | conserved hypothetical protein | -1.465 | 1.22E-02 |
| PA3445 | conserved hypothetical protein | conserved hypothetical protein | -1.809 | 1.75E-02 |
| PA3446 | conserved hypothetical protein | conserved hypothetical protein | -2.888 | 1.41E-04 |
| PA3449 | conserved hypothetical protein | conserved hypothetical protein | -2.316 | 6.65E-04 |
| PA3450 | *lsfA* | NA | -1.978 | 3.78E-03 |
| PA3470 | NUDIX hydrolase | NA | -1.102 | 3.36E-02 |
| PA3496 | hypothetical protein | NA | -1.478 | 5.69E-03 |
| PA3521 | *opmE* | NA | -1.371 | 1.90E-02 |
| PA3552 | *arnB* | ArnB | -3.426 | 4.19E-02 |

| | | | | |
|---|---|---|---|---|
| PA3553 | *arnC* | ArnC | -2.964 | 7.42E-04 |
| PA3554 | *arnA* | ArnA | -2.263 | 2.38E-02 |
| PA3555 | *arnD* | ArnD | -2.376 | 2.87E-02 |
| PA3556 | *arnT* | inner membrane L-Ara4N transferase ArnT | -1.700 | 1.79E-02 |
| PA3566 | conserved hypothetical protein | NA | -1.900 | 2.49e-05 |
| PA3567 | probable oxidoreductase | NA | -1.584 | 7.83E-04 |
| PA3575 | hypothetical protein | NA | -1.634 | 2.21E-02 |
| PA3619 | hypothetical protein | NA | -1.252 | 1.36E-03 |
| PA3669 | hypothetical protein | NA | -1.356 | 3.12E-02 |
| PA3672 | probable ATP-binding component of ABC transporter | NA | -1.836 | 4.90E-02 |
| PA3677 | *mexJ* | NA | -1.563 | 3.49e-05 |
| PA3711 | probable transcriptional regulator | NA | -1.516 | 2.07E-02 |
| PA3712 | hypothetical protein | NA | -2.258 | 4.29E-04 |
| PA3758 | probable N-acetylglucosamine-6-phosphate deacetylase | probable N-acetylglucosamine-6-phosphate deacetylase | -1.319 | 3.37E-02 |
| PA3759 | probable aminotransferase | probable aminotransferase | -1.213 | 3.91E-02 |
| PA3775 | hypothetical protein | NA | -1.610 | 2.96E-02 |
| PA3883 | probable short-chain dehydrogenase | NA | -1.128 | 4.19E-02 |
| PA3928 | hypothetical protein | NA | -1.260 | 2.52E-02 |
| PA3931 | putative methionine-binding protein | putative methionine-binding protein | -2.338 | 2.74E-03 |
| PA3932 | probable transcriptional regulator | NA | -1.060 | 2.87E-02 |
| PA3938 | probable periplasmic taurine-binding protein precursor | probable periplasmic taurine-binding protein precursor | -1.555 | 1.16E-02 |
| PA3959 | hypothetical protein | NA | -1.223 | 2.25E-03 |
| PA3964 | hypothetical protein | NA | -1.908 | 7.65E-04 |
| PA3973 | probable transcriptional regulator | NA | -1.451 | 1.04E-02 |
| PA3976 | *thiE* | thiamin-phosphate pyrophosphorylase | -1.541 | 3.02e-09 |
| PA4033 | *mucE* | NA | 1.048 | 1.28E-02 |
| PA4059 | hypothetical protein | NA | -1.522 | 4.38E-02 |
| PA4128 | conserved hypothetical protein | conserved hypothetical protein | -2.167 | 8.36E-03 |
| PA4131 | probable iron-sulfur protein | NA | -1.851 | 1.76e-15 |

| | | | | |
|---|---|---|---|---|
| PA4193 | probable permease of ABC transporter | NA | -1.254 | 2.16E-02 |
| PA4202 | *nmoA* | nitronate monooxygenase NmoA | -1.443 | 5.17E-03 |
| PA4203 | *nmoR* | NA | -1.665 | 3.89E-04 |
| PA4205 | *mexG* | NA | -1.460 | 1.27E-03 |
| PA4206 | *mexH* | probable Resistance-Nodulation-Cell Division (RND) efflux membrane fusion protein precursor | -1.621 | 7.83E-04 |
| PA4208 | *opmD* | NA | -1.883 | 2.43E-04 |
| PA4219 | *ampO* | NA | -4.553 | 2.06E-02 |
| PA4354 | conserved hypothetical protein | NA | 2.372 | 1.81e-08 |
| PA4355 | *pyeM* | NA | 2.227 | 8.66e-15 |
| PA4356 | *xenB* | NA | 3.326 | 1.48e-15 |
| PA4358 | *feoB* | NA | 1.613 | 4.6e-09 |
| PA4371 | hypothetical protein | NA | -1.553 | 1.43E-04 |
| PA4429 | probable cytochrome c1 precursor | probable cytochrome c1 precursor | 1.066 | 2.69E-02 |
| PA4525 | *pilA* | type 4 fimbrial precursor PilA | 1.346 | 3.03E-02 |
| PA4554 | *pilY1* | NA | 1.020 | 4.36E-02 |
| PA4555 | *pilY2* | NA | 1.452 | 6.95E-03 |
| PA4622 | probable major facilitator superfamily (MFS) transporter | NA | 2.086 | 5.51e-15 |
| PA4623 | hypothetical protein | NA | 4.981 | 5.15e-50 |
| PA4661 | *pagL* | NA | 1.794 | 8.41e-08 |
| PA4689 | hypothetical protein | NA | 1.429 | 2.65e-10 |
| PA4714 | conserved hypothetical protein | NA | -1.852 | 7.82E-03 |
| PA4763 | *recN* | NA | -1.155 | 4.24E-02 |
| PA4776 | *pmrA* | PmrA: two-component regulator system response regulator PmrA | -1.044 | 2.97E-02 |
| PA4816 | hypothetical protein | NA | -1.798 | 1.39E-02 |
| PA4818 | conserved hypothetical protein | NA | -1.579 | 1.77E-02 |
| PA4824 | hypothetical protein | NA | -1.771 | 1.52E-02 |
| PA4871 | hypothetical protein | NA | -1.803 | 1.30E-03 |
| PA4881 | hypothetical protein | NA | 3.805 | 1.09e-20 |
| PA4882 | hypothetical protein | NA | 1.467 | 4.22E-04 |
| PA4889 | probable oxidoreductase | NA | -1.475 | 4.66E-02 |
| PA4980 | probable enoyl-CoA hydratase/isomerase | probable enoyl-CoA hydratase/isomerase | -1.364 | 1.45E-02 |

| | | | | |
|---|---|---|---|---|
| PA4985 | Uncharacterized protein | Uncharacterized protein | -1.092 | 1.56E-02 |
| PA4988 | *waaA* | 3-deoxy-D-manno-octulosonic-acid (KDO) transferase | -1.489 | 5.73e-05 |
| PA5024 | conserved hypothetical protein | NA | -1.779 | 5.27E-03 |
| PA5135 | conserved hypothetical protein | NA | -1.213 | 6.95E-03 |
| PA5145 | hypothetical protein | NA | -1.252 | 3.66e-05 |
| PA5157 | probable transcriptional regulator | NA | 1.674 | 2.07E-02 |
| PA5230 | probable permease of ABC transporter | NA | 1.293 | 1.95E-02 |
| PA5431 | probable transcriptional regulator | NA | -1.025 | 4.36E-02 |
| PA5528 | hypothetical protein | NA | -1.785 | 4.39E-02 |
